# Exploration of BAY 11-7082 as a novel antibiotic

**DOI:** 10.1101/2021.09.28.462244

**Authors:** Victoria E. Coles, Patrick Darveau, Xiong Zhang, Hanjeong Harvey, Brandyn D. Henriksbo, Angela Yang, Jonathan D. Schertzer, Jakob Magolan, Lori L. Burrows

## Abstract

Exposure of the Gram-negative pathogen *Pseudomonas aeruginosa* to sub-inhibitory concentrations of antibiotics increases formation of biofilms. We exploited this phenotype to identify molecules with potential antimicrobial activity in a biofilm-based high-throughput screen. The anti-inflammatory compound BAY 11-7082 induced dose-dependent biofilm stimulation, indicative of antibacterial activity. We confirmed that BAY 11-7082 inhibits growth of *P. aeruginosa* and other priority pathogens, including methicillin-resistant *Staphylococcus aureus* (MRSA). We synthesized 27 structural analogues, including a series based on the related scaffold 3-(phenylsulfonyl)-2-pyrazinecarbonitrile (PSPC), 10 of which displayed increased anti-*Staphylococcal* activity. Because the parent molecule inhibits the NLR Family Pyrin Domain Containing 3 (NLRP3) inflammasome, we measured the ability of select analogues to reduce IL-1β production in mammalian macrophages, identifying minor differences in the structure-activity relationship for the anti-inflammatory and antibacterial properties of this scaffold. Although we could evolve stably resistant MRSA mutants with cross resistance to BAY 11-7082 and PSPC, their lack of shared mutations suggested that the two molecules could have multiple targets. Finally, we showed that BAY 11-7082 and its analogues potentiate the activity of penicillin G against MRSA, suggesting that this scaffold may serve as an interesting starting point for the development of antibiotic adjuvants.

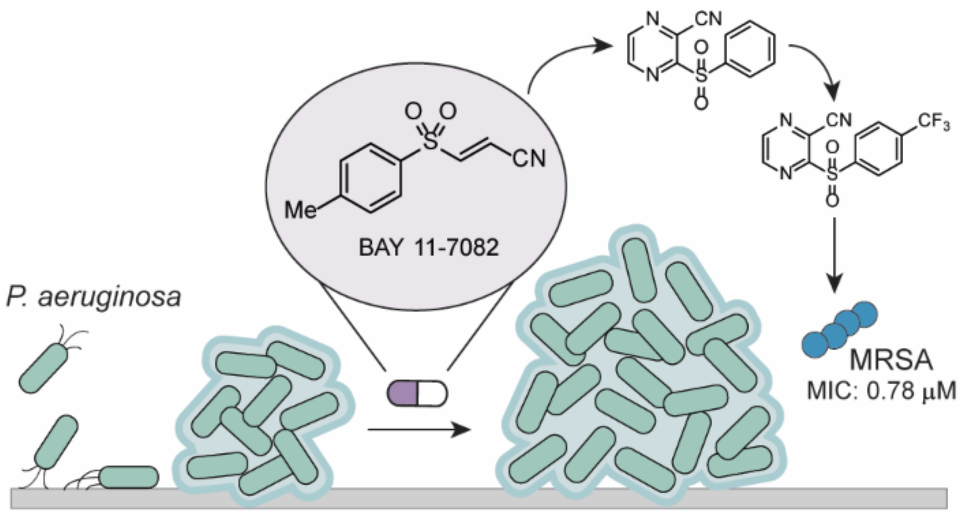

## Introduction

The overuse and misuse of antibiotics has selected for resistant bacteria, increasing the prevalence of drug-resistant infections. As resistance renders standard treatment options less effective, there is a need for the development of novel pharmaceuticals to treat those infections, particularly those caused by the ESKAPE (*Enterococcus faecium, Staphylococcus aureus, Klebsiella pneumoniae, Acinetobacter baumannii, Pseudomonas aeruginosa*, and *Enterobacter* species) pathogens.^1,2^ Despite this urgent need, approval of new antimicrobials by the Food and Drug Administration (FDA) has decreased in recent decades, with only nine new antibiotics approved between 1998-2002.^3^ The rate of approval increased slightly within the last ten years, with eight new antibiotics approved between 2010-2015; however, of these, only three are active against ESKAPE pathogens and only one has a novel mechanism of action, with the rest belonging to established antibiotic classes.^4^

Development of novel antimicrobials with potent activity against the ESKAPE pathogens – including methicillin-resistant *S. aureus* (MRSA) – is crucial. Since strains of MRSA are extensively β-lactam resistant, treatment options are scarce. They include vancomycin, daptomycin, linezolid, tetracyclines, and rifampin, though rifampin is given only in combination therapies due to high rates of resistance development.^5^ MRSA isolates resistant to vancomycin, daptomycin, and linezolid have already been observed in the clinic,^6–8^ and high rates of adverse reactions to linezolid further complicate treatment.^9^ There is therefore a need for safe, orally available anti-MRSA compounds with novel antimicrobial mechanisms to which resistance has not yet been observed.

Traditional methods for identifying new antibiotics include screening libraries of compounds for growth inhibition, but frequently fail to produce promising hits.^10^ Heavily-exploited historic techniques including screening soil extracts for natural products rarely uncover novel molecules, while high-throughput screening using *in vitro* targets often fails to produce hits with significant activity in whole-cell assays.^11,12^ One way to increase the likelihood of uncovering hit compounds is by using bioactive molecules, including drugs approved for non-antibiotic indications, since hit rates are significantly higher.^13^ To increase our chances of identifying new antimicrobials, we developed a biofilm-based assay for high-throughput screening of previously-approved drugs and compounds with known bioactivity. Formation of *P. aeruginosa* biofilms – surface-associated communities of bacteria encased in a self-produced extracellular matrix – acts as a sensitive proxy for detecting antimicrobial activity, since it is stimulated in a dose-dependent manner by sub-inhibitory concentrations of antibiotics.^14^ The biofilm stimulation phenotype allows us to screen compounds for growth inhibition at a low concentration of 10 µM, while alerting us to compounds that may have antibiotic activity above this concentration. Compounds with weaker antibiotic activity can then be improved through medicinal chemistry optimization, ensuring we miss fewer potential hits.

Using a biofilm stimulation screen, we identified the anti-inflammatory BAY 11-7082 as having antibacterial activity against multiple species, including MRSA. BAY 11-7082 is a synthetic anti-inflammatory that inhibits the nuclear factor-κB (NF-кB) pathway and the NLR Family Pyrin Domain Containing 3 (NLRP3)/caspase-1 inflammasome.^15,16^ It is proposed to act by blocking IкB-α phosphorylation but may have alternate cellular targets, and its exact mechanism of action remains unknown.^15,17^ BAY 11-7082 has also been proposed to act on mammalian protein tyrosine phosphatases (PTPs), which are involved in numerous cellular processes.^18^ We re-synthesized BAY 11-7082 and began a medicinal chemistry effort to optimize its anti-*Staphylococcal* activity and determine the compound’s structure-activity relationship (SAR). We ultimately transitioned to a related scaffold, 3-(phenylsulfonyl)-2-pyrazinecarbonitrile (PSPC) and identified 10 analogues with improved anti-MRSA activity, validating the utility of biofilm stimulation as a tool for novel antibiotic discovery.

## Results

### BAY 11-7082 stimulates *P. aeruginosa* biofilm formation and has potent anti-MRSA activity

Using a previously described biofilm assay,^19,20^ we screened a library of 3,921 FDA approved or biologically active compounds. Compounds were screened in duplicate at 10 µM against *P. aeruginosa* PAO1 in a dilute media comprising 10% lysogeny broth (LB), 90% phosphate buffered saline (PBS) (hereafter 10% LB). The screen yielded 60 compounds that inhibited *P. aeruginosa* growth to less than 50% of the vehicle (DMSO) control, 8 that inhibited biofilm formation to less than 50% of the control, and 60 that stimulated biofilm formation above 200% of the control.^20^ One of these biofilm stimulators was the anti-inflammatory BAY 11-7082 **1** (Figure 1A), which we further explored due to its previously unreported antibacterial activity. Using a dose-response assay (Figure 1B), we confirmed that BAY 11-7082 stimulated *P. aeruginosa* biofilm formation at low concentrations and inhibited growth at higher concentrations.

**Figure 1.**
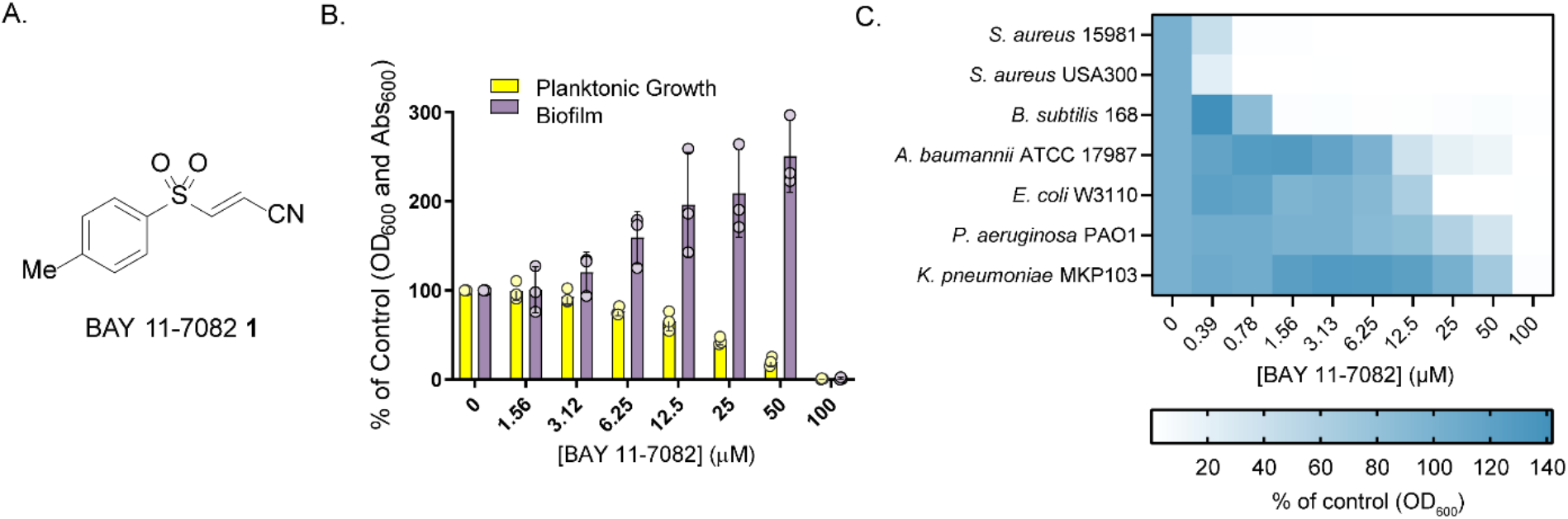
BAY 11-7082 **1** stimulates biofilm formation and has broad-spectrum antibacterial activity. (A) Structure of BAY 11-7082. (B) BAY 11-7082 stimulates biofilm formation (absorbance of crystal violet at 600 nm as a percent of the DMSO control) and decreases planktonic growth (optical density at 600 nm as a percent of the DMSO control) of *P. aeruginosa* PAO1 in 10% LB. (C) BAY 11-7082 inhibits planktonic growth (optical density at 600 nm as a percent of the DMSO control) of seven bacterial species in 10% LB. Heatmap represents an average of three independent experiments performed in triplicate.

To explore its antibacterial potency and spectrum of activity, we determined the minimal inhibitory concentration (MIC) in 10% LB for *P. aeruginosa* plus six other model and priority pathogens: *A. baumannii, Bacillus subtilis, Escherichia coli, K. pneumoniae*, and two strains of MRSA (*S. aureus* 15981 and USA300) (Figure 1C). While BAY 11-7082 displayed modest activity against *P. aeruginosa* (MIC: 100 µM, 20.73 µg/mL) and other Gram-negatives, it was particularly potent against Gram-positives, including MRSA (MIC: 0.78 µM, 0.16 µg/mL), leading us to focus our efforts on developing BAY 11-7082 as an anti-*Staphylococcal* agent. We also determined the MIC in two standard nutrient-rich growth media, LB and Mueller-Hinton Broth (MHB) (Supplementary Figure 1). Based on the MIC in MHB, we selected *S. aureus* 15981 over *S. aureus* USA300 as the strain of choice for comparing synthetic analogues, since the lower baseline MIC would allow us to see a greater range of activities.

To test whether BAY 11-7082 has activity against drug-resistant strains, we exploited its modest activity against *E. coli* to test it against the Antibiotic Resistance Platform (ARP),^21^ a collection of hyperpermeable efflux deficient *E. coli* BW25113 Δ*bamB*Δ*tolC* strains carrying genes that confer resistance to a wide variety of antibiotics. A subset of the ARP was pinned onto Mueller Hinton Agar (MHA) containing BAY 11-7082 at 25, 50, or 100 µM (Supplementary Figure 2). None of the *E. coli* hyperpermeable strains were able to grow at concentrations as low as 25 µM (5.18 µg/mL), indicating that BAY 11-7082 is an effective antibiotic even in the presence of common resistance elements.

### Synthesis and antibacterial activity of analogues

To improve potency and better understand the molecule’s SAR, we synthesized a series of 15 informative structural analogues (Figure 2) and evaluated them for anti-MRSA activity. Analogues **5**-**18** were synthesized following General Procedure A (Supplementary Information), where a sodium sulfinate **2** was combined with 2,3-dibromopropane nitrile, carboxamide or methyl ester **3** to allow for installation of the alkene and the nitrile, or an alternate electron withdrawing group, on the right side of the molecule. Analogues were designed with modifications to the left and/or right sides, including those with alternate heterocycles or alternate substituents on the benzene ring on the left side (**5-8, 10-16, 18, 19**) and those with alternate electron withdrawing groups on the right side (**7, 9-11, 13-18**). Included in this group of analogues is BAY 11-7085 **6** which, like BAY 11-7082, has reported anti-inflammatory activity.^15^ To determine if the alkene is essential for antimicrobial activity, which would suggest the mechanism requires irreversible binding to a target at that position, we synthesized **19** through hydrogenation over palladium on carbon of the potent analogue **5**.

**Figure 2.**
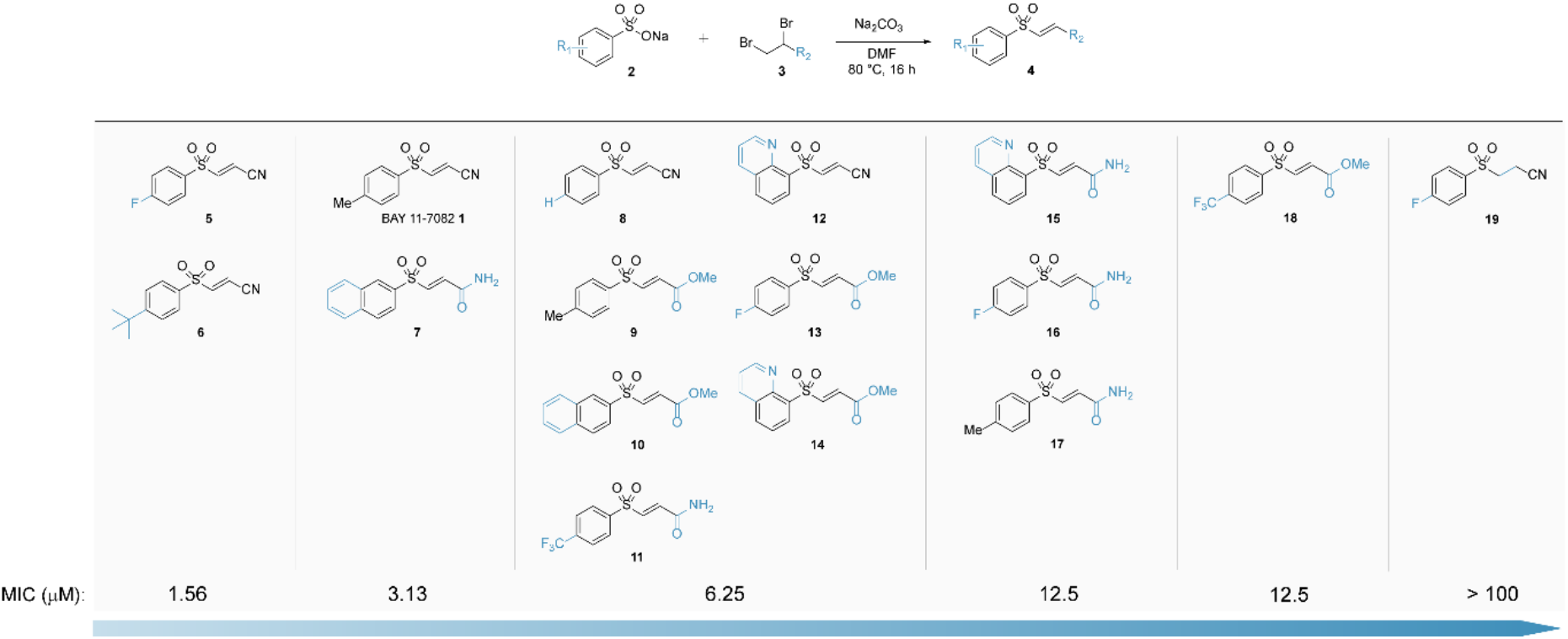
Synthesis (top) and antibacterial activity (bottom) of BAY 11-7082 **1** and its analogues **5**-**19**. Structures are arranged in order of increasing MIC values, reported in µM for *S. aureus* 15981 grown in MHB.

Only **5** and **6** had improved potency against MRSA compared to BAY 11-7082. Moving from a methyl group to the fluorine **5** or tert-butyl **6** increased potency. Similarly, **8**, which lacked substituents on the benzene ring, was less potent than BAY 11-7082. With the surprising exception of **7**, analogues with substituents larger than a tert-butyl group, including the bicyclic heterocycles, had reduced potency, potentially due to steric hinderance at the molecular target(s). All other analogues with a non-nitrile group on the right side of the molecule were less active, leading us to conclude the nitrile is necessary for potent antimicrobial activity. As expected, **19** lacked antimicrobial activity up to the highest concentration tested of 100 µM.

To further explore the molecule’s SAR, we transitioned to a similar scaffold with an additional ring, offering new sites for diversification. We selected 3-(phenylsulfonyl)-2-pyrazinecarbonitrile (PSPC) **20** due to its structural similarities to BAY 11-7082 and its reported *in vivo* activity against MRSA in a *Caenorhabditis elegans* nematode infection model.^22^ This scaffold also had the advantage of lacking the αβ-unsaturated sulfone in BAY 11-7082 that could lead to toxicity. We first confirmed that PSPC stimulated *P. aeruginosa* biofilm formation and had a similar spectrum of antibacterial activity, including potent activity against MRSA (Supplementary Figure 3), then synthesized the analogues **24**-**34** following General Procedure B (Supplementary Information), where a chlorocarbonitrile pyrazine or pyridine **21** was combined with a sulfonyl chloride **22** (Figure 3). PSPC differs from BAY 11-7082 by the addition of a pyrazine between the sulfone and the nitrile. While all analogues retain the nitrile group, some have a pyridine instead of a pyrazine (**33, 34**). Capitalizing on our success with the fluorine analogue **5**, we designed analogues with halogen substituents at the para position of the benzene (**24, 25, 28, 33, 34**), as well as **30** with a meta fluorine, three analogues with two fluorine substituents at differing positions (**27, 31, 32**), and **26** with a para trifluoromethyl group. We also explored a nitrile group (**29**) as an example of a non-halogen para electron withdrawing substituent.

**Figure 3.**
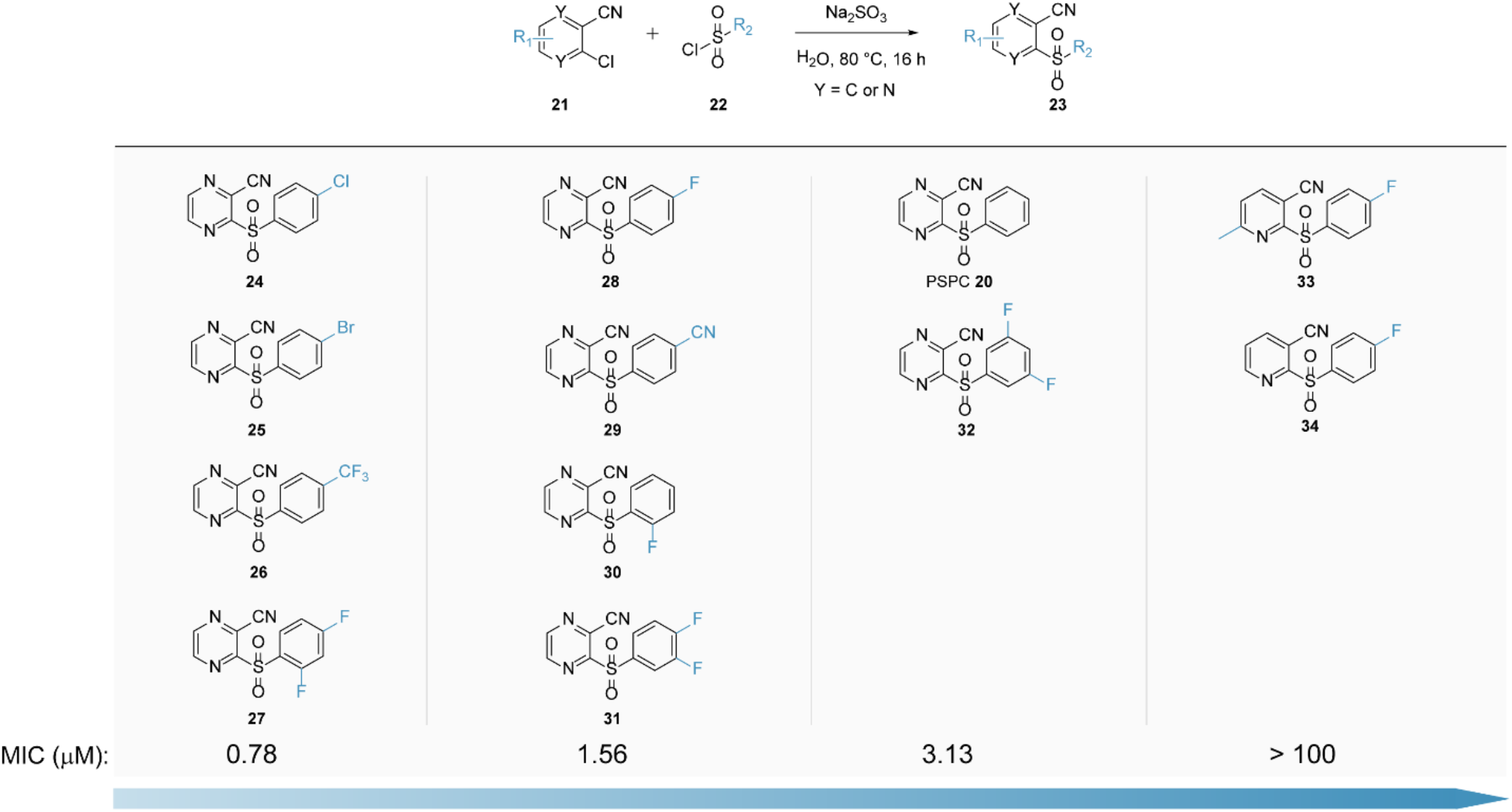
Synthesis (top) and antibacterial activity (bottom) of PSPC **20** and its analogues **24**-**34**. Structures are arranged in order of increasing MIC values reported in µM for *S. aureus* 15981 grown in MHB.

Analogues **24-31** all exhibited increased anti-MRSA activity relative to BAY 11-7082 and their parent compound PSPC, with MICs as low as 0.78 µM. Similar to our results with the first series of analogues, the addition of a halogen or other relatively small electron withdrawing substituent on the benzene ring increased potency, though the number or position of these groups had little impact. Pyridine analogues **33** and **34** lack antibacterial activity up to the highest concentration tested of 100 µM, indicating that while the pyrazine itself is not necessary for antibacterial activity – as in BAY 11-7082 and analogues **5**-**14** – the addition of a more basic heterocycle abrogates activity.

### Evolving mutants resistant to BAY 11-7082 and PSPC

To understand BAY 11-7082 and PSPC’s mechanism of action (MOA) and determine whether the two scaffolds shared a common target, we attempted to evolve resistant mutants of *S. aureus* 15981. While we initially isolated colonies grown on MHA using supra-MIC (6X and 8X MIC) concentrations of BAY 11-7082, those colonies did not grow when re-streaked on fresh MHA containing the compounds. We were similarly unable to develop spontaneous resistance in *E. coli* or *A. baumannii*. We had more success developing resistant mutants by serially passaging *S. aureus* 15981 cultures in liquid media containing ~0.5X, ~1X MIC, or ~2X MIC concentrations of BAY 11-7082 or PSPC for 14 days. After purification of resistant mutants on MHA, six colonies were selected for further evaluation, all of which displayed a 4-fold increase in MIC compared to the parent strain (Figure 4A). Mutants selected using BAY 11-7082 displayed cross-resistance to PSPC and vice versa, suggesting that these two compounds act via a conserved antibacterial mechanism (Figure 4B). To test whether resistance was retained in the absence of selective pressure, the six mutants were serially passaged in MHB for five days and the compounds’ MIC was recorded each day. Over five days there was no decrease in MIC, indicating resistance is stably inherited (Figure 4C). In growth experiments, all six mutants showed shorter lag times and increased final optical density over 16 hours compared to the unpassaged parent strain (p <0.0001) (Figure 4D), perhaps due to a general increase in fitness resulting from passage. To ensure these mutants did not have elevated antibiotic tolerance in general, we determined the MICs to tetracycline and piperacillin, and both were unchanged in the mutants compared to the parent strain (Supplementary Figure 4). To identify the genetic changes responsible for these mutants’ resistance to BAY 11-7082 and PSPC, we purified and sequenced genomic DNA from the parent strain *S. aureus* 15981 and the six mutants. Unexpectedly, the mutants lacked shared genetic changes that could be associated with the acquisition of resistance (Supplementary Table 1), forcing us to pursue other strategies to better understand the antibacterial MOA.

**Figure 4.**
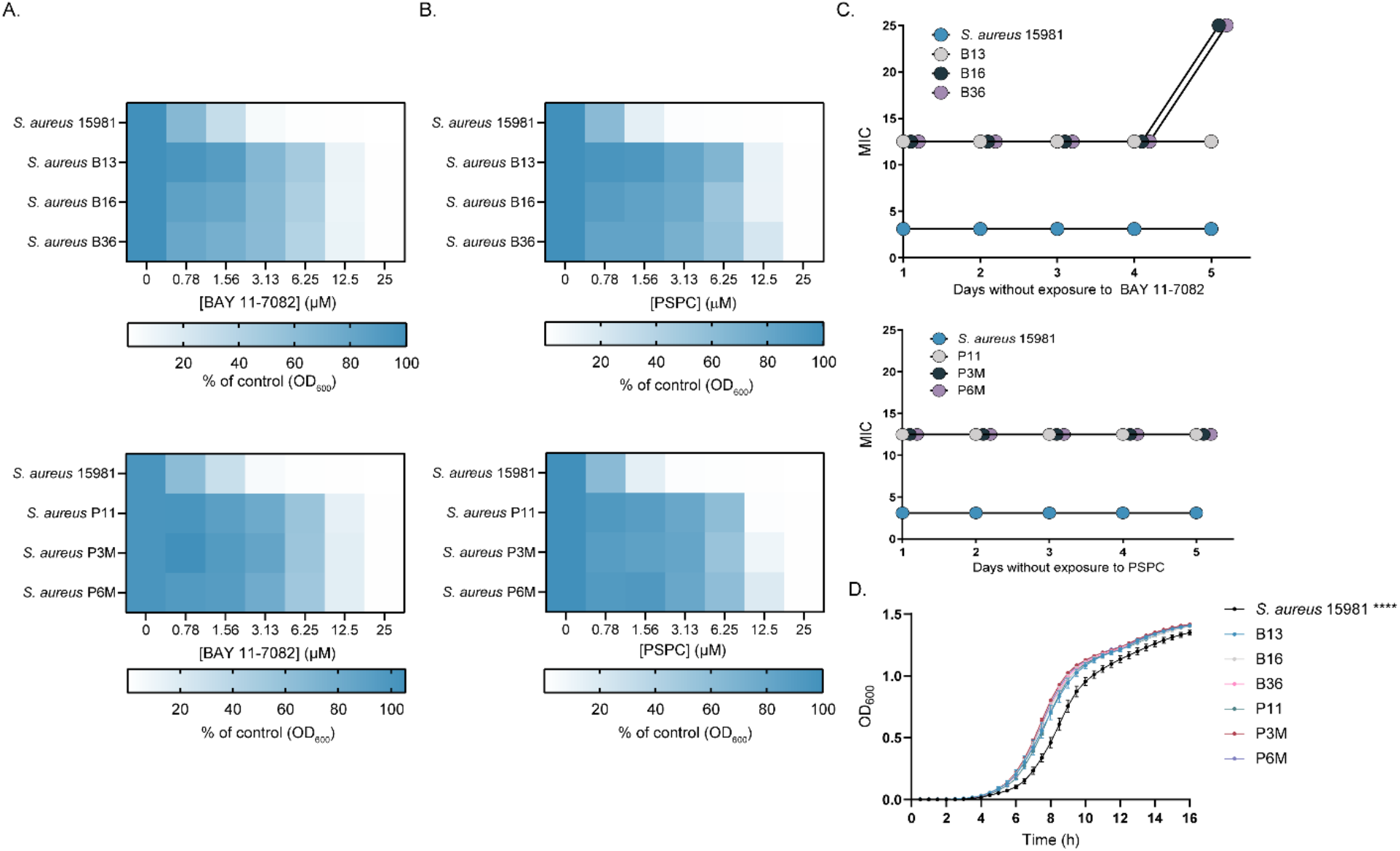
Mutants resistant to BAY 11-7082 and PSPC. (A) Planktonic growth (optical density at 600 nm as a percent of the DMSO control) of *S. aureus* 15981 and BAY 11-7082-(top) or PSPC-resistant (bottom) mutants grown with increasing concentrations of the compound used for selection. (B) Planktonic growth (optical density at 600 nm as a percent of the DMSO control) of *S. aureus* 15981 and BAY 11-7082-(top) or PSPC-resistant (bottom) mutants grown with increasing concentrations of the other compound. (C) MIC values for *S. aureus* 15981 or resistant mutants determined each day for five days during serial passage in the absence of compound. (D) Planktonic growth (optical density at 600 nm) of *S. aureus* 15981 or resistant mutants over 16 h. **** p<0.0001.

### Exploring inhibition of TAT secretion as a potential mechanism

BAY 11-7082 was previously identified as a hit in a screen for inhibitors of the *P. aeruginosa* twin arginine translocase (TAT) system, a Sec-independent protein export system responsible for transporting folded proteins across the cytoplasmic membrane.^23^ While TAT is not essential,^24,25^ we were curious whether BAY 11-7082 would inhibit the *S. aureus* TAT system and if this would provide insight into its antibacterial mechanism. The TAT system is broadly conserved in prokaryotes but its protein composition varies between species, with the *P. aeruginosa* TAT system comprising three genes (*tatA, tatB*, and *tatC*) while the *S. aureus* system comprises two (*tatA* and *tatC*), with *tatA* and *tatC* essential for protein translocation in both species.^26,27^ We determined the potency of BAY 11-7082 and PSPC against *tatC* transposon mutants of *P. aeruginosa* PA14, from the PA14NR Set transposon mutant library,^28^ and *S. aureus* USA300, from the Nebraska transposon mutant library,^29^ and saw a 2-fold increase in sensitivity to BAY 11-7082 and PSPC relative to wild type only for *P. aeruginosa* (Figure 5). This was consistent with the fact that we saw no genetic changes in TAT-related genes in the BAY 11-7082 and PSPC-resistant mutants of MRSA and allowed us to rule out involvement of the TAT system or proteins exported by the system in the compounds’ MOA.

**Figure 5.**
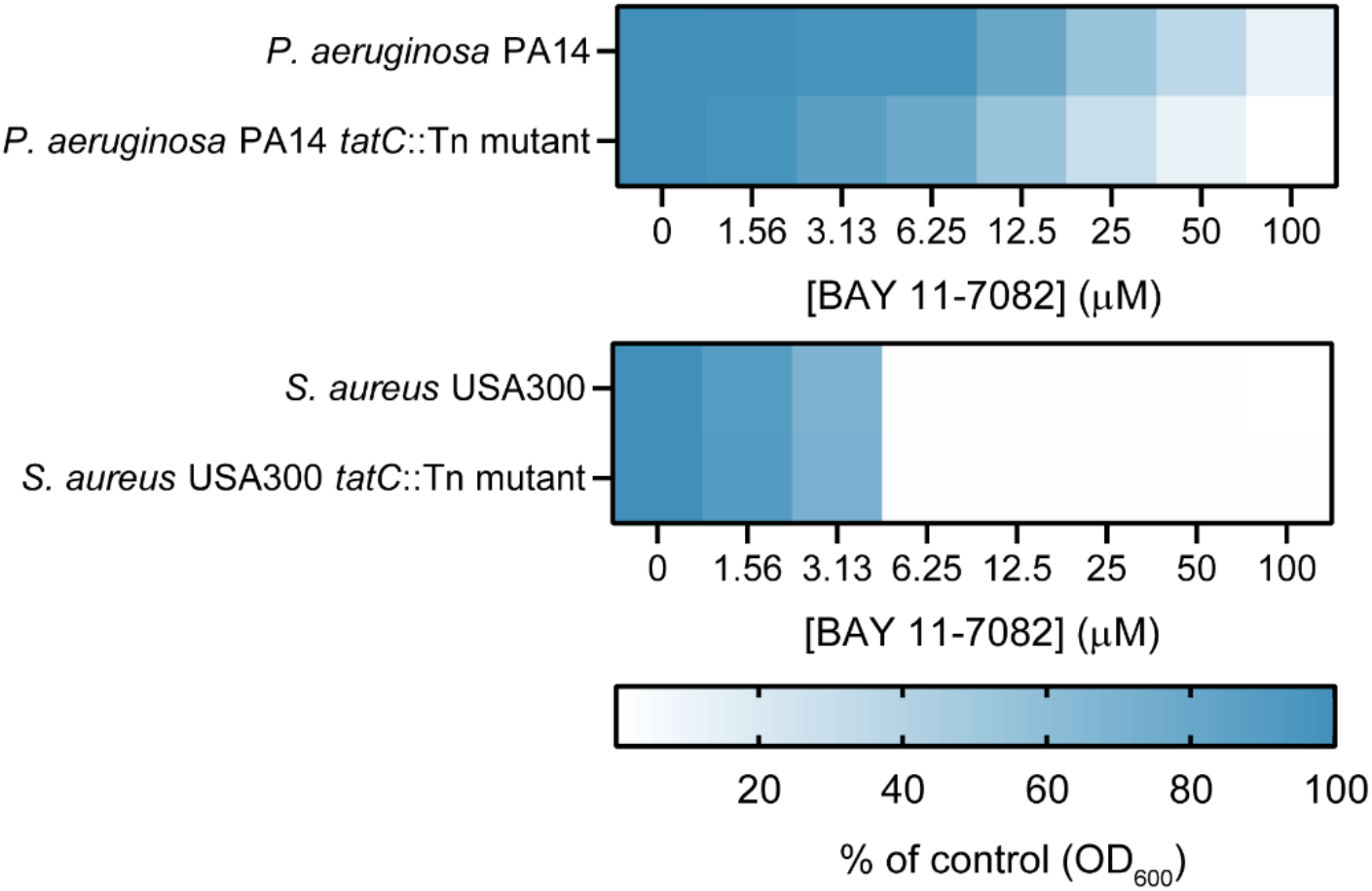
Planktonic growth (optical density at 600 nm as a percent of the DMSO control) of the parent strain *P. aeruginosa* PA14 and a *tatC* transposon mutant grown in 10% LB (top) or the parent strain *S. aureus* USA300 and a *tatC* transposon mutant grown in MHB (bottom) in the presence of increasing concentrations of BAY 11-7082.

### Insight into BAY 11-7082’s uptake and effect on the proton motive force

Given the challenges of isolating BAY 11-7082- and PSPC-resistant mutants and our inability to isolate mutants with greater than a 4-fold increase in MIC, plus the lack of consistent genetic changes in the mutants isolated, we wondered if BAY 11-7082 and PSPC had a more general MOA, such as perturbation of the proton motive force (PMF). The PMF has two components, a proton gradient (ΔpH) and an electric potential (Δψ), which the cell coordinates to control the electrochemical gradient across the membrane.^30^ To test this idea, we determined the MIC of BAY 11-7082 and PSPC against *S. aureus* 15981 in pH adjusted MHB (Figure 6A). Both BAY 11-7082 and PSPC displayed reduced potency in more basic conditions and increased potency in more acidic conditions, similar to tetracycline, which is taken up via the ΔpH component,^31^ suggesting that BAY 11-7082 and PSPC may also experience ΔpH-dependent uptake across the cytoplasmic membrane.

**Figure 6.**
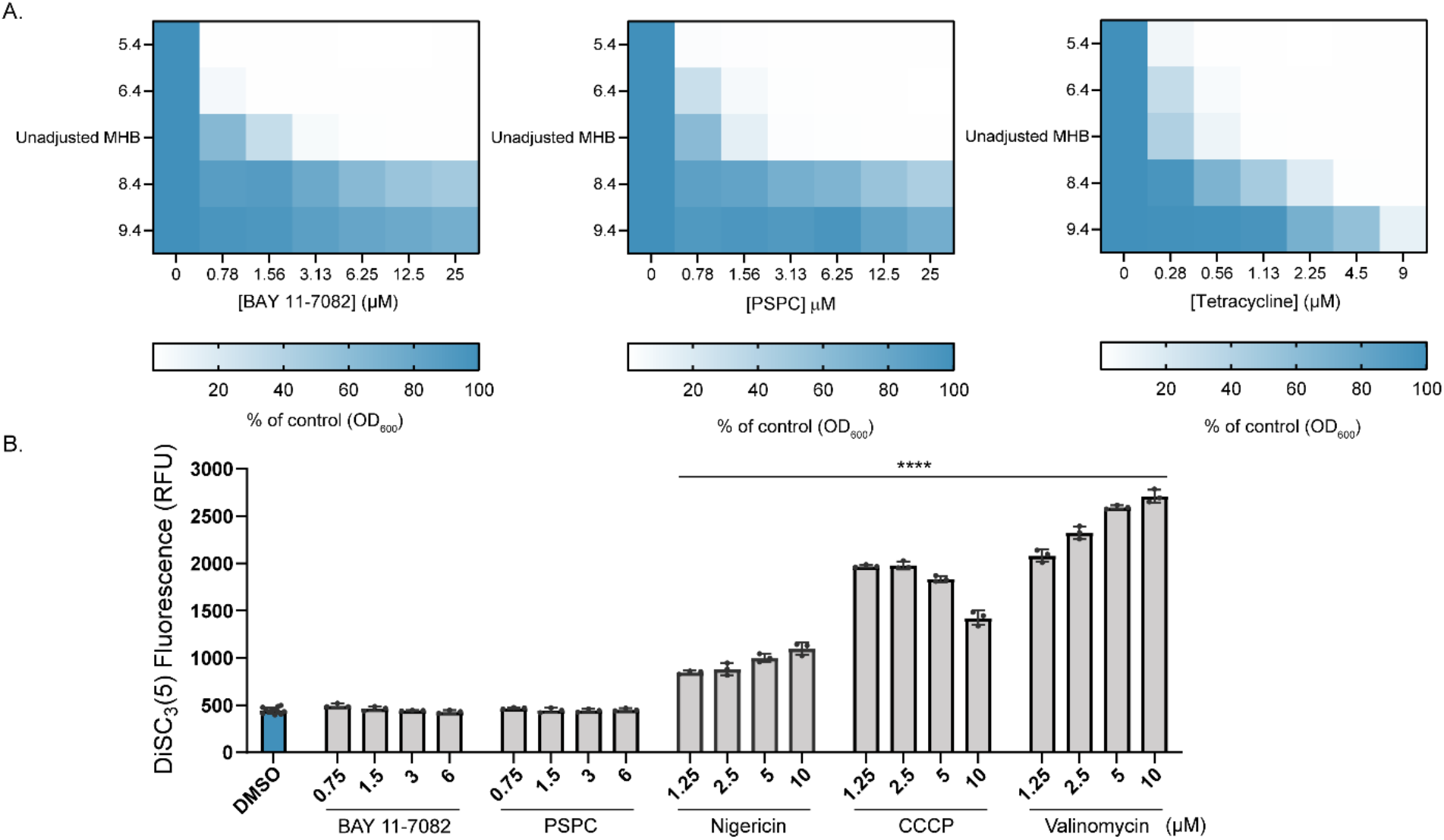
(A) Planktonic growth (optical density at 600 nm as a percent of the DMSO control) of *S. aureus* 15981 grown in pH adjusted MHB with increasing concentrations of BAY 11-7082 (left), PSPC (center), or tetracycline (right). Y-axis indicates pH of the media. (B) DiSC_3_(5) fluorescence in relative fluorescence units of *S. aureus* USA300 in the presence of the vehicle DMSO, BAY 11-7082, PSPC, or the membrane active controls nigericin, CCCP, or valinomycin. **** p<0.0001.

To determine if BAY 11-7082 and PSPC affected the PMF, we used 3,3’-dipropylthiadicarbocyanine iodide (DiSC_3_(5)), a fluorescent dye sensitive to PMF perturbations (Figure 6B). The dye is taken up and transported to the cytoplasm via the Δψ component, where it accumulates and self-quenches fluorescence. If Δψ is inhibited, dye does not accumulate, resulting in increased fluorescence, while if ΔpH is inhibited, the cell compensates by increasing flux through Δψ, resulting in decreased fluorescence. Valinomycin, nigericin, and carbonyl cyanide m-chlorophenyl hydrazine (CCCP) all dissipate the PMF and were used as positive controls. Unlike the controls (p<0.0001), neither BAY 11-7082 nor PSPC significantly altered DiSC_3_(5) fluorescence (p>0.9999), confirming that while their uptake is PMF-dependent, they themselves do not destabilize the PMF.

### BAY 11-7082 synergizes with multiple classes of antibiotics

We next looked at whether BAY 11-7082 and its analogues could act as adjuvants to potentiate the activity of other antibiotics. We conducted checkerboard experiments using the clinically-relevant MRSA strain *S. aureus* USA300. It should be noted that the MIC for BAY 11-7082 and its analogues were ~4-fold higher for *S. aureus* USA300 than the MICs for *S. aureus* 15981 reported in Schemes 2 and 3. Using the fractional inhibitory concentration index (FICI), where values < 0.5 represent synergy and values > 4 represent antagonism,^32^ we determined that BAY 11-7082 synergized with the β-lactam antibiotics penicillin G and piperacillin, the RNA polymerase inhibitor rifampin, and the translation inhibitor thiostrepton (Figure 7A). BAY 11-7082 did not affect the activity of antibiotics dependent on the PMF for uptake, including tetracycline (ΔpH-dependent) and gentamicin (Δψ-dependent) (Figure 7A), consistent with our observations that it does not alter the PMF. BAY 11-7082 did not synergize with the glycopeptide cell wall synthesis inhibitor vancomycin or the β-lactam methicillin, to which *S. aureus* USA300 is intrinsically resistant (Figure 7A). Given that we saw the strongest synergy between BAY 11-7082 and penicillin G, we tested some of our more potent analogues in combination with that β-lactam. We selected the BAY 11-7082 para-fluorine analogue **5** (MIC for *S. aureus* USA300: 12.5 µM) and the para-bromine and para-trifluoromethyl PSPC analogues **25** and **26** (MICs for *S. aureus* USA300: 6.25 µM); both potentiated penicillin G activity to the same degree as BAY 11-7082 (FICI: 0.19) (Figure 7B).

**Figure 7.**
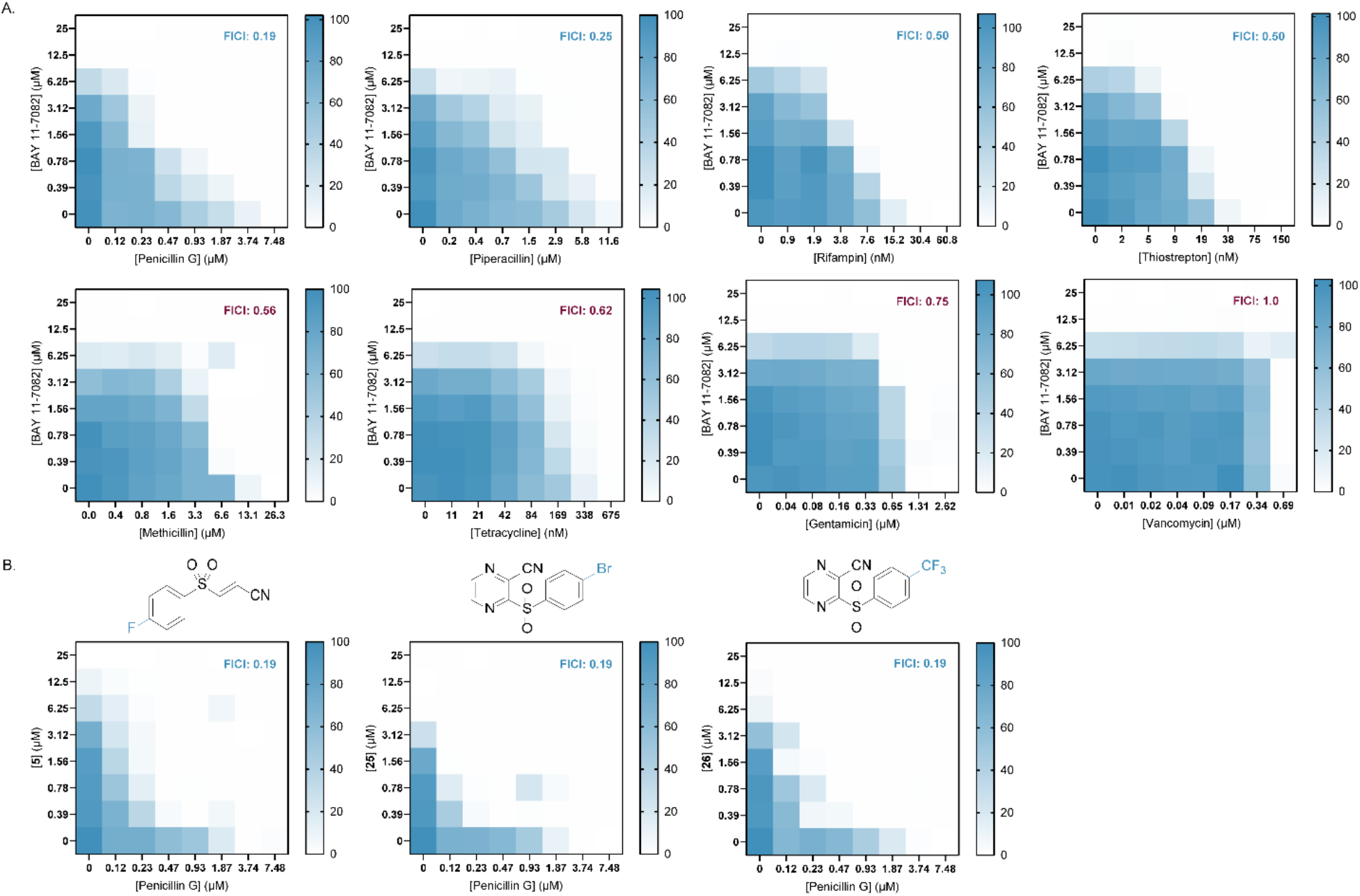
Checkerboards of *S. aureus* USA300 planktonic growth (optical density at 600 nm as a percent of the DMSO control) grown in MHB with increasing concentrations of BAY 11-7082 (A) or its analogues **5, 25**, or **26** (B) with antibiotics of different classes. Synergy (FICI<0.5) is represented in blue text while indifference (FICI>0.5) is represented in red. Checkerboards represent an average of at least three independent experiments.

### Anti-inflammatory properties of BAY 11-7082 and its analogues

Since the original BAY 11-7082 compound has anti-inflammatory activity, we tested whether alterations to the scaffold would allow us to separate its anti-inflammatory and antimicrobial properties. We treated mouse bone marrow-derived macrophages with *E. coli* O111:B4 lipopolysaccharide (LPS) and nigericin and measured IL-1β production, a key effector cytokine produced by priming and activating the NLRP3/caspase-1 inflammasome, in the presence or absence of BAY 11-7082 and four analogues (**5, 12, 17**, and **19**) (Figure 8). As expected, treatment with BAY 11-7082 at 1 and 2.5 µM significantly reduced (p<0.0001) IL-1β production compared to the no-treatment control. The potent antimicrobial **5** caused a similar reduction in IL-1β production at both 1 (p=0.0057) and 2.5 (p<0.0001) µM, as did the weaker antimicrobials **12** (p<0.0001) and **17** (p<0.0001). As expected, there was no significant difference in IL-1β production between treatment with **19** – which lacks antimicrobial activity due to the reduction of the alkene – at 2.5 µM versus the no-treatment control (p=0.5600), while treatment with 1 µM resulted in a significant increase in IL-1β production (p=0.0061), supporting our notion that covalent binding is required for the scaffold’s anti-inflammatory activity. Surprisingly, treatment with **17**, which has an MIC 4-fold higher than that of BAY 11-7082, resulted in a greater, though not significant (p>0.9999), reduction in IL-1β production at 2.5 µM compared to BAY 11-7082, indicating the terminal nitrile is not required for anti-inflammatory potency. This result suggests that while it may not be possible to develop a potent antimicrobial that does not also target the inflammasome, there may be room to develop a potent anti-inflammatory compound with a reduced impact on the microbiome.

**Figure 8.**
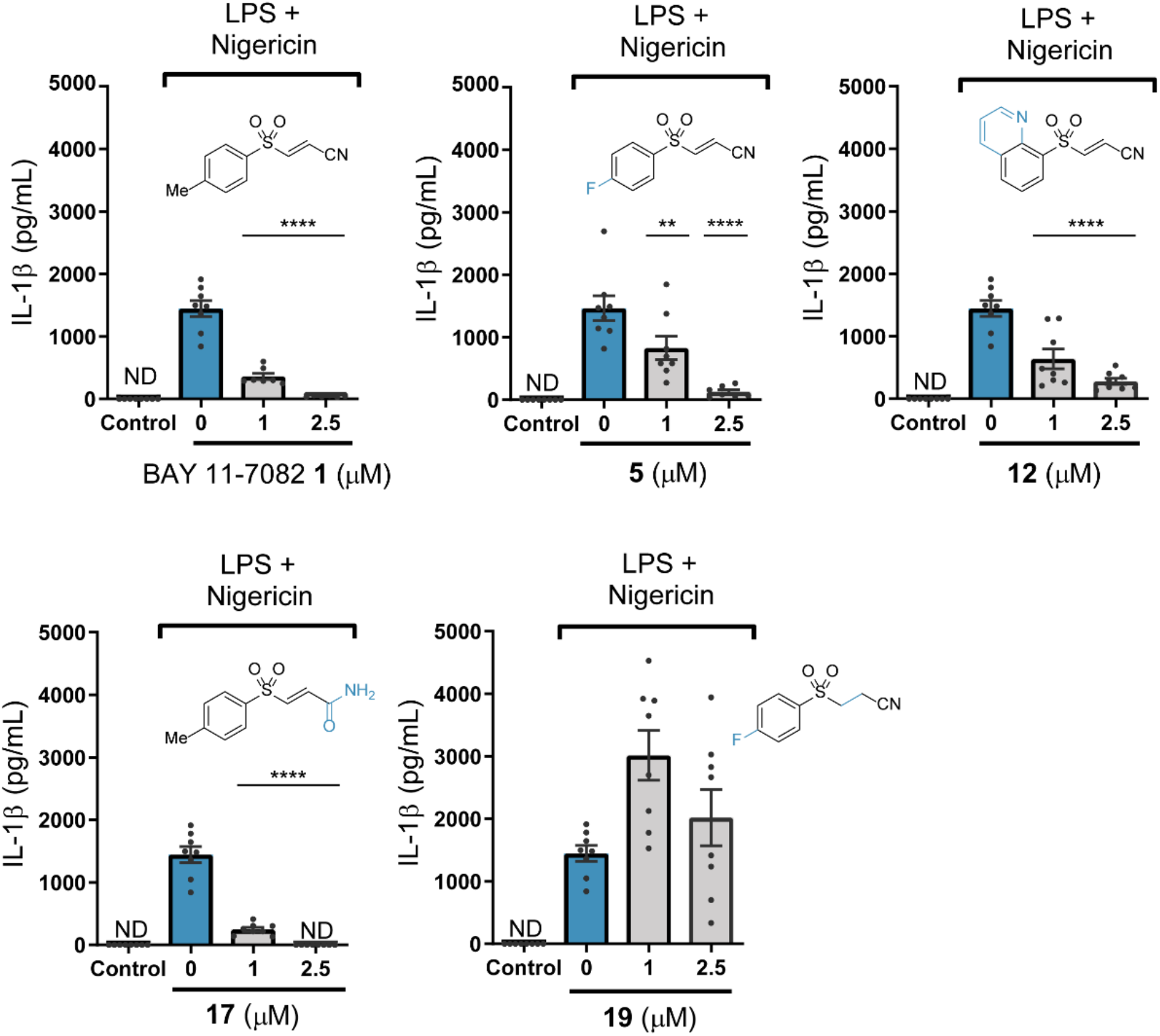
Production of IL-1β by bone marrow-derived macrophages stimulated with 200 ng/mL LPS and 10 µM nigericin in the presence or absence of BAY 11-7082, or the analogue **5, 12, 17**, or **19** at 1 or 2.5 µM. Control cells did not receive stimulation with LPS and nigericin. ND (not detected) indicates IL-1β levels for that condition were below the level of detection based on a standard curve generated for IL-1β concentration. ** p<0.01, **** p<0.0001

## Discussion

Biofilm stimulation occurs in response to exposure to sub-inhibitory concentrations of antibiotics, independent of their MOA,^14,33^ making it a useful tool with advantages over traditional growth assays. The biofilm assay does not rely on growth inhibition at an arbitrary screening concentration, alerting us to hits with potential MICs above the test concentration. Our previous work showed that *P. aeruginosa* biofilm production is a useful proxy for antimicrobial activity, both for narrow spectrum anti-*Pseudomonal* compounds and for broad range molecules like the antibiotic thiostrepton.^20^ Here we confirmed that *P. aeruginosa* biofilm stimulation by the anti-inflammatory BAY 11-7082 at 10 µM signalled its antimicrobial activity against a variety of species, including MRSA.

BAY 11-7085 **6**, another anti-inflammatory and structural analogue of BAY 11-7082, was recently identified as an antimicrobial with activity against *S. aureus* and *Candida albicans*.^34^ Similar to our findings, BAY 11-7085 lacked activity against *P. aeruginosa* and *K. pneumoniae* strains up to 276.7 µM, and Escobar *et al*. were unable to isolate BAY 11-7085-resistant mutants of *S. aureus*. Using vancomycin-resistant *S. aureus* VRS1, they found an MIC of 16 µM in MHB and showed that BAY 11-7085 inhibited cell surface attachment and biofilm formation and eradicated established biofilms in brain heart infusion broth media with 0.1% glucose. Interestingly, they did not report biofilm stimulation in response to sub-MIC concentrations of BAY 11-7085.

When initially describing BAY 11-7082 and BAY 11-7085’s anti-inflammatory activity, Pierce *et al*. suggested they act via preventing IκB-α phosphorylation initiated by cytokines like tumor necrosis factor-α (TNF-α), resulting in lower NF-κB activation and nuclear translocation, where it regulates transcription of κB sequences with various roles in immunity, cell survival, and apoptosis. Pierce *et al*. proposed BAY 11-7082 and BAY 11-7085 could inhibit a protein tyrosine kinase upstream of IκB-α but did not propose a specific target. Subsequent work showed that BAY 11-7082 binds and irreversibly inhibits mammalian PTPs through nucleophilic attack by a cysteine in the enzyme’s active site,^18^ suggesting that BAY 11-7082’s antibacterial target could be a PTP. In bacteria, classical PTPs play roles in virulence and host immune evasion, while low-molecular-weight PTPs are primarily involved in polysaccharide transport.^35^*S. aureus* contains two low molecular weight PTPs, PtpA and PtpB, both of unknown function.^36^ We also considered the TAT system as a target, as BAY 11-7082 was previously identified as a *P. aeruginosa* TAT inhibitor, where it was also suggested to bind a nucleophilic cysteine.^23^ However, another study reported that BAY 11-7082 failed to inhibit protein export via the *E. coli* TAT system.^37^ Further, loss of the TAT system in these species is not lethal,^24,25^ incongruent with the antimicrobial activity of BAY 11-7082, nor was its activity different in a *S. aureus* TAT mutant (Figure 3). Finally, we saw no insertions, deletions, or single nucleotide polymorphisms in PTP or TAT-related genes in mutants with increased resistance to BAY 11-7082 or PSPC (Supplementary Table 1), consistent with our conclusion that BAY 11-7082’s antibacterial MOA is unrelated to either of these systems. Given our inability to identify a molecular target for BAY 11-7082, we suggest that its mechanism is likely complex. While this may be due in part to the potential promiscuity conferred by BAY 11-7082’s αβ-unsaturated sulfonyl, this moiety is absent in PSPC, suggesting it does not play a role in the compound’s MOA.

The αβ-unsaturated center, characteristic of highly-reactive Michael acceptors, makes BAY 11-7082 an unlikely drug candidate. Michael acceptors have traditionally been considered problematic in drug development due to concerns that the compounds could irreversibly bind off-target proteins. However, views on the suitability of Michael acceptors and other covalent modifiers as drugs are changing – particularly in the anticancer space – with several recent examples of αβ-unsaturated carbonyls under development.^38–41^ Such compounds work best when the αβ-unsaturated carbonyl binds a known and biologically-uncommon target, reducing the likelihood of irreversible covalent off-target binding. Despite our efforts, we were unable to identify a specific molecular target for BAY 11-7082, suggesting the compound enacts its antibacterial effect via multiple mechanisms. We were also unable to identify an analogue of BAY 11-7082 lacking anti-inflammatory activity but retaining antibiotic potency. While dual antibiotic-anti-inflammatory compounds are an interesting concept and, indeed, many antibiotics have been shown to affect the inflammasome,^42^ this raises concern for potential side effects. Research from the Taunton lab aimed at developing reversible cysteine residue inhibitors showed that positioning a nitrile group on olefins increases thiol reactivity reversibility, as does positioning electron withdrawing groups at the α or β position of acrylonitriles.^43,44^ It is important to note that with the exploration of PSPC and the development of analogues **24-31**, we showed that the αβ-unsaturated center is not required for potent anti-MRSA activity.

Beyond its anti-inflammatory activity, BAY 11-7082 has anticancer, anti-diabetes, and neuroprotective properties.^45–48^ It is thus noteworthy that its potent activity against Gram-positive species could prove problematic should the compound be developed for one of these indications, as it has potential to impact the microbiome. This is not uncommon among drugs with human targets. Indeed, a recent study found 24% of all human-directed drugs tested inhibited growth of at least one bacterial species common to the microbiome.^49^ We found that analogue **17**, a weak anti-*Staphylococcal*, maintains strong anti-inflammatory activity, suggesting it could be possible to separate the activities through careful design. Due to its potential off-target effects, perhaps a better use of this antibacterial scaffold would be as a lead for adjuvants used in combination with β-lactams to treat MRSA infections. Although BAY 11-7082 did not re-sensitize MRSA strain USA300 to methicillin, at 1.56 µM it reduced the MIC of penicillin G from 3.74 µM to 0.23 µM – a 16-fold reduction. This concentration of BAY 11-7082 is well below the 10 µM reported IC_50_ for inhibition of IκB-α phosphorylation,^15^ but above the concentration that significantly reduced inflammation in our macrophage model (Figure 6) and the K_i_ for some mammalian PTPs,^18^ suggesting that even at the low concentrations afforded by combination therapy, using this scaffold may result in off-target effects. The BAY 11-7082 analogue **5** and the PSPC analogues **25** and **26** also synergized with penicillin G and can be used at even lower concentrations (0.78 µM for **7** and 0.39 µM for **25** and **26**) than BAY 11-7082 to achieve the same 0.23 µM MIC for penicillin G. Both **25** and **26** have the advantage of lacking the αβ-unsaturated center found in BAY 11-7082 and **5**, making them the most promising candidates for development of antibacterial combination therapies.

## Conclusion

A high-throughput screen using *P. aeruginosa* biofilm stimulation as an indicator of antibacterial activity led to identification of BAY 11-7082 as a broad-spectrum antimicrobial, with activity against multiple priority pathogens, including MRSA. This work confirms biofilm stimulation as a useful screening method for identifying compounds with broad activity. We were able to improve upon the potency of BAY 11-7082 by switching to a new chemical scaffold based on the molecule PSPC, which lacks the αβ-unsaturated center that may be associated with off-target effects and toxicity. Our analogues **25** and **26** potentiate the activity of β-lactams like penicillin G to reduce the concentration needed to inhibit the growth of MRSA. These two compounds therefore represent an interesting starting point for the development of an antibacterial adjuvant for the treatment of MRSA.

## Methods

### Bacterial strains, culture conditions, and chemicals

Bacterial strains used in this study include *P. aeruginosa* PAO1^50^, *S. aureus* 15981,^51^ *B. subtilis* 168,^52^ *E. coli* K-12 W3110^53^, *A. baumannii* ATCC 17987^20^, and *K. pneumoniae* MKP103.^54^ *P. aeruginosa* PA14 and the Δ*tatC* transposon mutant were from the PA14NR Set transposon mutant library^28^ and *S. aureus* USA300 and the Δ*tatC* transposon mutant was from the Nebraska transposon mutant library.^29^ Bacterial cultures were grown in lysogeny broth (LB), 10% LB (10% LB, 90% 1X phosphate buffered saline), or Mueller-Hinton broth (MHB). Solid media was solidified with 1.5% agar. Compounds **1, 7**-**22**, and **26**-**37** were synthesized for this work. One hundred mM stock solutions were made in DMSO and stored at -20°C. Stock solutions of tetracycline (EMD Millipore), piperacillin (Sigma, 40 mg/mL stock solution), penicillin G (Fluka Analytica), rifampin (Sigma, 1 mg/mL stock solution prepared in DMSO), thiostrepton (Sigma, 5 mM stock solution prepared in DMSO), methicillin (Cayman Chemicals), gentamicin (BioShops), and vancomycin (Sigma, stock solution prepared in DMSO) were prepared at 10 mg/mL in sterile de-ionized H_2_O except where noted and stored at -20°C.

### Compound screening

The McMaster Bioactive compound collection was screened using a biofilm modulation assay, as previously described.^20^

### Inhibitory concentration assays

All bacterial strains were inoculated from -80°C glycerol stocks into 5 mL media, and incubated with shaking, 200 rpm, 37°C for 16 h. Cultures were sub-cultured in a 1:500 dilution, grown for 4 h, and diluted in fresh media to an OD_600_ of ~ 0.1, then diluted 1:500. Minimal inhibitory concentration values were determined in 96-well plates (Nunc) using compounds serially diluted in DMSO. Each well had a final volume of 150 µL. Sterile control wells contained 148 µL of corresponding broth and 2 µL DMSO while vehicle control wells contained 148 µL dilute bacterial culture and 2 µL DMSO. Plates were sealed in a container and incubated with shaking, 200 rpm, 37°C, 16 h. The OD_600_ of plates was read using a Multiscan GO (Thermo Fisher Scientific) and used to calculate the inhibitory concentration. All MIC values presented in Scheme 1 and Scheme 2 were conducted with *S. aureus* 15981 grown in MHB and are based on experiments performed in duplicate. All other inhibitory concentration data was conducted in media as specified and is based on at least three independent experiments performed in triplicate. MIC values were defined as < 20% of the vehicle control, where growth is no longer visible.^20^ Where specified, media was pH-adjusted using 6 N sodium hydroxide and 6 M hydrogen chloride.

### Biofilm modulation assays

Biofilm assays were performed as previously described.^20^ Briefly, *P. aeruginosa* PAO1 was inoculated from -80°C glycerol stocks into 5 mL 10% LB, and incubated with shaking, 200 rpm, 37°C for 16 h. Cultures were sub-cultured in a 1:500 dilution, grown for 4 h, and diluted in fresh media to an OD_600_ of ~ 0.1, then diluted 1:500. Plates were set up as for the inhibitory concentration assays. Biofilms were formed on the polystyrene pegs of MicroWell lids (Nunc), which were placed on assay plates prior to 16 h incubation with shaking, 200 rpm, 37°C. Following incubation, lids were removed and planktonic growth (OD_600_) was measured. Lids were immersed in 1X phosphate buffered saline and washed for 10 min, then biofilms adhering to the lids were stained in 0.1% crystal violet for 15 min. Excess crystal violet was removed by immersing the lids in deionized water for 5 min, then repeating this wash step four additional times. Lids dried for 1 h then were transferred to a new 96-well plate containing 200 µL 33.3% acetic acid per well for 5 min to solubilize the crystal violet. The absorbance (Abs_600_) of the 96-well plates was read using a Multiscan GO (Thermo Fisher Scientific). Three independent experiments were performed in triplicate and technical triplicates were averaged. The mean and standard error of the independent biological replicates are shown.

### Liquid serial passage assays

*S. aureus* 15981 was taken from -80°C stocks, inoculated into 5 mL MHB, and grown with shaking, 200 rpm, 37°C for 24 h. One hundred µL culture was inoculated into fresh MHB supplemented with BAY 11-7082 or PSPC at 1 (B13, B16, and P11), 3 (B36, P3M), or 6 (P6M) µM and grown with shaking 200 rpm, 37°C for 24 h. One hundred µL culture was inoculated into fresh MHB supplemented with BAY 11-7082 or PSPC at the same concentration as the previous day. This serial liquid passage continued for 14 days, at which point cultures were purified on MHA with or without compound.

### Genome assembly and identification of SNPs

Genomic DNA was purified using the Wizard Genomic DNA purification kit (Promega) then sequenced via Illumina sequencing. Quality control and trimming of sequencing reads was accomplished using Galaxy through FASTQC for quality control, FASTQC Groomer to reformat reads, Trimmomatic to trim poor quality reads, FASTQC Interlacer to reorder reads, and FASTQC De-Interlacter. Contigs were assembled for the *S. aureus* 15981 genome using SPAdes and genomes were mapped to reference genomes of two closely related species, *S. aureus* USA300 and USA500, using BreSeq.

### Growth Curves

*S. aureus* 15981 and the six BAY 11-7082- or PSPC-resistant mutants (B13, B16, B36, P11, P3M, P6M) were taken from -80°C stocks, inoculated into 2 mL MHB, and grown with shaking, 200 rpm, 37°C for 16 h. Overnight cultures were sub-cultured in a 1:500 dilution, grown for 4 h, and diluted in fresh media to an OD_600_ of ~ 0.1, then diluted 1:500. Three replicates of 150 µL of each sample was added to a 96-well plate and incubated with moderate shaking at 37°C (BioTek Synergy 4 Plate Reader) and the OD_600_ was measured at 30 min intervals for 16 h. Technical replicates were averaged and the experiment was repeated at least three times for each strain. The mean and standard error of the independent biological replicates are shown. P-values were calculating using a one-way ANOVA and Tukey’s multiple comparison test.

### DiSC_3_(5) assays

The DiSC_3_(5) assay was performed in accordance with modifications by Farha *et al*^55^ to a method by Epand *et al*^56^, with some modifications. Briefly, *S. aureus* USA300 was taken from -80°C stock, inoculated into 5 mL MHB, and grown with shaking, 200 rpm, 37°C for 16 h. Cultures were sub-cultured in a 1:500 dilution and grown for 4 h. Cells were pelleted and washed three times in buffer (10 mM potassium phosphate, 5 mM MgSO4, and 250 mM sucrose, pH 7.0), then resuspended in buffer to an OD_600_ of 0.1. DiSC_3_(5) was added at 1 µM and allowed to stabilize at room temperature in the dark for 20 minutes. BAY 11-7082, PSPC, valinomycin, nigericin, and CCCP were diluted in DMSO and added in triplicate to a white, clear-bottom, 96-well plate (Costar, Corning Inc), along with cells. Vehicle control wells contained 2 µL DMSO and 148 µL bacteria suspended in buffer with dye. Fluorescence was measured (BioTek Synergy 4 Plate Reader) at an excitation wavelength of 620 nm and an emission wavelength of 685 nm at the start time, then every 50 s for 10 min and every 5 min for the next 30 min. The mean and standard error of three technical replicates for each condition are plotted at the start time. P-values were calculating using a one-way ANOVA and Tukey’s multiple comparison test.

### Checkerboard assays

Checkerboard assays were set up identically to inhibitory concentration assays, except compounds were added to an 8×8 section of a 96-well plate with increasing concentrations of one compound across the X-axis and increasing concentrations of the other compound across the Y-axis. Plates were incubated with shaking, 200 rpm, 37°C, 16 h. The OD_600_ of plates was read using a Multiscan GO (Thermo Fisher Scientific). At least three biological replicates were averaged for each checkerboard. Synergy was determined using the following formular for fractional inhibitory concentration index (FICI), where values < 0.5 represent synergy and values > 4 represent antagonism^32^:

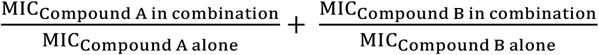

### NLRP3 inflammasome assays

In-house male wild type C57BL/6 J mice (Jackson Laboratory) aged four to twelve weeks were humanely sacrificed to generate bone marrow derived macrophages (BMDMs), as previously described.^57^ Briefly, bone marrow was isolated by flushing the marrow cavity of the femur with sterile saline. Cells recovered from femur flushes were propagated in tissue culture flasks at 1 × 10^6^ cells/mL in L929 differentiation media at 37°C for four days. Additional L929 media was added and cells were transferred to 24-well plates and incubated for another 5-7 days to allow for maturation of BMDMs. L929 media was removed from BMDMs, wells were washed with PBS, and Dulbecco’s Modified Eagle Medium (DMEM) (600 µL) was added. BMDMs were treated with BAY 11-7082 or analogues at a final concentration of 1 and 2.5 µM and *E. coli* O111:B4 LPS diluted in endotoxin-free ultra pure water at a final concentration of 200 ng/mL and incubated at 37°C for 3 h. Nigericin diluted in DMEM was added at a final concentration of 10 µM and cells were incubated for an additional 45 min. Each condition was tested in at least six biological replicates in BMDMs from a single mouse. Two hundred and fifty µL of media was removed from each well and frozen until IL-1β secretion could be measured by ELISA, (DuoSet ELISA, Mouse 1L-1β/IL-1F2, R&D Systems), as previously described.^58^ Media was diluted in a 1:10 ratio in ELISA reagent dilute (1% BSA in PBS) for the LPS and nigericin condition and all test conditions. Media from the control condition, which lacked LPS, nigericin, or the analogues, was diluted in a 1:1 ratio in ELISA reagent dilute. Absorbance at 450 and 570 nm of the 96-well plate following ELISA was read and values at 570 nm were subtracted from values at 450 nm to correct for optical imperfections. IL-1β levels corresponding to the subtracted absorbance values were determined by interpolating from a standard curve generated using IL-1β concentrations ranging from 0 pg/mL to 1000 pg/mL. The mean and standard error for at least six independent biological replicates is shown. P-values were calculating using a one-way ANOVA and Tukey’s multiple comparison test.

## Supporting information

Supplementary Figures for Coles et al.

## Supporting Information

Antibacterial activity of BAY 11-7082 in rich media; activity of BAY 11-7082 against a collection of bacterial strains containing common resistance elements; antibacterial and antibiofilm activity of PSPC; growth of BAY 11-7082- and PSPC-resistant mutants in the presence of tetracycline and piperacillin; synthetic schemes and characterization data (1H NMR, C13 NMR, and HRMS) for compounds 1, **5-20**, and **24-34**.

## Author Contributions

VEC and HH completed antibacterial susceptibility testing. PD, XZ, and VEC completed chemical synthesis, supervised by JM. VEC and BDH conducted the NLRP3 inflammasome assays, supervised by JDS. VEC and AY compiled genetic data related to the resistant mutants. All other experiments were performed by VEC. VEC and LLB wrote the paper, with input from JM and JDS.

## Acknowledgements

We thank Gerry Wright for access to the Antibiotic Resistance Platform, Derek Chan and Kara Tsang for their assistance with BreSeq, and Meghan Fragis for determining exact masses of the compounds. This work was supported by grants from the Natural Sciences and Engineering Research Council (NSERC) RGPIN-2016-06521 and Ontario Research Fund RE07-048 to LLB and a seed grant from the David Braley Centre for Antibiotic Discovery to LLB and JM. The NMR machine was purchased with funding from a Canada Foundation for Innovation grant. VEC was supported by a Summer Studentship from the David Braley Centre for Antibiotic Discovery and a CIHR Canadian Graduate Scholarship – Master’s Award.

## Abbreviations Used

ARP: Antibiotic Resistance Platform;
CCCP: carbonyl cyanide m-chlorophenyl hydrazine;
DiSC_3_(5): 3,3’-dipropylthiadicarbocyanine iodide;
FICI: fractional inhibitory concentration index;
LB: lysogeny broth;
MHA: Mueller-Hinton agar;
MHB: Mueller-Hinton broth;
PMF: proton motive force;
PSPC: 3-(phenylsulfonyl)-2-pyrazinecarbonitrile;
PTP: protein tyrosine phosphatase;
TAT: twin arginine translocase

