## Supplementary Figures for Coles et al. for "Exploration of BAY 11-7082 as a novel antibiotic"

### Exploration of BAY 11-7082 as a novel antimicrobial

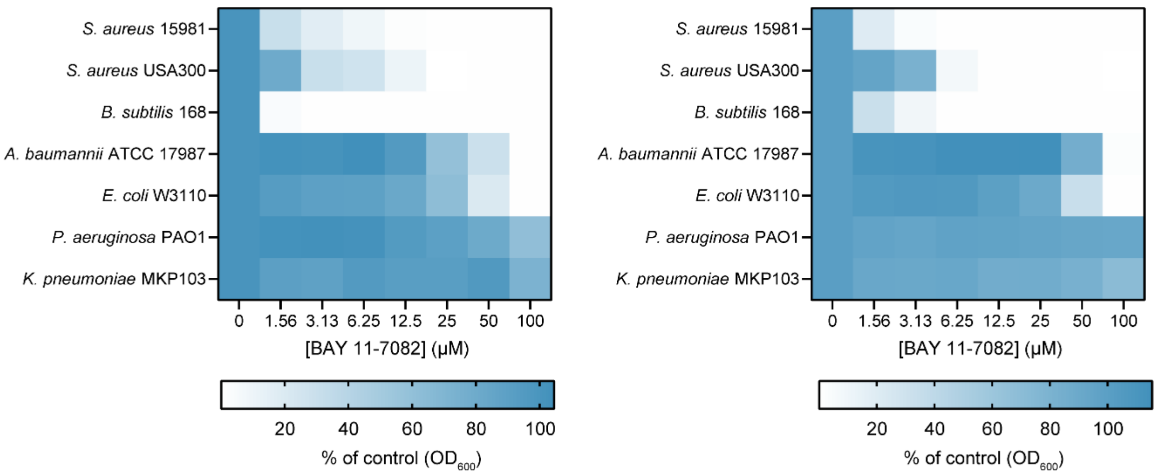


**Supplementary Figure 1.** Planktonic growth (optical density at 600 nm as a percent of the DMSO control) of seven bacterial species grown in LB (left) or MHB (right) with increasing concentrations of BAY 11-7082. The MIC is equal to 20% or less of control growth.

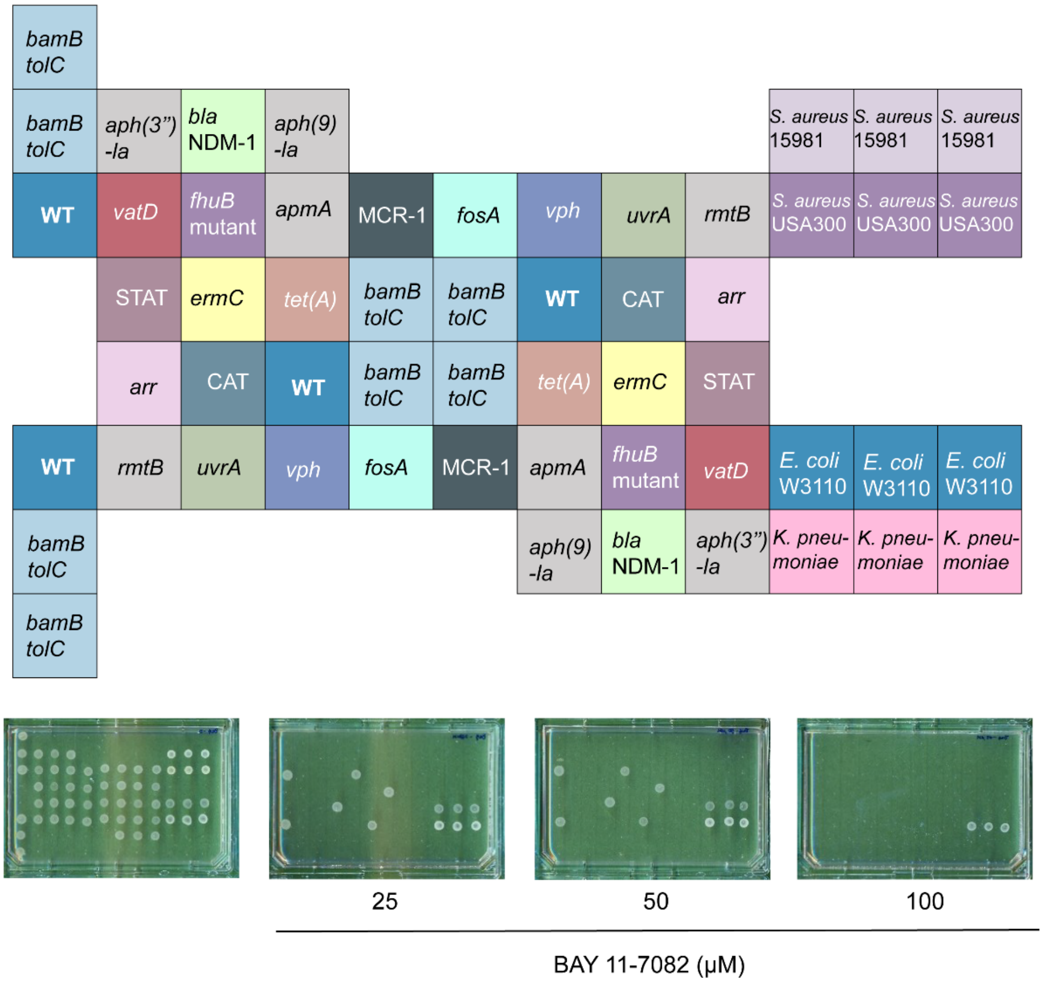


**Supplementary Figure 2.** BAY 11-7082 inhibits the growth of a collection of clinically relevant pathogens and strains from the Antibiotic Resistance Platform.[1] The collection of strains (top), including *S. aureus* 15981, *S. aureus* USA300, *E. coli* W3110, *K. pneumoniae* MKP103, *E. coli* BW25113, *E. coli* BW25113 (WT) *ΔbamBΔtolC* (*bamBtolC*), and *E. coli* BW25113 *ΔbamBΔtolC* strains containing common antibiotic resistance elements, as labeled, were spotted on MHA with or without BAY 11-7082 at concentrations 25, 50, or 100 µM. At 25 and 50 µM only *E. coli* WT and W3110 strains, an *mcr-1* strain (which is in the WT background), and *K. pneumoniae* grew, while at 100 µM, only *K. pneumoniae* grew.

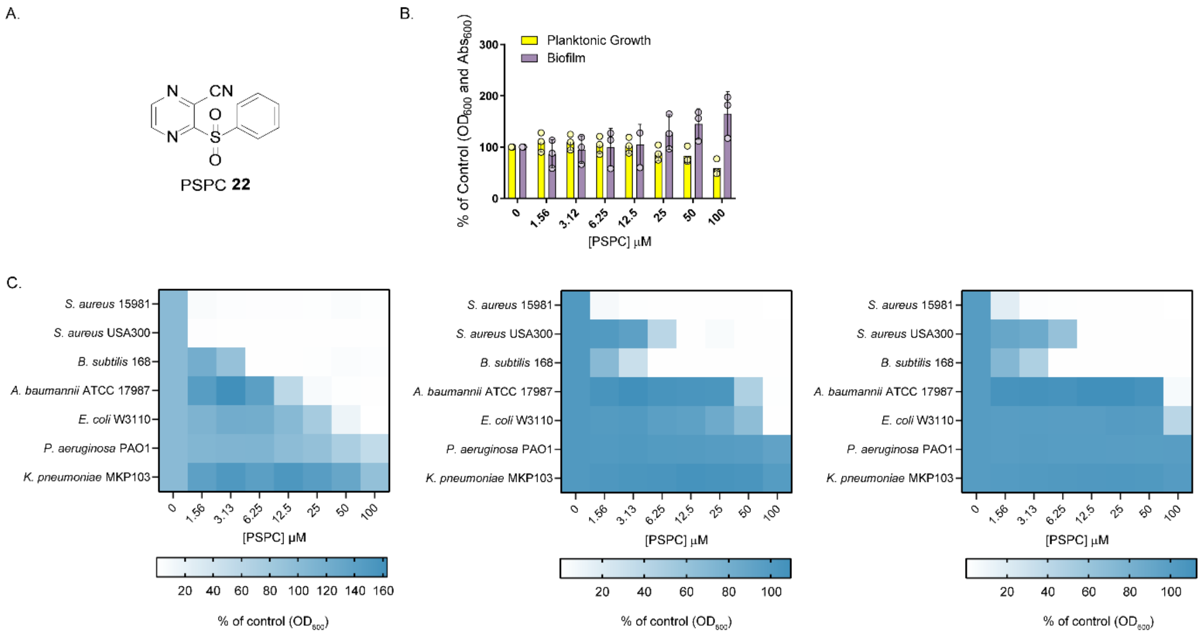


**Supplementary Figure 3.** PSPC (**22**), like BAY 11-7082, stimulates biofilm formation and has potent anti-*Staphylococcal* activity. (A) Structure of PSPC. (B) PSPC stimulates biofilm formation (absorbance of crystal violet at 600 nm as a percent of the DMSO control) and decreases planktonic growth (optical density at 600 nm as a percent of the DMSO control) of *P. aeruginosa* PAO1 in 10% LB. (C) Planktonic growth (optical density at 600 nm as a percent of the DMSO control) of seven bacterial species grown in 10% LB (left), LB (center) or MHB (right) with increasing concentrations of PSPC.

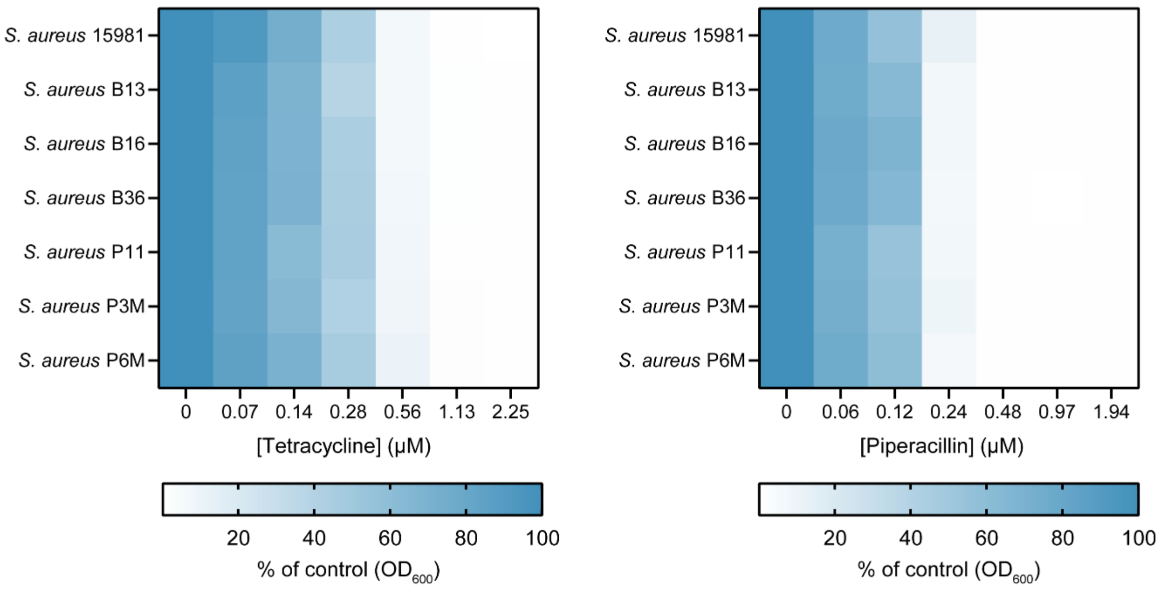


**Supplementary Figure 4.** BAY 11-7082 and PSPC-resistant mutants and the parent strain *S. aureus* 15981 are equally susceptible to tetracycline and piperacillin. Planktonic growth (optical density at 600 nm as a percent of the DMSO control) of *S. aureus* 15981 or a mutant strain grown in MHB with increasing concentrations of tetracycline (left) or piperacillin (right).

**Supplementary Table 1.** Summary of single nucleotide polymorphisms in BAY 11-7082 and PSPC-resistant mutants that differ from the parent *S. aureus* 15981 strain. Available from the authors on request as an Excel file.

**General Procedure A** for the formation of acrylonitriles, acrylamides and methyl acrylates sulfones:

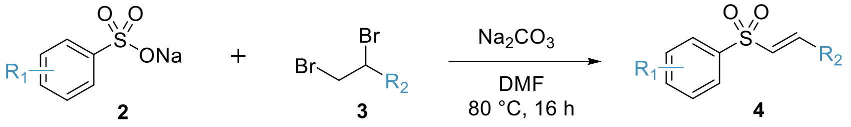


Adapted from a procedure by Gian *et al.*,[2] to a stirring solution of **sodium sulfinate (2)** (2 equiv.) and **$Na_2CO_3$** (2 equiv.) in **DMF** (0.25 M) was added **2,3-dibromopropane nitrile**, **carboxamide** or **methyl ester (3)** (1 equiv.). The resulting mixture was heated up to 80 °C and was stirred overnight. The crude mixture was cooled to room temperature, diluted in water, and extracted with EtOAc (3x). The organic layers were combined and washed with brine (1x), dried over $MgSO_4$, filtered and concentrated under reduced pressure. Purification was executed by normal phase flash chromatography.

**General Procedure B** for the formation of *o*-sulfonylcarbonitrile pyrazines and pyridines:

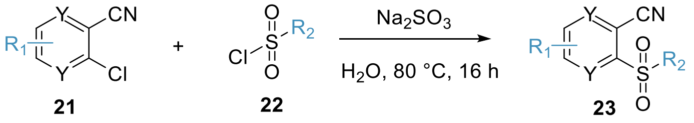


To a stirring solution of ***o*-chlorocarbonitrile pyrazine** or **pyridine (21)** (1 equiv.) and **$Na_2SO_3$** (1.3 equiv.) in **$H_2O$** (0.2 M) was added **sulfonyl chloride (22)** (1.2 equiv.). The resulting mixture was heated to 100 °C and stirred overnight. The crude mixture was cooled to room temperature and extracted with EtOAc (3x). The organic layers were combined and washed with $H_2O$ (1x) and brine (1x), dried over $MgSO_4$, filtered and concentrated under reduced pressure. Purification was executed by reverse phase chromatography, compound-rich fractions were concentrated to dryness, taken into nanopure water, frozen, and lyophilized to yield the desired products.

**Synthesis of (*E*)-3-tosylacrylonitrile (1):**

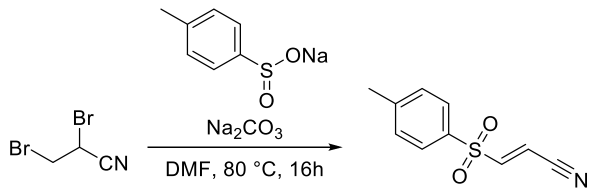


Following the general procedure **A** on 213 mg (1.00 mmol) of **2,3-dibromopropanenitrile**, purification afforded 7.0 mg (3% yield) of **(*E*)-3-tosylacrylonitrile** as a white solid. (Gradient from 0 to 20% EtOAc/hexanes)

$^1$H NMR (700 MHz, Chloroform-*d*) δ 7.78 (d, *J* = 8.1 Hz, 2H), 7.41 (d, *J* = 8.0 Hz, 2H), 7.21 (d, *J* = 15.7 Hz, 1H), 6.51 (d, *J* = 15.7 Hz, 1H), 2.48 (s, 3H).

$^{13}$C NMR (176 MHz, Chloroform-*d*) δ 149.48, 146.68, 134.35, 130.73, 128.73, 113.54, 110.25, 21.93.

HRMS (ESI$^+$): [M+Na]$^+$ calculated for $C_{10}H_9NO_2SNa$ = 230.02462 m/z; found = 230.02529 m/z.

**Synthesis of (*E*)-3-((4-fluorophenyl)sulfonyl)acrylonitrile (5):**

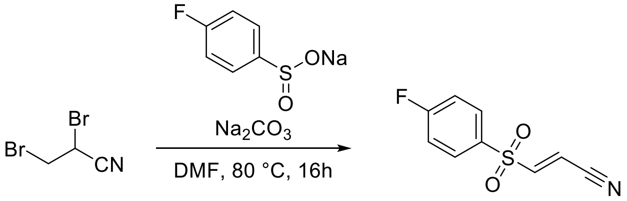


Following the general procedure **A** on 213 mg (1.00 mmol) of **2,3-dibromopropanenitrile**, purification afforded 17.8 mg (9% yield) of **(*E*)-3-((4-fluorophenyl)sulfonyl)acrylonitrile** as a white solid. (Gradient from 0 to 20% EtOAc/hexanes)

$^1$H NMR (700 MHz, Chloroform-*d*) δ 7.95 – 7.92 (m, 2H), 7.31 (t, *J* = 8.9, 8.0 Hz, 2H), 7.21 (d, *J* = 15.6 Hz, 1H), 6.56 (d, *J* = 15.6 Hz, 1H).

$^{13}$C NMR (176 MHz, Chloroform-*d*) δ 167.40, 165.93, 148.81, 131.63, 117.53, 113.19, 110.92.

HRMS (ESI$^+$): [M+Na]$^+$ calculated for $C_9H_6FNO_2SNa$ = 233.99952 m/z; found = m/z.

**Synthesis of (*E*)-3-((4-(*tert*-butyl)phenyl)sulfonyl)acrylonitrile (6):**

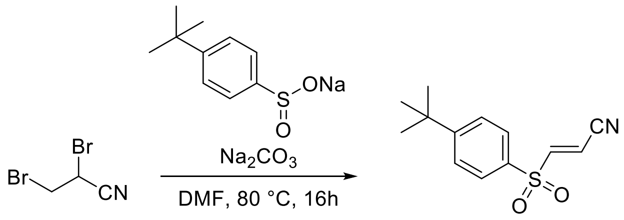


Following the general procedure **A** on 212.9 mg (1.00 mmol) of **2,3-dibromopropanenitrile**, purification afforded 75.6 mg (30% yield) of **(*E*)-3-((4-(*tert*-butyl)phenyl)sulfonyl)acrylonitrile** as a white solid. (Gradient from 0 to 20% EtOAc/hexanes)

$^1$H NMR (700 MHz, DMSO-$d_6$) δ 8.22 (d, J = 15.6 Hz, 1H), 7.86 – 7.82 (m, 2H), 7.76 – 7.72 (m, 2H), 6.89 (d, J = 15.6 Hz, 1H), 1.31 (s, 9H).

$^{13}$C NMR (176 MHz, DMSO-$d_6$) δ 158.82, 149.85, 135.07, 128.51, 127.44, 115.11, 112.22, 35.65, 31.13.

HRMS (ESI$^+$): $[M+H]^+$ calculated for $C_{13}H_{16}NO_2S$ = 250.08958 m/z; found = 250.08857 m/z.

HRMS (ESI$^+$): $[M+Na]^+$ calculated for $C_{13}H_{15}NO_2SNa$ = 272.07152 m/z; found = 272.07121 m/z.

**Synthesis of (*E*)-3-(naphthalen-2-ylsulfonyl)acrylamide (7):**

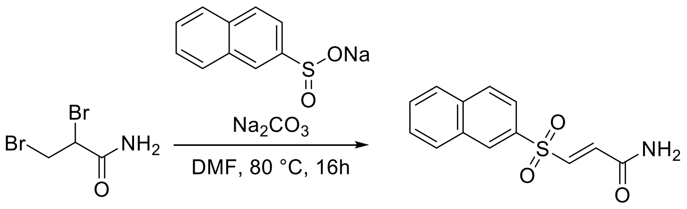


Following the general procedure **A** on 230.9 mg (1.00 mmol) of **2,3-dibromopropanecarboxamide**, purification afforded 256.9 mg (98% yield) of **(*E*)-3-(naphthalen-2-ylsulfonyl)acrylamide** as a white solid. (Gradient from 0 to 20% EtOAc/hexanes)

$^1$H NMR (700 MHz, DMSO-$d_6$) δ 8.64 (d, *J* = 1.9 Hz, 1H), 8.24 (dd, *J* = 8.1, 1.2 Hz, 1H), 8.21 (d, *J* = 8.7 Hz, 1H), 8.12 – 8.08 (m, 1H), 8.04 (s, 1H), 7.89 (dd, *J* = 8.7, 1.9 Hz, 1H), 7.77 (ddd, *J* = 8.2, 6.8, 1.3 Hz, 1H), 7.72 (ddd, *J* = 8.1, 6.8, 1.3 Hz, 1H), 7.69 (s, 1H), 7.52 (d, *J* = 15.0 Hz, 1H), 7.03 (d, *J* = 15.0 Hz, 1H).

$^{13}$C NMR (176 MHz, DMSO-$d_6$) δ 163.08, 139.02, 135.81, 135.36, 134.94, 131.82, 129.94, 129.71, 129.58, 129.49, 127.98, 127.93, 122.42.

HRMS (ESI$^+$): $[M+Na]^+$ calculated for $C_{13}H_{11}NO_3SNa$ = 284.03522 m/z; found = 284.03519 m/z.

**Synthesis of (*E*)-3-(phenylsulfonyl)acrylonitrile (8):**

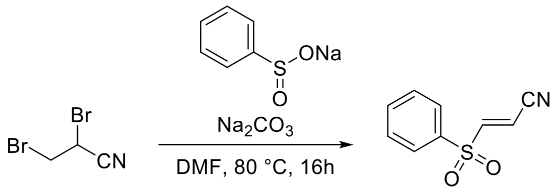


Following the general procedure **A** on 212.9 mg (1.00 mmol) of **2,3-dibromopropanenitrile**, purification afforded 75.6 mg of **(*E*)-3-(phenylsulfonyl)acrylonitrile** as a brown solid. (Gradient from 0 to 20% EtOAc/hexanes)

**Synthesis of methyl (*E*)-3-tosylacrylate (9):**

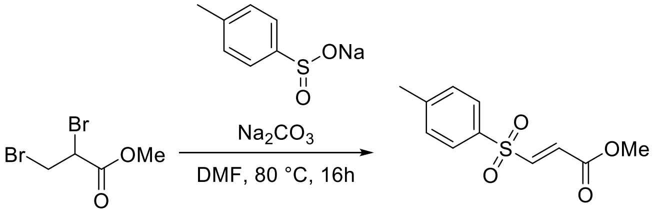


Following the general procedure **A** on 230.9 mg (1.00 mmol) of **methyl 2,3-dibromopropanate**, purification afforded 180.5 mg (80% yield) of **methyl (*E*)-3-tosylacrylate** as a white solid. (Gradient from 0 to 20% EtOAc/hexanes)

$^1$H NMR (700 MHz, DMSO-$d_6$) δ 1H NMR (700 MHz, DMSO-d6) δ 7.85 – 7.82 (m, 2H), 7.78 (d, *J* = 15.2 Hz, 1H), 7.52 – 7.48 (m, 2H), 6.75 (d, *J* = 15.2 Hz, 1H), 3.73 (s, 3H), 2.42 (s, 3H).

$^{13}$C NMR (176 MHz, DMSO-$d_6$) δ 164.16, 145.95, 143.73, 135.76, 130.79, 130.76, 128.57, 53.11, 21.63.

HRMS (ESI$^+$): $[M+H]^+$ calculated for $C_{11}H_{13}O_4S$ = 241.05288 m/z; found = 241.05289 m/z.

HRMS (ESI$^+$): $[M+Na]^+$ calculated for $C_{11}H_{12}O_4SNa$ = 263.03482 m/z; found = 263.03452 m/z.

**Synthesis of methyl (*E*)-3-(naphthalen-2-ylsulfonyl)acrylate (10):**

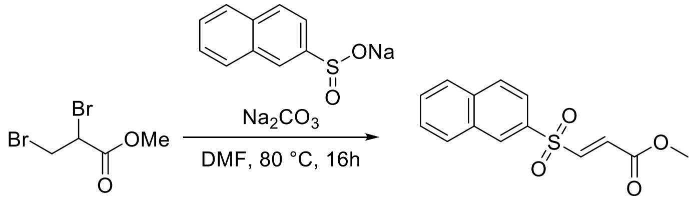


Following the general procedure **A** on 212.8 mg (1.00 mmol) of **methyl 2,3-dibromopropanate**, purification afforded 10.0 mg (4% yield) of **methyl (*E*)-3-(naphthalen-2-ylsulfonyl)acrylate** as a white solid. (Gradient from 0 to 20% EtOAc/hexanes)

$^1$H NMR (700 MHz, Chloroform-*d*) δ 8.56 – 8.51 (m, 1H), 8.05 – 7.99 (m, 2H), 7.95 (d, *J* = 8.2 Hz, 1H), 7.84 (dd, *J* = 8.6, 1.9 Hz, 1H), 7.71 (ddd, *J* = 8.2, 6.9, 1.3 Hz, 1H), 7.67 – 7.64 (m, 1H), 7.40 (d, *J* = 15.1 Hz, 1H), 6.89 (d, *J* = 15.1 Hz, 1H), 3.80 (s, 3H).

$^{13}$C NMR (176 MHz, Chloroform-*d*) δ 163.93, 143.51, 135.61, 135.20, 132.30, 130.52, 130.51, 130.04, 129.80, 129.54, 128.08, 127.99, 122.57, 52.79.

HRMS (ESI$^+$): [M+Na]$^+$ calculated for $C_{14}H_{12}O_4SNa$ = 299.03482 m/z; found = 299.03521 m/z.

**Synthesis of (*E*)-3-((4-(trifluoromethyl)phenyl)sulfonyl)acrylamide (11):**

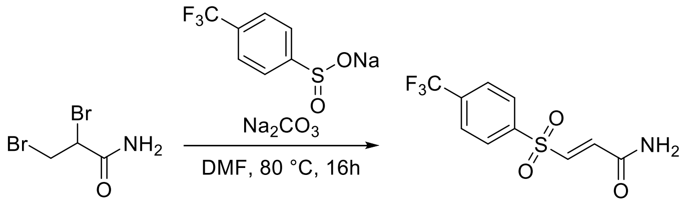


Following the general procedure **A** on 230.9 mg (1.00 mmol) of **2,3-dibromopropanecarboxamide**, purification afforded 209.5 mg (75% yield) of **(*E*)-3-((4-(trifluoromethyl)phenyl)sulfonyl)acrylamide** as a white solid. (Gradient from 0 to 40% EtOAc/hexanes)

$^1$H NMR (700 MHz, DMSO-$d_6$) δ 8.16 (d, *J* = 8.2 Hz, 2H), 8.08 (d, *J* = 8.3 Hz, 2H), 8.06 (s, 1H), 7.73 (s, 1H), 7.57 (d, *J* = 15.0 Hz, 1H), 7.07 (d, *J* = 15.0 Hz, 1H).

$^{13}$C NMR (176 MHz, DMSO-$d_6$) δ 163.33, 143.30, 138.67, 137.25, 134.23, 129.33, 127.43, 123.01.

HRMS (ESI$^+$): [M+H]$^+$ calculated for $C_{10}H_9F_3NO_3S$ = 280.02498 m/z; found = 280.02643 m/z.

**Synthesis of (*E*)-3-(quinolin-8-ylsulfonyl)acrylonitrile (12):**

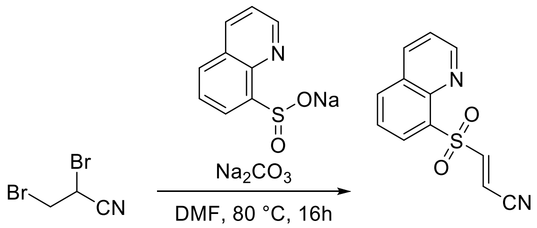


Following the general procedure **A** on 212.9 mg (1.00 mmol) of **2,3-dibromopropanenitrile**, purification afforded 75.6 mg of **(*E*)-3-(quinolin-8-ylsulfonyl)acrylonitrile** as a brown solid. (Gradient from 0 to 20% EtOAc/hexanes)

$^1$H NMR (700 MHz, Chloroform-*d*) δ 9.11 (dd, *J* = 4.2, 1.7 Hz, 1H), 8.53 (dd, *J* = 7.3, 1.5 Hz, 1H), 8.31 (dd, *J* = 8.3, 1.8 Hz, 1H), 8.25 – 8.17 (m, 2H), 7.74 (dd, *J* = 8.2, 7.4 Hz, 1H), 7.62 (dd, *J* = 8.3, 4.2 Hz, 1H), 6.70 (d, *J* = 15.9 Hz, 1H).

$^{13}$C NMR (176 MHz, Chloroform-*d*) δ 152.08, 150.46, 144.02, 136.96, 135.67, 135.60, 131.89, 129.14, 126.00, 122.92, 114.01, 111.55.

HRMS (ESI$^+$): $[M+Na]^+$ calculated for $C_{12}H_8N_2O_2SNa$ = 267.01982 m/z; found = 267.02129 m/z.

**Synthesis of methyl (*E*)-3-((4-fluorophenyl)sulfonyl)acrylate (13):**

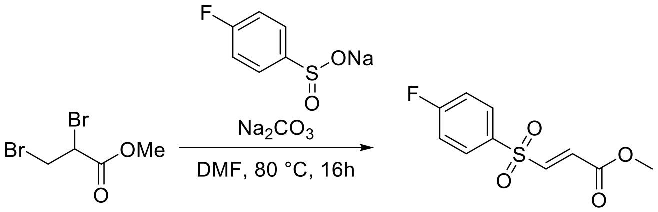


Following the general procedure **A** on 245.9 mg (1.00 mmol) of **methyl 2,3-dibromopropanate**, purification afforded 131.2 mg (34% yield) of **methyl (*E*)-3-((4-fluorophenyl)sulfonyl)acrylate** as a white solid. (Gradient from 0 to 20% EtOAc/hexanes)

$^1$H NMR (700 MHz, Chloroform-*d*) δ 7.94 – 7.91 (m, 2H), 7.31 (d, *J* = 15.2 Hz, 1H), 7.28 – 7.23 (m, 2H), 6.83 (d, *J* = 15.1 Hz, 1H), 3.79 (s, 3H).

$^{13}$C NMR (176 MHz, Chloroform-*d*) δ 167.15, 165.68, 163.94, 143.39, 131.46, 130.83, 117.21, 52.99.

HRMS (ESI$^+$): $[M+Na]^+$ calculated for $C_{10}H_9FO_4SNa$ = 267.00982 m/z; found = 267.01055 m/z.

**Synthesis of methyl (*E*)-3-(quinolin-8-ylsulfonyl)acrylate (14):**

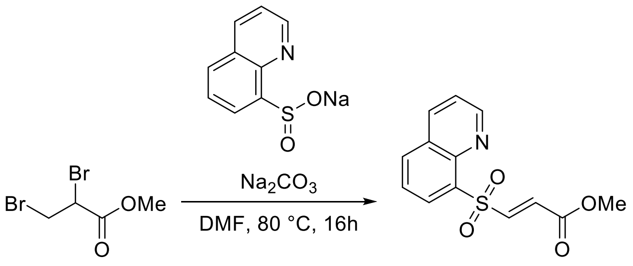


Following the general procedure **A** on 245.9 mg (1.00 mmol) of **2,3-dibromopropanecarboxamide**, purification afforded 139.4 mg (50% yield) of **methyl (*E*)-3-(quinolin-8-ylsulfonyl)acrylate** as a brown solid. (Gradient from 0 to 20% EtOAc/hexanes)

$^1$H NMR (700 MHz, DMSO-$d_6$) δ 9.14 (dd, $J$ = 4.2, 1.8 Hz, 1H), 8.63 – 8.59 (m, 1H), 8.49 (dd, $J$ = 7.3, 1.4 Hz, 1H), 8.45 (dd, $J$ = 8.3, 1.4 Hz, 1H), 8.15 (d, $J$ = 15.4 Hz, 1H), 7.87 (dd, $J$ = 8.1, 7.3 Hz, 1H), 7.76 (dd, $J$ = 8.3, 4.2 Hz, 1H), 6.92 (d, $J$ = 15.4 Hz, 1H), 3.72 (s, 3H).

$^{13}$C NMR (176 MHz, DMSO-$d_6$) δ 164.28, 152.51, 144.21, 143.49, 137.73, 136.27, 135.87, 131.77, 131.65, 129.18, 126.60, 123.46, 53.16.

HRMS (ESI$^+$): [M+H]$^+$ calculated for $C_{13}H_{12}NO_4S$ = 278.04818 m/z; found = 278.04803 m/z.

HRMS (ESI$^+$): [M+Na]$^+$ calculated for $C_{13}H_{11}NO_4SNa$ = 300.03012 m/z; found = 300.03023 m/z.

**Synthesis of (*E*)-3-(quinolin-8-ylsulfonyl)acrylamide (15):**

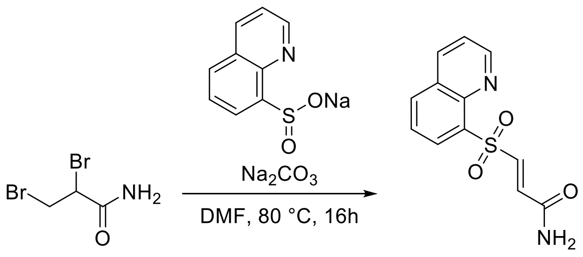


Following the general procedure **A** on 230.9 mg (1.00 mmol) of **2,3-dibromopropanecarboxamide**, purification afforded 43.3 mg (17% yield) of **(*E*)-3-(quinolin-8-ylsulfonyl)acrylamide** as a brown solid. (Gradient from 0 to 20% EtOAc/hexanes)

$^1$H NMR (700 MHz, DMSO-$d_6$) δ 9.14 (dd, $J$ = 4.2, 1.7 Hz, 1H), 8.61 (dd, $J$ = 8.4, 1.8 Hz, 1H), 8.45 (ddd, $J$ = 24.6, 7.8, 1.5 Hz, 2H), 8.05 (s, 1H), 7.94 (d, $J$ = 15.2 Hz, 1H), 7.86 (dd, $J$ = 8.2, 7.3 Hz, 1H), 7.76 (dd, $J$ = 8.3, 4.2 Hz, 1H), 7.67 (s, 1H), 7.12 (d, $J$ = 15.2 Hz, 1H).

$^{13}$C NMR (176 MHz, DMSO-$d_6$) δ 163.69, 152.41, 143.52, 140.33, 137.68, 136.56, 136.35, 135.98, 131.32, 129.19, 126.56, 123.40.

HRMS (ESI$^+$): [M+Na]$^+$ calculated for $C_{12}H_{10}N_2O_3SNa$ = 285.03042 m/z; found = 285.03116 m/z.

**Synthesis of (*E*)-3-((4-fluorophenyl)sulfonyl)acrylamide (16):**

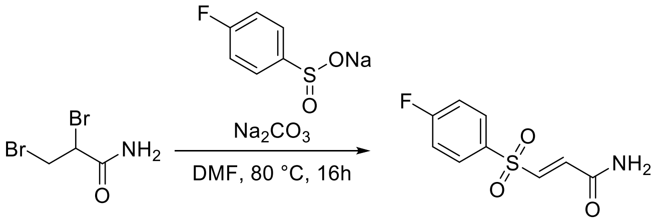


Following the general procedure **A** on 230.9 mg (1.00 mmol) of **2,3-dibromopropanecarboxamide**, purification afforded 125.0 mg (55% yield) of **(*E*)-3-((4-fluorophenyl)sulfonyl)acrylamide** as a white solid. (Gradient from 0 to 20% EtOAc/hexanes)

$^{1}$H NMR (700 MHz, DMSO-$d_6$) δ 8.03 (s, 1H), 8.02 – 7.99 (m, 2H), 7.69 (s, 1H), 7.56 – 7.52 (m, 2H), 7.49 (d, $J$ = 15.0 Hz, 1H), 6.98 (d, $J$ = 15.0 Hz, 1H).

$^{13}$C NMR (176 MHz, DMSO-$d_6$) δ 165.82, 163.50, 139.41, 135.89, 135.70, 131.58, 117.57.

HRMS (ESI$^{+}$): [M+H]$^{+}$ calculated for $C_9H_9FNO_3S$ = 230.02818 m/z; found = 230.02854 m/z.

**Synthesis of (*E*)-3-tosylacrylamide (17):**

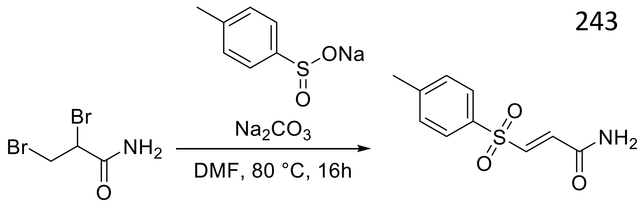


Following the general procedure **A** on 230.9 mg (1.00 mmol) of **2,3-dibromopropanecarboxamide**, purification afforded 180.5 mg (80% yield) of **(*E*)-3-tosylacrylamide** as a white solid. (Gradient from 0 to 40% EtOAc/hexanes)

$^{1}$H NMR (700 MHz, DMSO-$d_6$) δ 8.02 (s, 1H), 7.80 (d, $J$ = 7.9 Hz, 2H), 7.67 (s, 1H), 7.48 (d, $J$ = 8.0 Hz, 2H), 7.41 (d, $J$ = 15.1 Hz, 1H), 6.95 (d, $J$ = 14.9 Hz, 1H), 2.41 (s, 3H).

$^{13}$C NMR (176 MHz, DMSO-$d_6$) δ 163.14, 145.09, 139.30, 135.96, 134.81, 130.26, 127.77, 21.13.

HRMS (ESI$^{+}$): [M+Na]$^{+}$ calculated for $C_{10}H_{11}NO_3SNa$ = 248.03522 m/z; found = 248.03636 m/z.

**Synthesis of methyl (*E*)-3-((4-(trifluoromethyl)phenyl)sulfonyl)acrylate (18):**

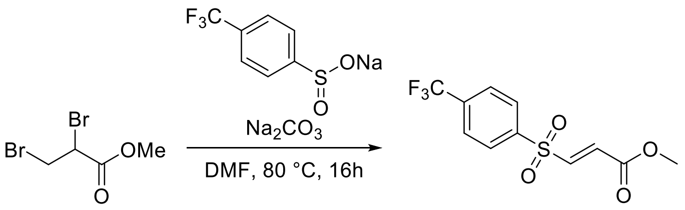


Following the general procedure **A** on 245.9 mg (1.00 mmol) of **methyl 2,3-dibromopropanate**, purification afforded 129.0 mg (44% yield) of **methyl (*E*)-3-((4-(trifluoromethyl)phenyl)sulfonyl)acrylate** as a white solid. (Gradient from 0 to 20% EtOAc/hexanes)

$^1$H NMR (700 MHz, DMSO-$d_6$) δ 8.21 – 8.16 (m, 2H), 8.10 – 8.06 (m, 2H), 7.93 (d, *J* = 15.2 Hz, 1H), 6.89 (d, *J* = 15.2 Hz, 1H), 3.74 (s, 3H).

$^{13}$C NMR (176 MHz, DMSO-$d_6$) δ 163.94, 142.71, 142.53, 134.47, 132.76, 129.65, 127.42, 123.63, 53.19.

HRMS (ESI$^+$): $[M+H]^+$ calculated for $C_{11}H_{10}F_3O_4S$ = 295.02468 m/z; found = 295.02445 m/z.

HRMS (ESI$^+$): $[M+Na]^+$ calculated for $C_{11}H_9F_3O_4SNa$ = 317.00662 m/z; found = 317.00673 m/z.

**Synthesis of 3-((4-fluorophenyl)sulfonyl)propanenitrile (19):**

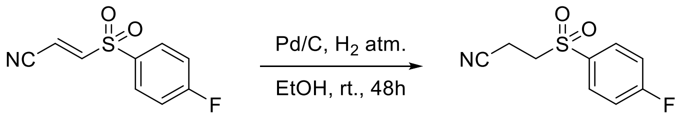


110 mg (0.52 mmol) of **(*E*)-3-((4-fluorophenyl)sulfonyl)acrylonitrile (5)** was dissolved in 5 mL ethanol and 20% w/w Pd/C was added. The reaction was stirred at room temperature under hydrogen atmosphere for 48 h and filtered through a silica pad using DCM. Purification afforded 100 mg (90% yield) of (**3-((4-fluorophenyl)sulfonyl)propanenitrile** as a white solid.

$^1$H NMR (700 MHz, DMSO-$d_6$) δ 8.03 – 7.99 (m, 2H), 7.56 – 7.51 (m, 2H), 3.75 (t, *J* = 6.9 Hz, 2H), 2.87 (t, *J* = 6.9 Hz, 2H).

$^{13}$C NMR (176 MHz, DMSO-$d_6$) δ 165.89, 134.79, 131.86, 118.14, 117.33, 50.10, 11.94.

HRMS (ESI$^+$): $[M+Na]^+$ calculated for $C_9H_8FNO_2SNa$ = 236.01522 m/z; found = 236.01540 m/z.

**Synthesis of 3-(phenylsulfonyl)pyrazine-2-carbonitrile (20):**

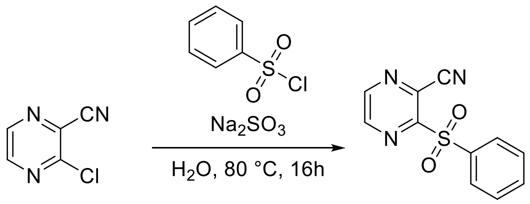


Following the general procedure **B** on 100 mg (0.72 mmol) of **3-chloropyrazine-2-carbonitrile**, purification afforded 26.2 mg (15% yield) of **3-(phenylsulfonyl)pyrazine-2-carbonitrile** as a white solid. (Gradient from 5 to 100% $MeCN/H_2O$)

$^1$H NMR (700 MHz, Chloroform-*d*) δ 8.86 (d, $J$ = 2.3 Hz, 1H), 8.82 (d, $J$ = 2.3 Hz, 1H), 8.19 – 8.16 (m, 2H), 7.73 (ddt, $J$ = 8.7, 7.4, 1.3 Hz, 1H), 7.65 – 7.61 (m, 2H).

$^{13}$C NMR (176 MHz, Chloroform-*d*) δ 157.14, 147.06, 145.92, 137.03, 135.26, 129.69, 128.40, 113.05.

HRMS (ESI$^+$): [M+H]$^+$ calculated for $C_{11}H_8N_3O_2S$ = 246.03318 m/z; found = 246.03185 m/z.

HRMS (ESI$^+$): [M+Na]$^+$ calculated for $C_{11}H_7N_3O_2SNa$ = 268.01512 m/z; found = 268.01459 m/z.

**Synthesis of 3-((4-chlorophenyl)sulfonyl)pyrazine-2-carbonitrile (24):**

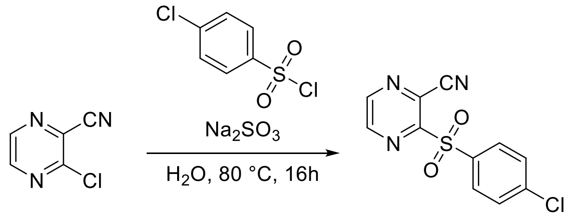


Following the general procedure **B** on 100 mg (0.72 mmol) of **3-chloropyrazine-2-carbonitrile**, purification afforded 11.6 mg (6% yield) of **3-((4-chlorophenyl)sulfonyl)pyrazine-2-carbonitrile** as a white solid. (Gradient from 5 to 100% $MeCN/H_2O$)

$^1$H NMR (700 MHz, DMSO-$d_6$) δ 9.12 – 9.09 (m, 1H), 9.04 – 9.01 (m, 1H), 8.06 (d, $J$ = 8.3 Hz, 2H), 7.81 (d, $J$ = 8.4 Hz, 2H).

$^{13}$C NMR (176 MHz, DMSO-$d_6$) δ 155.10, 148.63, 147.42, 140.65, 135.34, 131.20, 130.01, 126.82, 113.80.

HRMS (ESI$^+$): [M+Na]$^+$ calculated for $C_{11}H_6ClN_3O_2SNa$ = 301.97612 m/z; found = 301.97751 m/z.

**Synthesis of 3-((4-bromophenyl)sulfonyl)pyrazine-2-carbonitrile (25):**

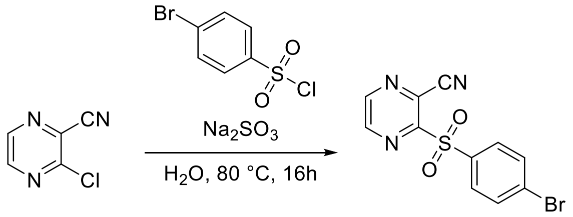


Following the general procedure **B** on 100 mg (0.72 mmol) of **3-chloropyrazine-2-carbonitrile**, purification afforded 10.0 mg (4% yield) of **3-((4-bromophenyl)sulfonyl)pyrazine-2-carbonitrile** as a white solid. (Gradient from 5 to 100% MeCN/$H_2O$)

$^1$H NMR (700 MHz, DMSO-$d_6$) δ 9.11 (d, $J$ = 2.3 Hz, 1H), 9.02 (d, $J$ = 2.3 Hz, 1H), 7.99 – 7.93 (m, 4H).

$^{13}$C NMR (176 MHz, DMSO-$d_6$) δ 155.54, 149.10, 147.91, 136.25, 133.43, 131.64, 130.40, 127.29, 114.28.

HRMS (ESI$^+$): [M+Na]$^+$ calculated for $C_{11}H_6BrN_3O_2SNa$ = 345.92562 m/z; found = 345.92701 m/z.

**Synthesis of 3-((4-(trifluoromethyl)phenyl)sulfonyl)pyrazine-2-carbonitrile (26):**

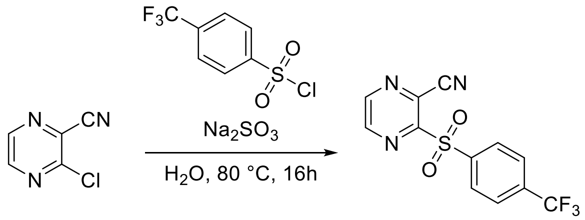


Following the general procedure **B** on 100 mg (0.72 mmol) of **3-chloropyrazine-2-carbonitrile**, purification afforded 24.0 mg (11% yield) of **3-((4-(trifluoromethyl)phenyl)sulfonyl)pyrazine-2-carbonitrile** as a white solid. (Gradient from 5 to 100% MeCN/$H_2O$)

$^1$H NMR (700 MHz, DMSO-$d_6$) δ 9.12 (d, $J$ = 2.2 Hz, 1H), 9.03 (dd, $J$ = 2.3, 1.0 Hz, 1H), 8.28 (d, $J$ = 8.2 Hz, 2H), 8.12 (d, $J$ = 8.2 Hz, 2H).

$^{13}$C NMR (176 MHz, DMSO-$d_6$) δ 154.68, 148.77, 147.44, 140.56, 134.62, 134.44, 130.39, 127.14, 126.92, 113.77.

HRMS (ESI$^+$): [M+H]$^+$ calculated for $C_{12}H_7F_3N_3O_2S$ = 314.02057 m/z; found = 314.02177 m/z.

HRMS (ESI$^+$): [M+Na]$^+$ calculated for $C_{12}H_6F_3N_3O_2SNa$ = 336.00252 m/z; found = 336.00414 m/z.

**Synthesis of 3-((2,4-difluorophenyl)sulfonyl)pyrazine-2-carbonitrile (27):**

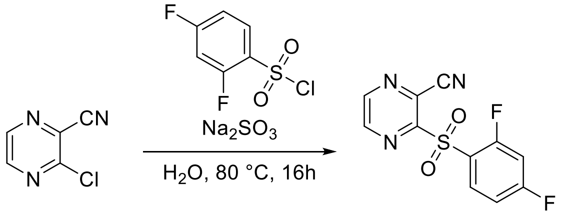


Following the general procedure **B** on 100 mg (0.72 mmol) of **3-chloropyrazine-2-carbonitrile**, purification afforded 39.8 mg (20% yield) of **3-((2,4-difluorophenyl)sulfonyl)pyrazine-2-carbonitrile** as a white solid. (Gradient from 5 to 100% MeCN/$H_2O$)

$^1$H NMR (700 MHz, DMSO-$d_6$) δ 9.18 – 9.15 (m, 1H), 9.07 – 9.04 (m, 1H), 8.19 (q, $J$ = 8.1 Hz, 1H), 7.67 (td, $J$ = 10.3, 1.9 Hz, 1H), 7.48 (ddd, $J$ = 10.0, 5.8, 2.1 Hz, 1H).

$^{13}$C NMR (176 MHz, DMSO-$d_6$) δ 167.86, 166.40, 161.23, 159.76, 154.55, 149.13, 147.55, 133.62, 126.74, 121.07, 113.48.

HRMS (ESI$^+$): [M+Na]$^+$ calculated for $C_{11}H_5F_2N_3O_2SNa$ = 303.99632 m/z; found = 303.99700 m/z.

**Synthesis of 3-((4-fluorophenyl)sulfonyl)pyrazine-2-carbonitrile (28):**

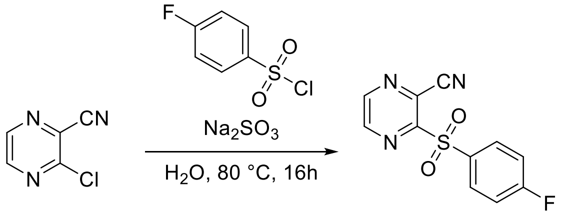


Following the general procedure **B** on 100 mg (0.72 mmol) of **3-chloropyrazine-2-carbonitrile**, purification afforded 15.7 mg (12% yield) of **3-((4-fluorophenyl)sulfonyl)pyrazine-2-carbonitrile** as a white solid. (Gradient from 5 to 100% MeCN/$H_2O$)

$^1$H NMR (700 MHz, DMSO-$d_6$) δ 9.10 (d, $J$ = 2.3 Hz, 1H), 9.02 (d, $J$ = 2.3 Hz, 1H), 8.16 – 8.11 (m, 2H), 7.61 – 7.55 (m, 2H).

$^{13}$C NMR (176 MHz, DMSO-$d_6$) δ 166.02, 155.30, 147.99, 132.79, 132.71, 126.71, 117.21, 113.83.

HRMS (ESI$^+$): [M+Na]$^+$ calculated for $C_{11}H_6FN_3O_2SNa$ = 286.00572 m/z; found = 286.00707 m/z.

**Synthesis of 3-((4-cyanophenyl)sulfonyl)pyrazine-2-carbonitrile (29):**

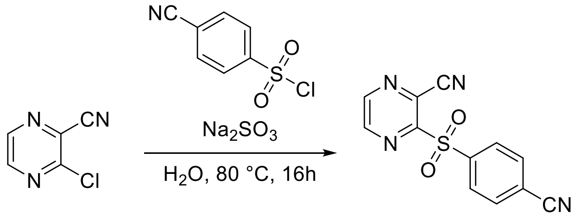


Following the general procedure **B** on 100 mg (0.72 mmol) of **3-chloropyrazine-2-carbonitrile**, purification afforded 42.8 mg (22% yield) of **3-((4-cyanophenyl)sulfonyl)pyrazine-2-carbonitrile** as a white solid. (Gradient from 5 to 100% MeCN/$H_2O$)

$^1$H NMR (700 MHz, DMSO-$d_6$) δ 9.13 (d, $J$ = 2.4 Hz, 1H), 9.03 (d, $J$ = 2.4 Hz, 1H), 8.22 (q, $J$ = 8.4 Hz, 4H).

$^{13}$C NMR (176 MHz, DMSO-$d_6$) δ 154.57, 148.82, 147.41, 140.62, 133.75, 129.99, 127.18, 117.50, 117.34, 113.74.

HRMS (ESI$^+$): [M+Na]$^+$ calculated for $C_{12}H_6N_4O_2SNa$ = 293.01032 m/z; found = 293.00973 m/z.

**Synthesis of 3-((2-fluorophenyl)sulfonyl)pyrazine-2-carbonitrile (30):**

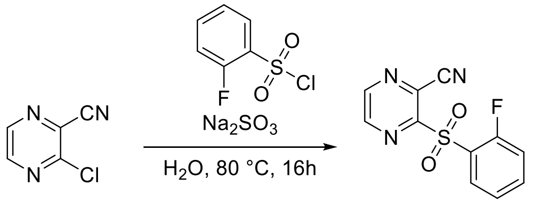


Following the general procedure **B** on 100 mg (0.72 mmol) of **3-chloropyrazine-2-carbonitrile**, purification afforded 29.2 mg (15% yield) of **3-((2-fluorophenyl)sulfonyl)pyrazine-2-carbonitrile** as a white solid. (Gradient from 5 to 100% MeCN/$H_2O$)

$^1$H NMR (700 MHz, DMSO-$d_6$) δ 9.17 (t, $J$ = 1.8 Hz, 1H), 9.05 (dd, $J$ = 2.3, 1.3 Hz, 1H), 8.15 – 8.10 (m, 1H), 7.95 – 7.90 (m, 1H), 7.58 (td, $J$ = 7.7, 1.3 Hz, 1H), 7.53 (dd, $J$ = 10.2, 8.5 Hz, 1H).

$^{13}$C NMR (176 MHz, DMSO-$d_6$) δ 159.26, 154.72, 149.14, 147.58, 138.75, 131.06, 126.70, 125.71, 124.40, 117.69, 113.48.

HRMS (ESI$^+$): [M+Na]$^+$ calculated for $C_{11}H_6FN_3O_2SNa$ = 286.00572 m/z; found = 286.00673 m/z.

**Synthesis of 3-((3,4-difluorophenyl)sulfonyl)pyrazine-2-carbonitrile (31):**

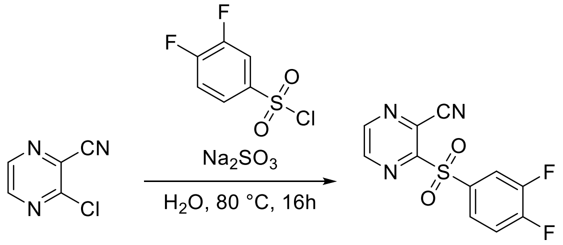


Following the general procedure **B** on 100 mg (0.72 mmol) of **3-chloropyrazine-2-carbonitrile**, purification afforded 48.4 mg (24% yield) of **3-((3,4-difluorophenyl)sulfonyl)pyrazine-2-carbonitrile** as a white solid. (Gradient from 5 to 100% MeCN/$H_2O$)

$^1$H NMR (700 MHz, DMSO-$d_6$) δ 9.12 (dd, $J$ = 2.3, 1.1 Hz, 1H), 9.03 (dd, $J$ = 2.3, 1.0 Hz, 1H), 8.14 (ddd, $J$ = 9.6, 7.0, 2.3 Hz, 1H), 8.00 – 7.96 (m, 1H), 7.86 – 7.81 (m, 1H).

$^{13}$C NMR (176 MHz, DMSO-$d_6$) δ 154.81, 154.63, 153.17, 150.21, 148.78, 148.03, 133.41, 127.70, 126.94, 119.30, 113.77.

HRMS (ESI$^+$): [M+Na]$^+$ calculated for $C_{11}H_5F_2N_3O_2SNa$ = 303.99632 m/z; found = 303.99690 m/z.

**Synthesis of 3-((3,5-difluorophenyl)sulfonyl)pyrazine-2-carbonitrile (32):**

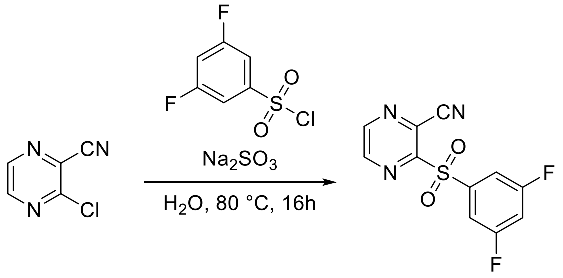


Following the general procedure **B** on 100 mg (0.72 mmol) of **3-chloropyrazine-2-carbonitrile**, purification afforded 44.8 mg (22% yield) of **3-((3,5-difluorophenyl)sulfonyl)pyrazine-2-carbonitrile** as a white solid. (Gradient from 5 to 100% MeCN/$H_2O$)

$^1$H NMR (700 MHz, DMSO-$d_6$) δ 9.14 (d, $J$ = 2.3 Hz, 1H), 9.04 (d, $J$ = 2.3 Hz, 1H), 7.88 – 7.83 (m, 1H), 7.82 – 7.77 (m, 2H).

$^{13}$C NMR (176 MHz, DMSO-$d_6$) δ 162.96, 161.52, 154.38, 148.83, 147.38, 139.70, 127.18, 113.72, 111.30.

HRMS (ESI$^+$): [M+Na]$^+$ calculated for $C_{11}H_5F_2N_3O_2SNa$ = 303.99632 m/z; found = 303.99757 m/z.

**Synthesis of 2-((4-fluorophenyl)sulfonyl)-6-methylnicotinonitrile (33):**

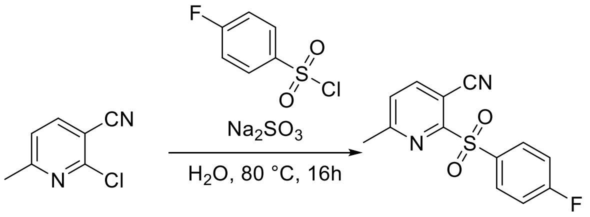


Following the general procedure **B** on 100 mg (0.66 mmol) of **2-chloro-6-methylnicotinonitrile**, purification afforded 55.9 mg (31% yield) of **2-((4-fluorophenyl)sulfonyl)-6-methylnicotinonitrile** as a white solid. (Gradient from 5 to 100% MeCN/$H_2O$)

$^1$H NMR (700 MHz, Chloroform-*d*) δ 8.20 – 8.15 (m, 2H), 8.04 (d, *J* = 8.0 Hz, 1H), 7.43 (d, *J* = 8.0 Hz, 1H), 7.28 – 7.22 (m, 2H), 2.66 (s, 3H).

$^{13}$C NMR (176 MHz, Chloroform-*d*) δ 166.46, 163.49, 159.09, 143.28, 133.83, 132.54, 126.27, 116.71, 114.15, 105.52, 24.89.

HRMS (ESI$^+$): [M+H]$^+$ calculated for $C_{13}H_{10}FN_2O_2S$ = 277.04418 m/z; found = 277.04492 m/z.

HRMS (ESI$^+$): [M+Na]$^+$ calculated for $C_{13}H_9FN_2O_2SNa$ = 299.02612 m/z; found = 299.02738 m/z.

**Synthesis of 2-((4-fluorophenyl)sulfonyl)nicotinonitrile (34):**

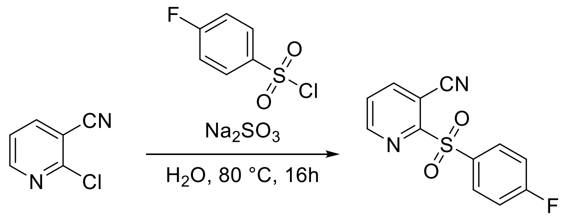


Following the general procedure **B** on 100 mg (0.73 mmol) of **2-chloronicotinonitrile**, purification afforded 53.8 mg (28% yield) of **2-((4-fluorophenyl)sulfonyl)nicotinonitrile** as a white solid. (Gradient from 5 to 100% MeCN/$H_2O$)

$^1$H NMR (700 MHz, DMSO-$d_6$) δ 8.90 (dd, *J* = 4.7, 1.6 Hz, 1H), 8.67 (dd, *J* = 7.9, 1.6 Hz, 1H), 8.13 – 8.08 (m, 2H), 7.90 (dd, *J* = 7.9, 4.8 Hz, 1H), 7.59 – 7.54 (m, 2H).

$^{13}$C NMR (176 MHz, DMSO-$d_6$) δ 166.28, 158.76, 153.31, 145.51, 133.69, 132.91, 128.12, 117.61, 114.46, 107.35.

HRMS (ESI$^+$): [M+H]$^+$ calculated for $C_{12}H_8FN_2O_2S$ = 263.02848 m/z; found = 263.02951 m/z.

**(*E*)-3-tosylacrylonitrile (1)**

$^1$H NMR (700 MHz, Chloroform-*d*)

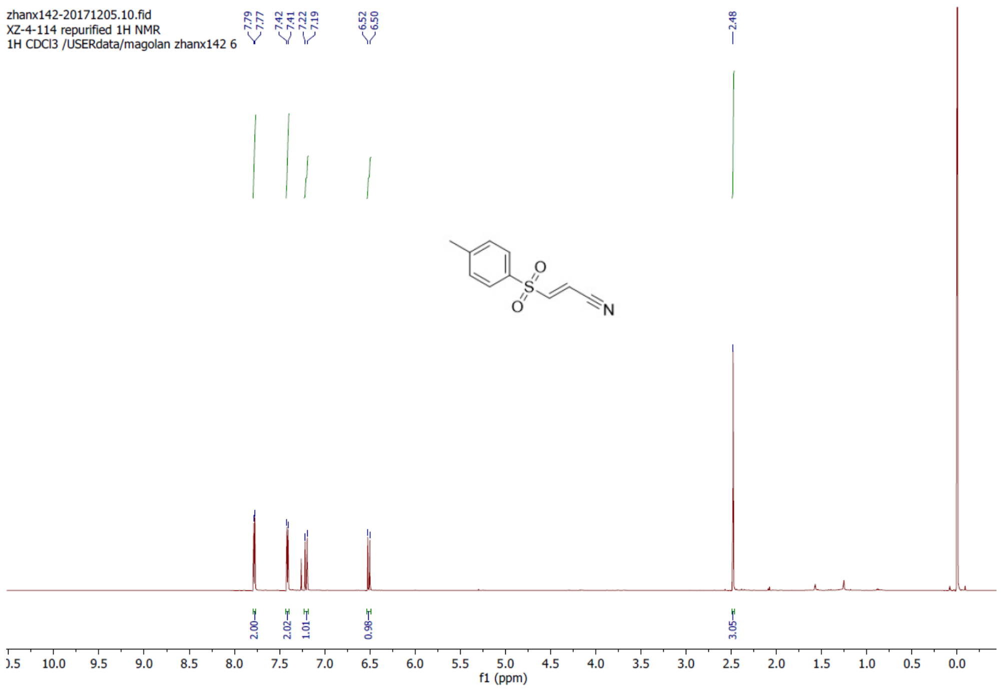


**(*E*)-3-tosylacrylonitrile (1)**

$^{13}$C NMR (176 MHz, Chloroform-*d*)

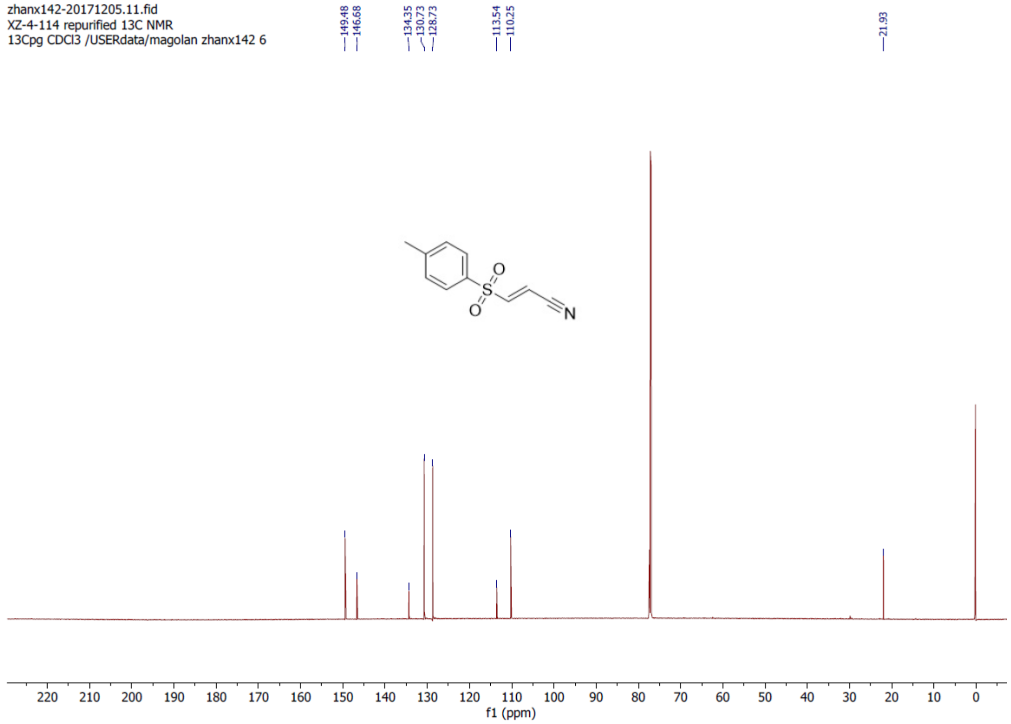

**(*E*)-3-((4-fluorophenyl)sulfonyl)acrylonitrile (5)**

$^{1}$H NMR (700 MHz, Chloroform-*d*)

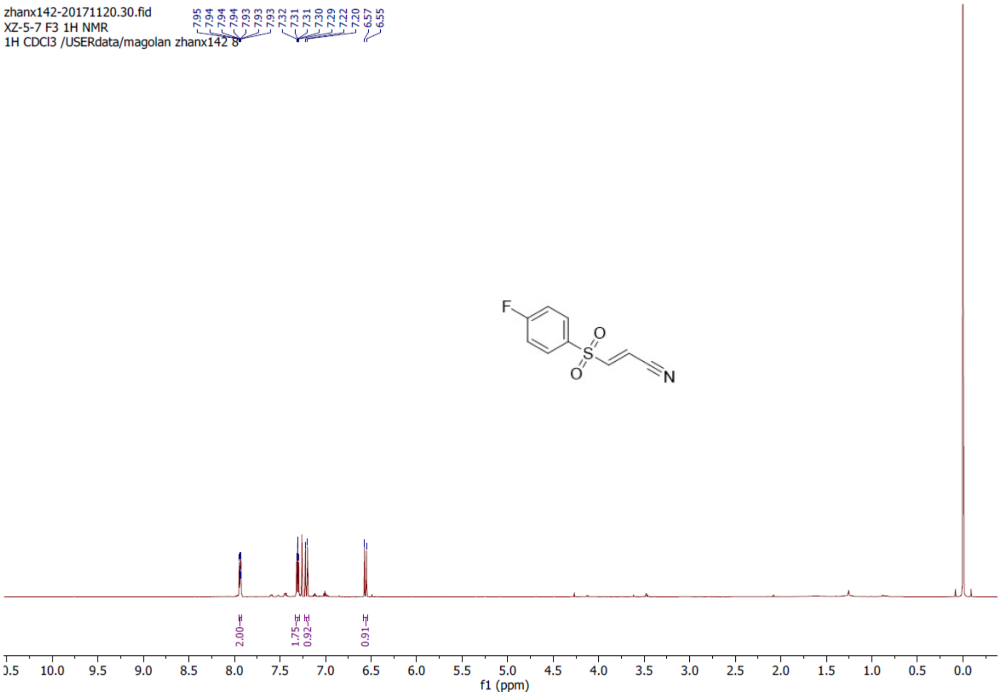


**(*E*)-3-((4-fluorophenyl)sulfonyl)acrylonitrile (5)**

$^{13}$C NMR (176 MHz, Chloroform-*d*)

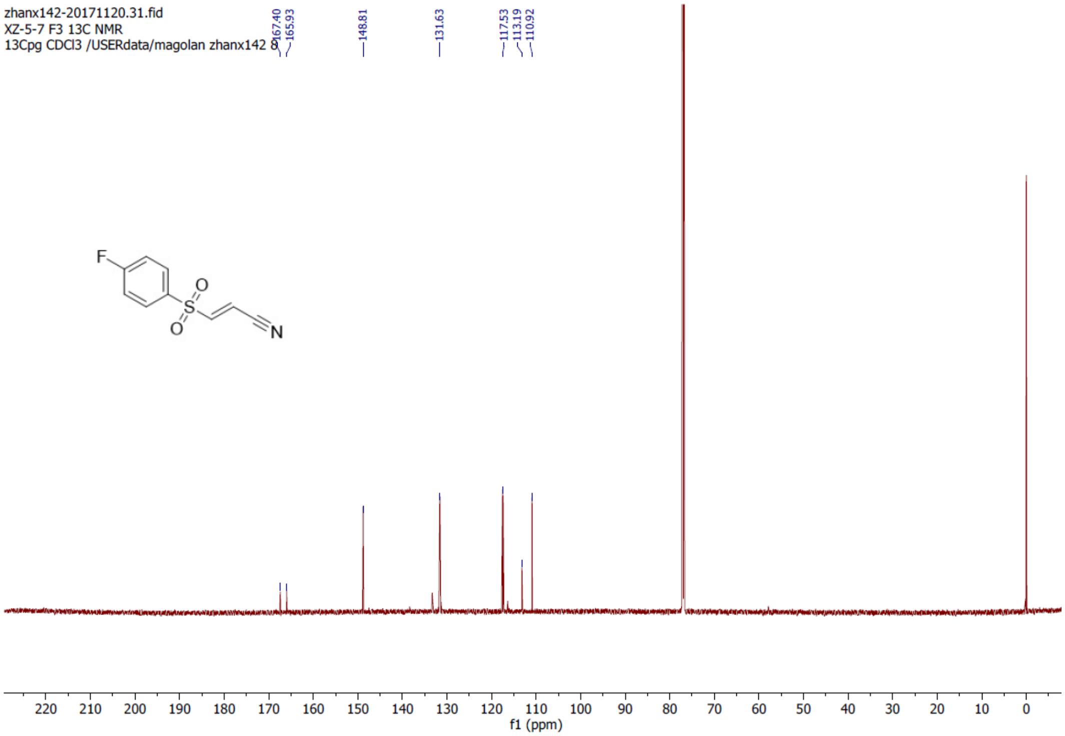

**(*E*)-3-((4-(*tert*-butyl)phenyl)sulfonyl)acrylonitrile (6)**

$^{1}H$ NMR (700 MHz, DMSO-$d_6$)

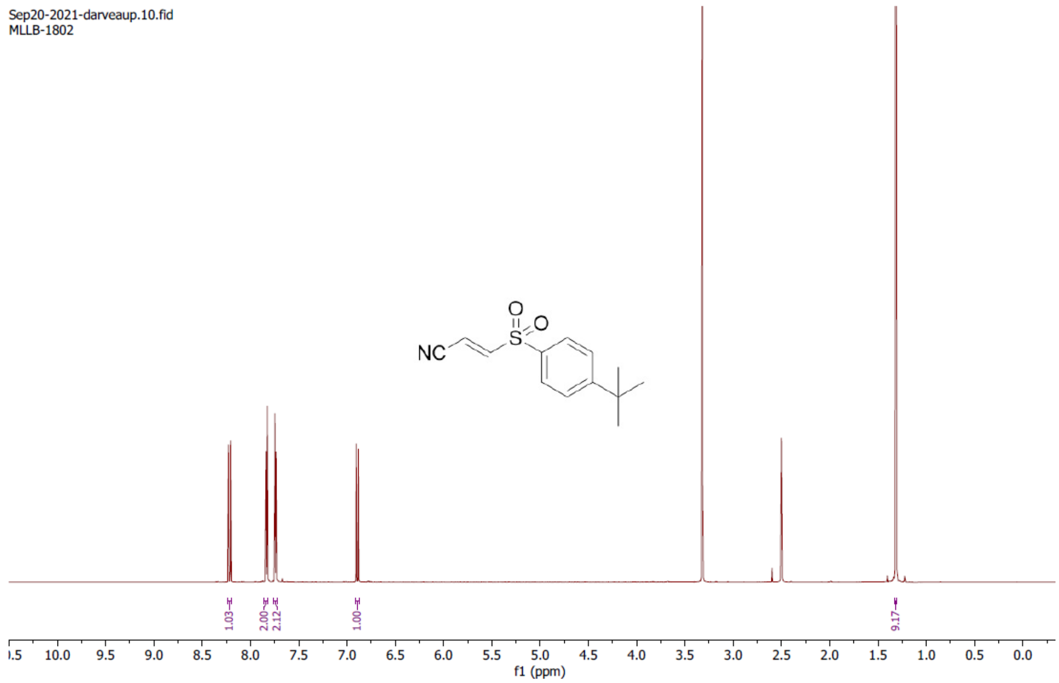


**(*E*)-3-((4-(*tert*-butyl)phenyl)sulfonyl)acrylonitrile (6)**

$^{13}C$ NMR (176 MHz, DMSO-$d_6$)

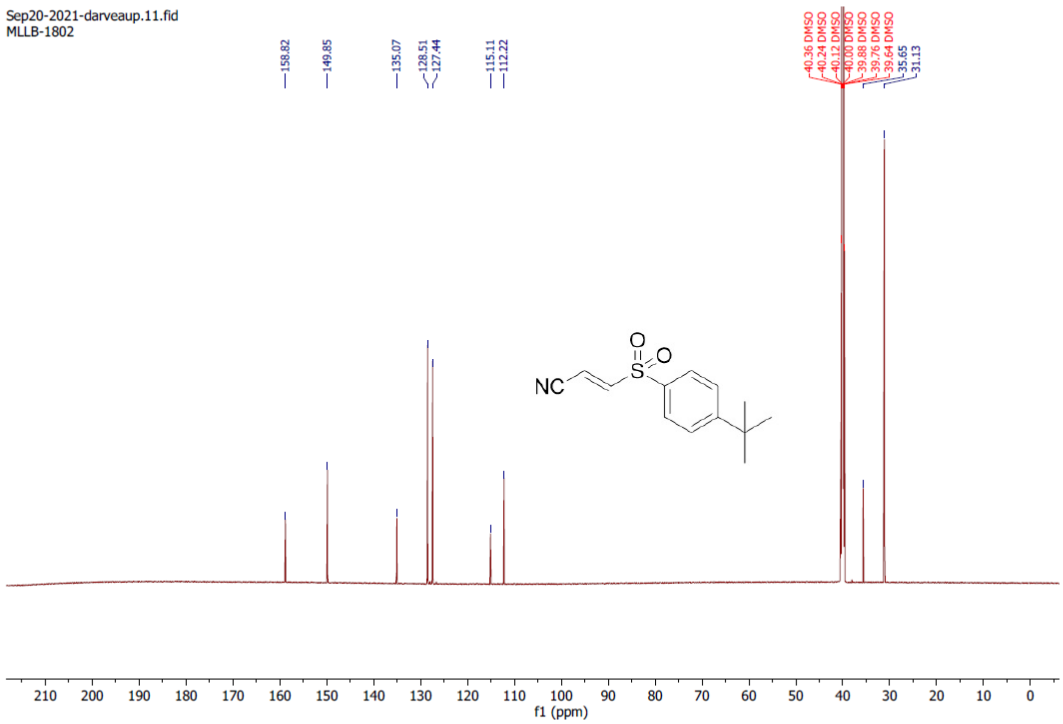

**(*E*)-3-(naphthalen-2-ylsulfonyl)acrylamide (7)**

$^{1}H$ NMR (700 MHz, DMSO-$d_6$)

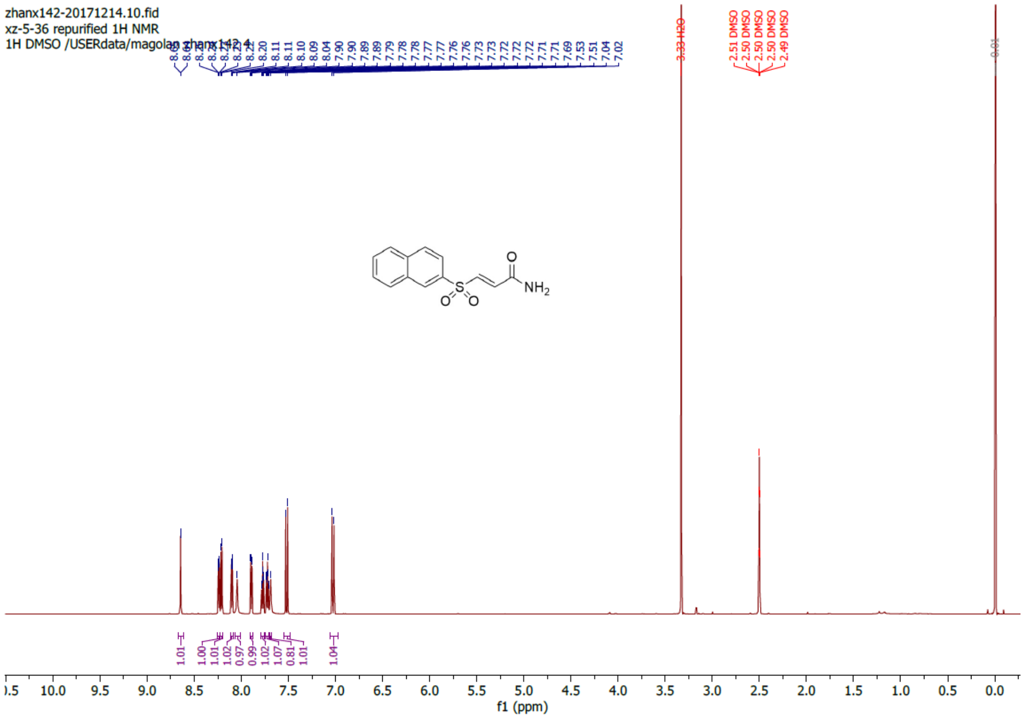


**(*E*)-3-(naphthalen-2-ylsulfonyl)acrylamide (7)**

$^{13}C$ NMR (176 MHz, DMSO-$d_6$)

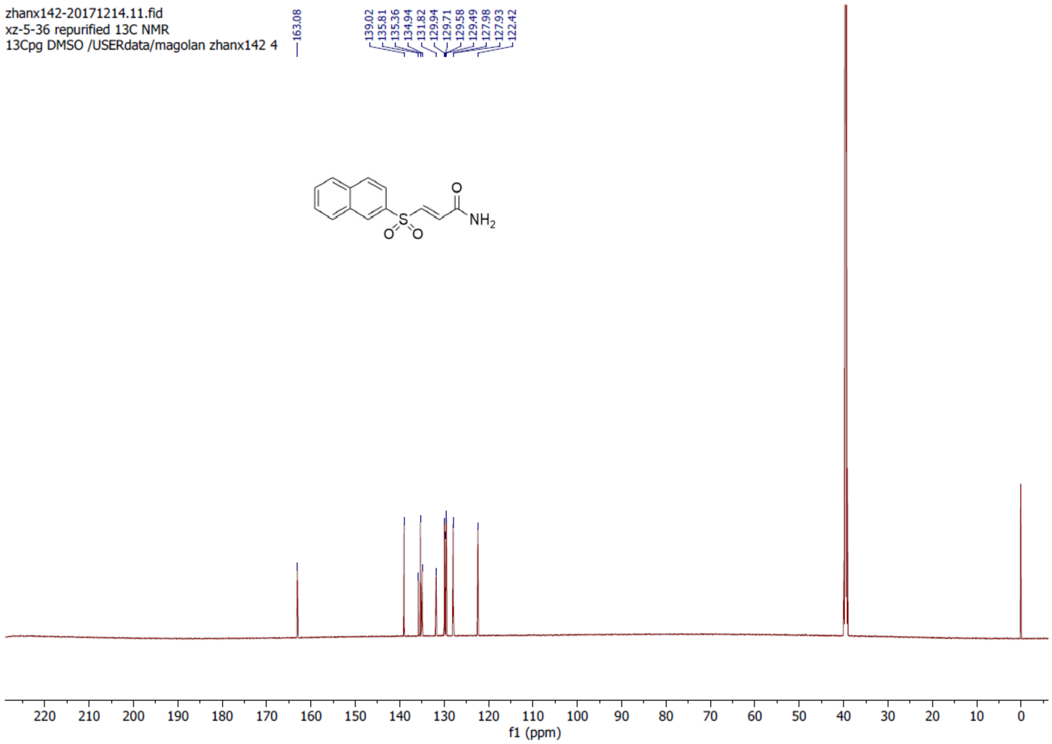

**methyl (*E*)-3-tosylacrylate (9)**

$^{1}H$ NMR (700 MHz, DMSO-$d_6$)

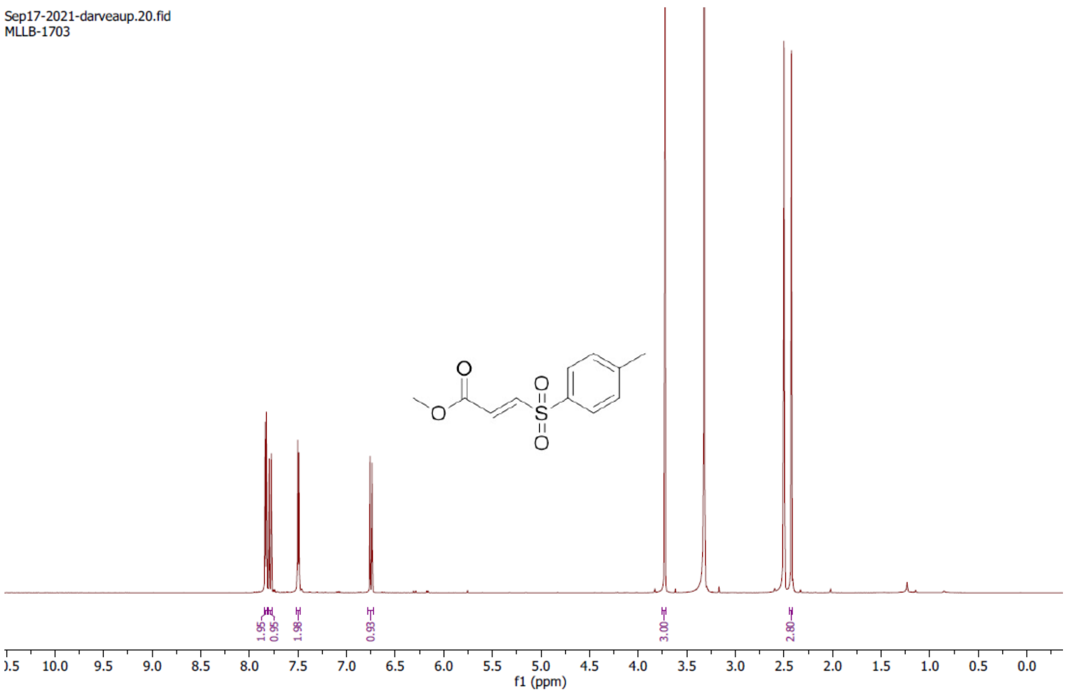


**methyl (*E*)-3-tosylacrylate (9)**

$^{13}C$ NMR (176 MHz, DMSO-$d_6$)

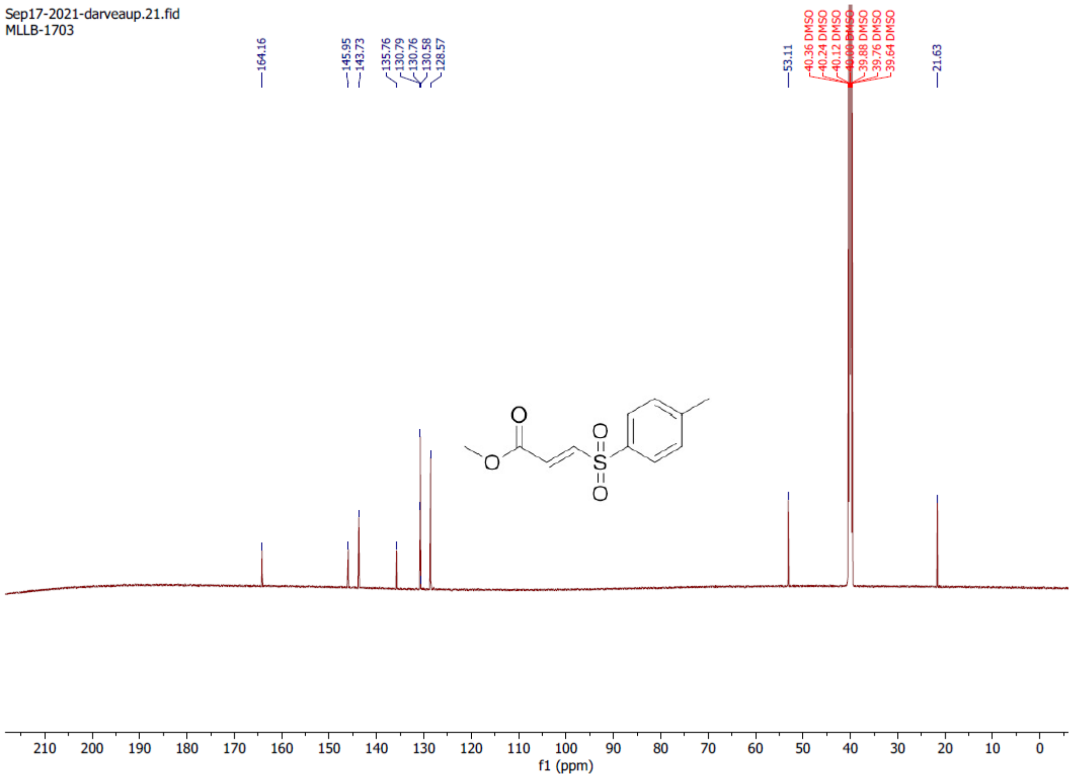

**Methyl (*E*)-3-(naphthalen-2-ylsulfonyl)acrylate (10)**

$^{1}$H NMR (700 MHz, Chloroform-*d*)

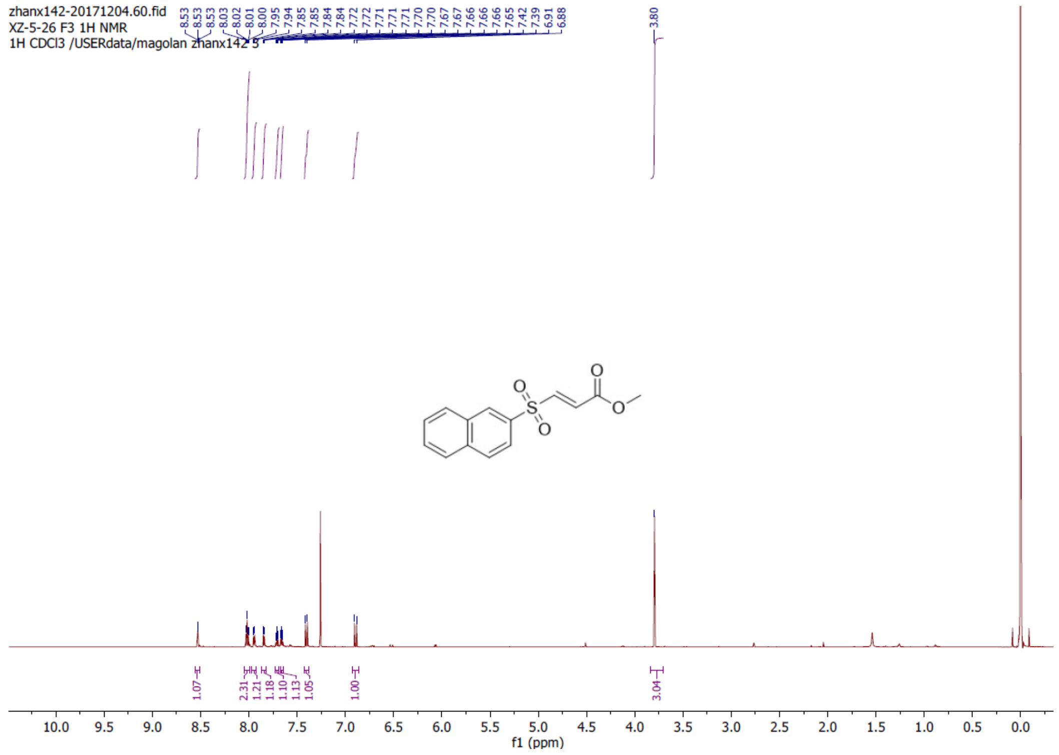


**Methyl (*E*)-3-(naphthalen-2-ylsulfonyl)acrylate (10)**

$^{13}$C NMR (176 MHz, Chloroform-*d*)

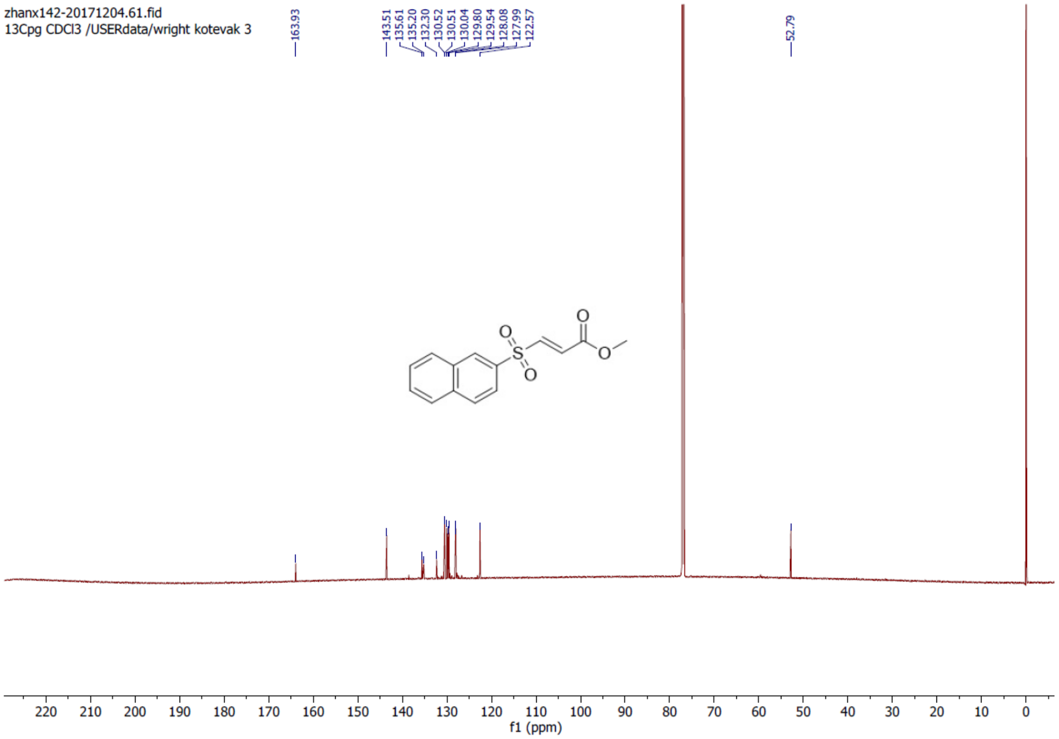

**(*E*)-3-((4-(trifluoromethyl)phenyl)sulfonyl)acrylamide (11)**

$^{1}H$ NMR (700 MHz, DMSO-$d_6$)

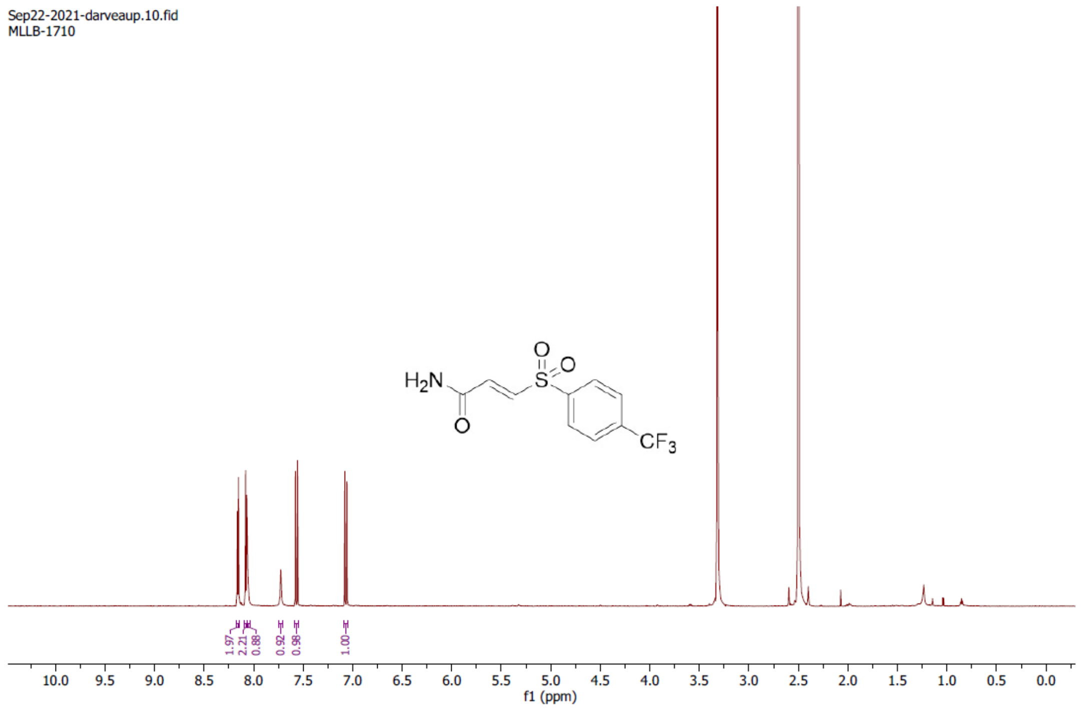


**(*E*)-3-((4-(trifluoromethyl)phenyl)sulfonyl)acrylamide (11)**

$^{13}C$ NMR (176 MHz, DMSO-$d_6$)

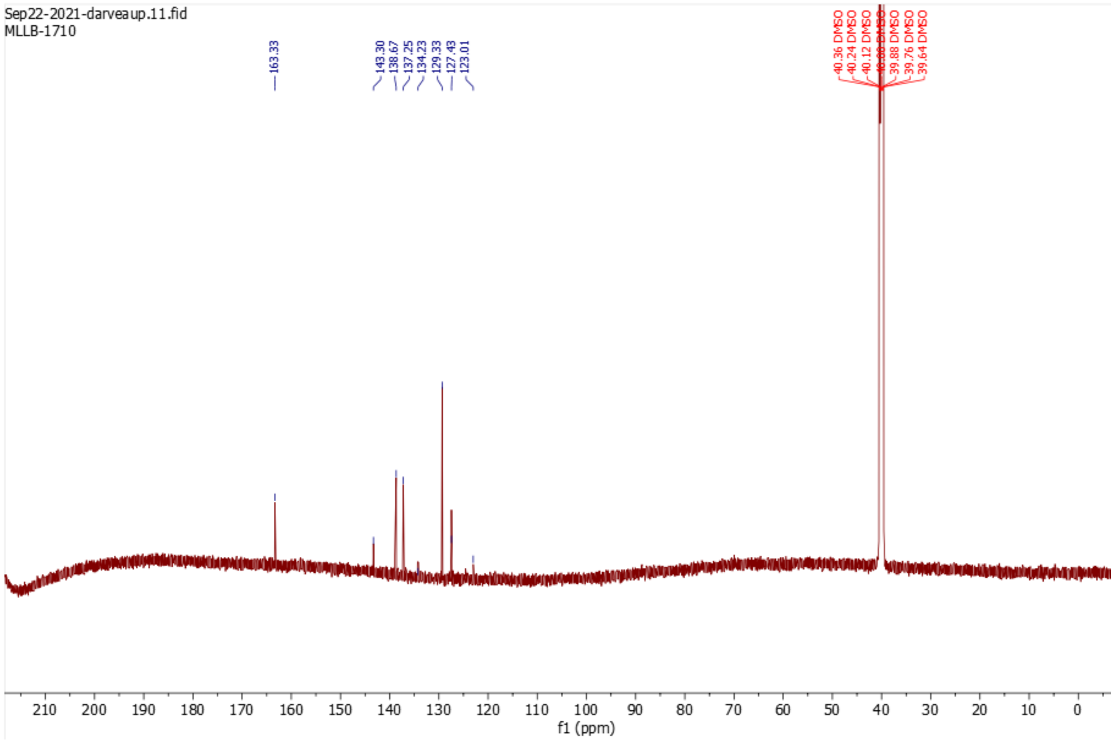

**(*E*)-3-(quinolin-8-ylsulfonyl)acrylonitrile (12)**

$^{1}$H NMR (700 MHz, Chloroform-*d*)

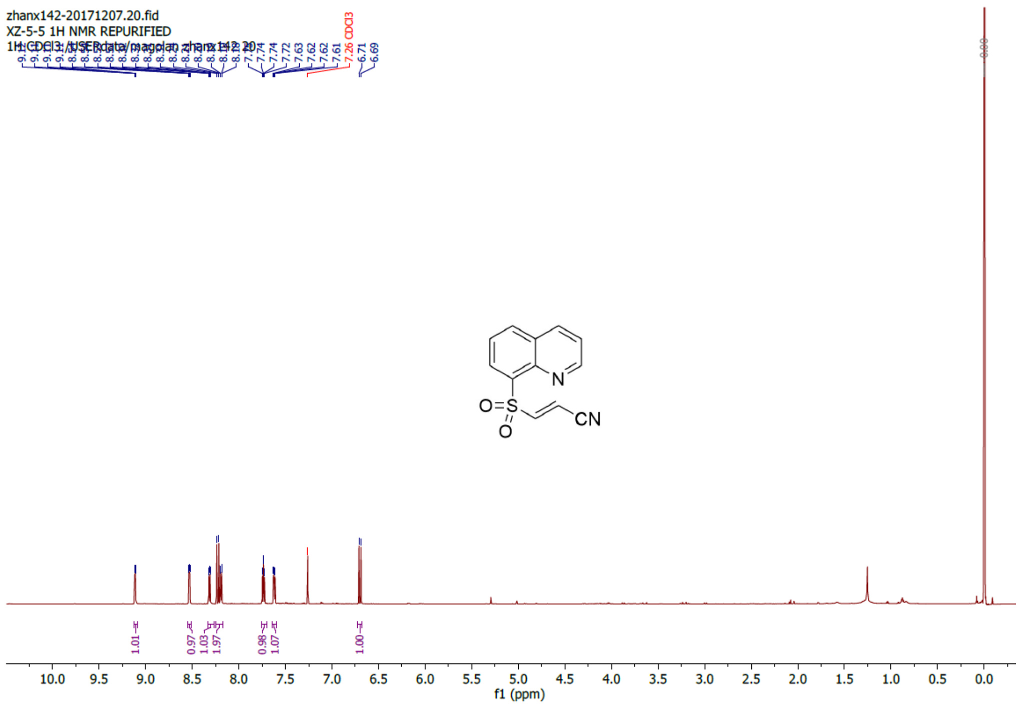


**(*E*)-3-(quinolin-8-ylsulfonyl)acrylonitrile (12)**

$^{13}$C NMR (176 MHz, Chloroform-*d*)

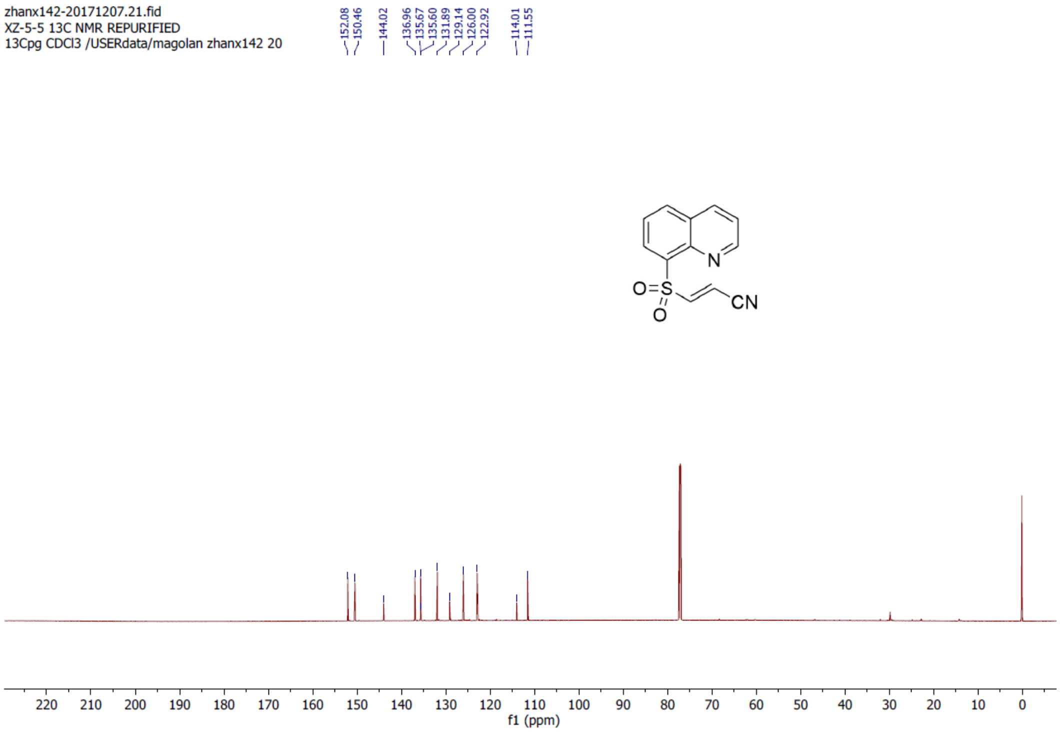

**Methyl (*E*)-3-((4-fluorophenyl)sulfonyl)acrylate (13)**

$^{1}$H NMR (700 MHz, Chloroform-*d*)

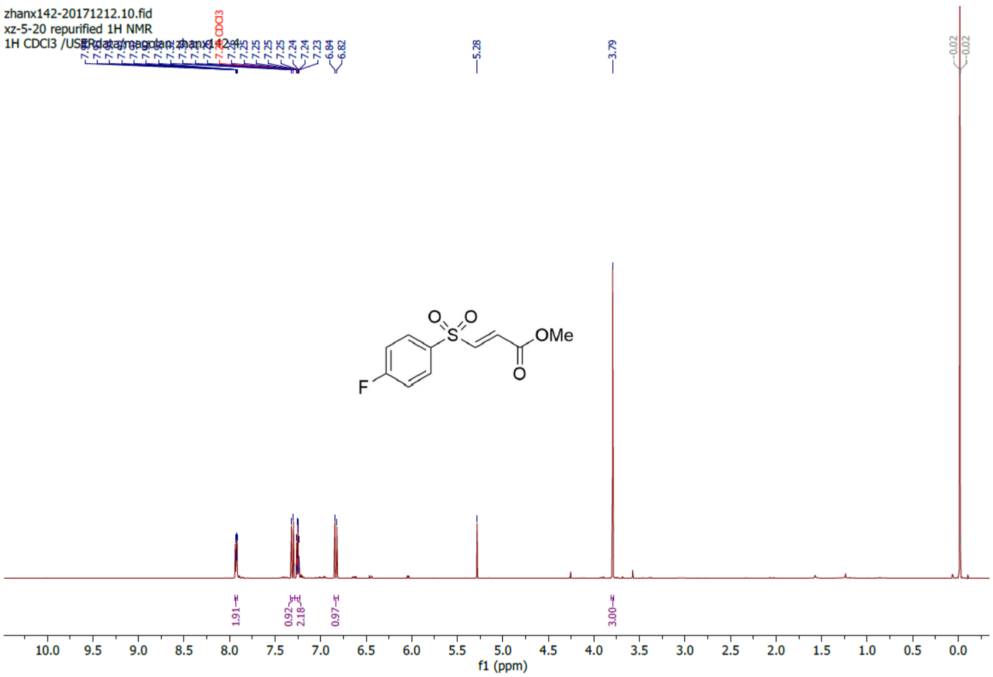


**Methyl (*E*)-3-((4-fluorophenyl)sulfonyl)acrylate (13)**

$^{13}$C NMR (176 MHz, Chloroform-*d*)

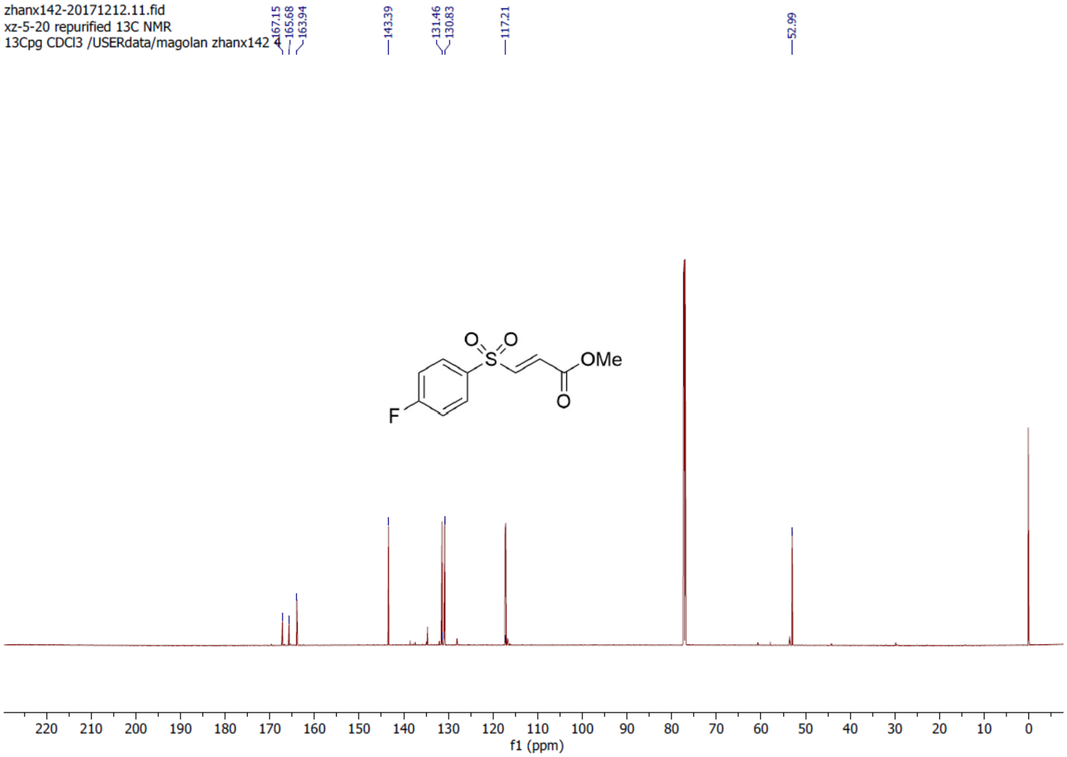

**Methyl (*E*)-3-(quinolin-8-ylsulfonyl)acrylate (14)**

$^{1}$H NMR (700 MHz, DMSO-$d_6$)

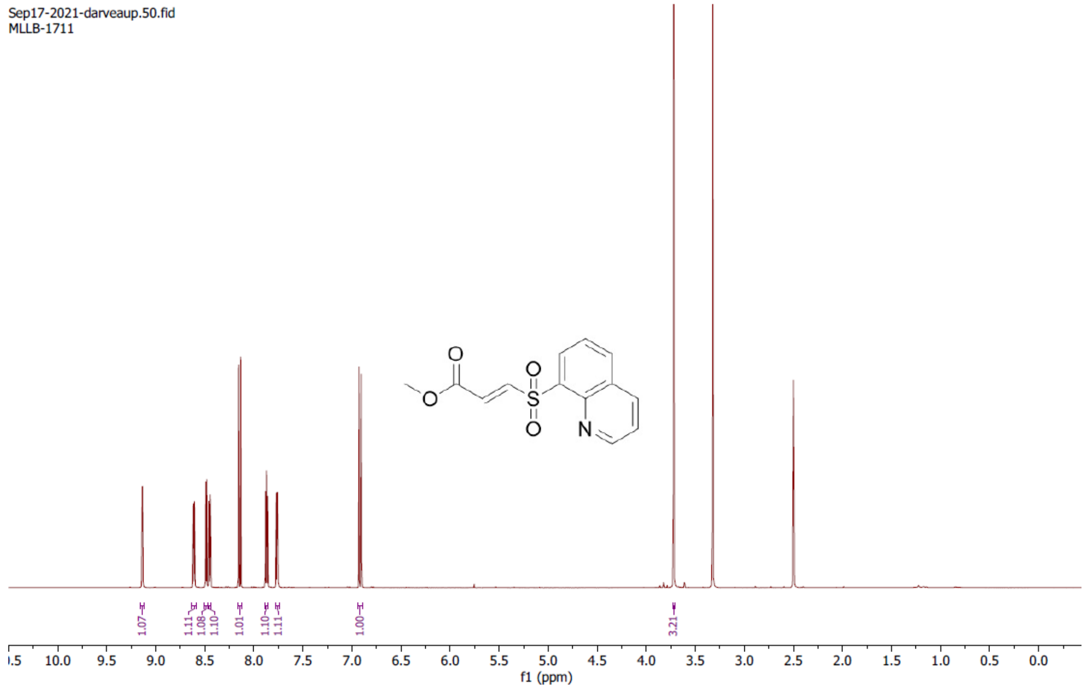


**Methyl (*E*)-3-(quinolin-8-ylsulfonyl)acrylate (14)**

$^{13}$C NMR (176 MHz, DMSO-$d_6$)

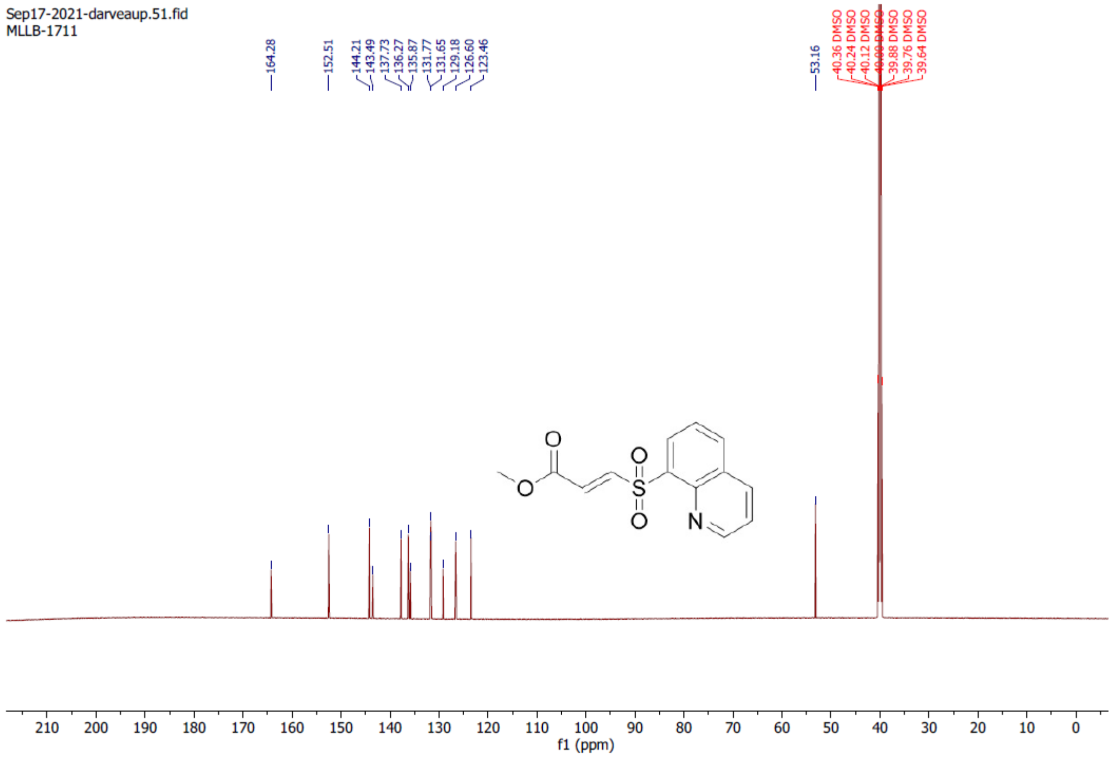

**(*E*)-3-(quinolin-8-ylsulfonyl)acrylamide (15)**

$^{1}$H NMR (700 MHz, DMSO-$d_6$)

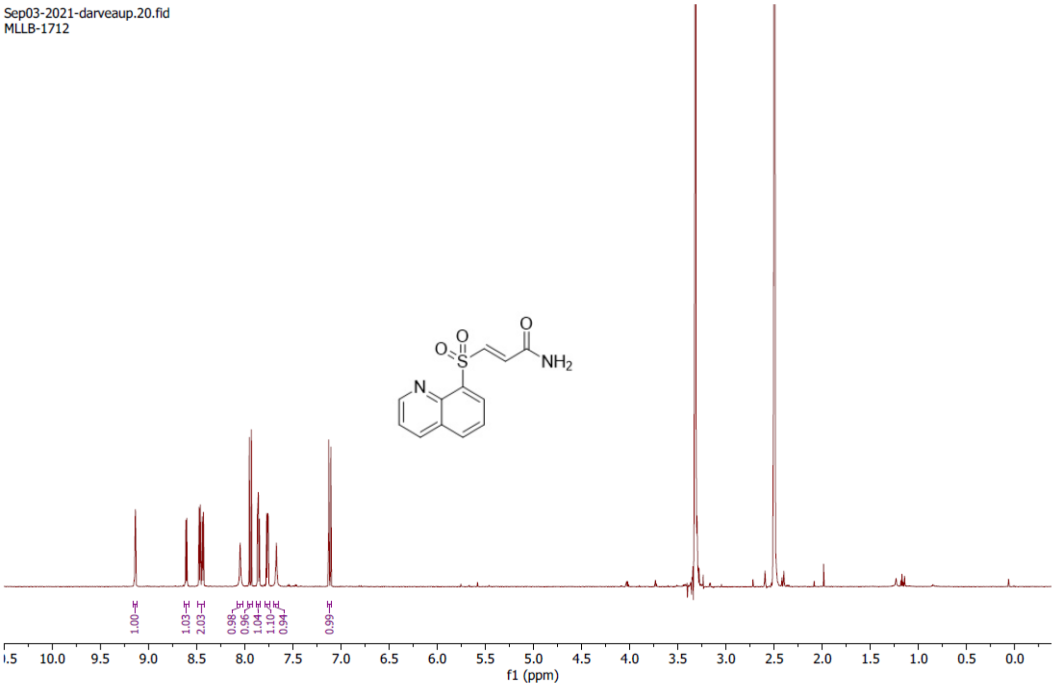


**(*E*)-3-(quinolin-8-ylsulfonyl)acrylamide (15)**

$^{13}$C NMR (176 MHz, DMSO-$d_6$)

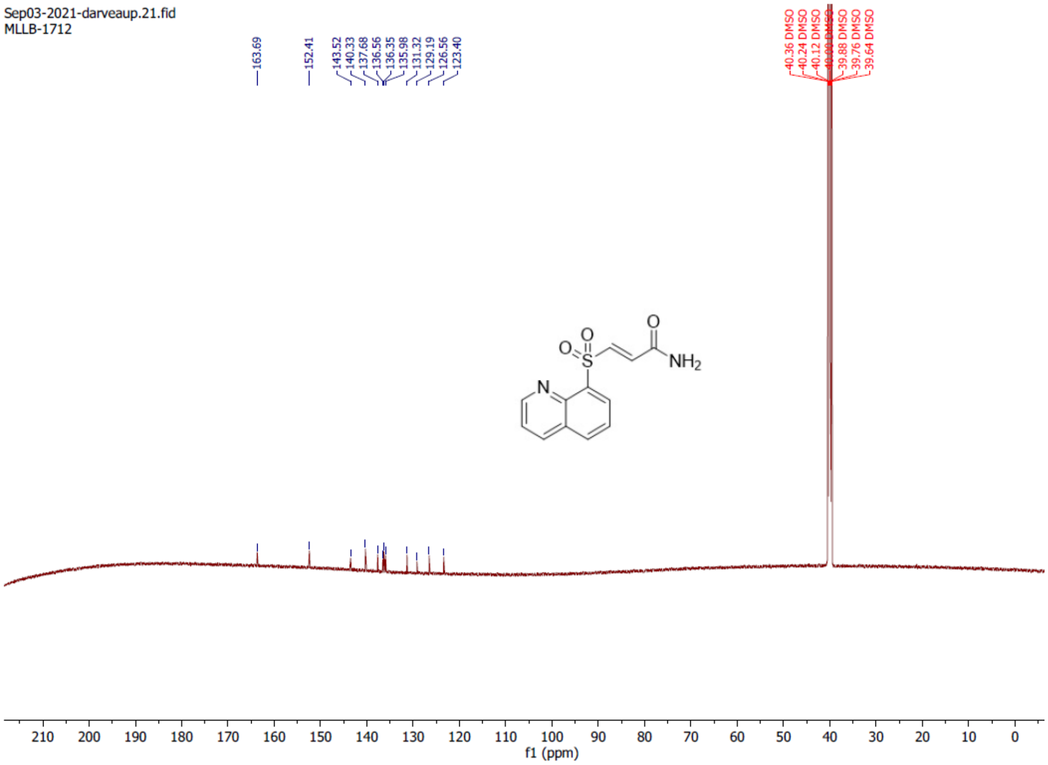

**(*E*)-3-((4-fluorophenyl)sulfonyl)acrylamide (16)**

$^{1}$H NMR (700 MHz, DMSO-$d_6$)

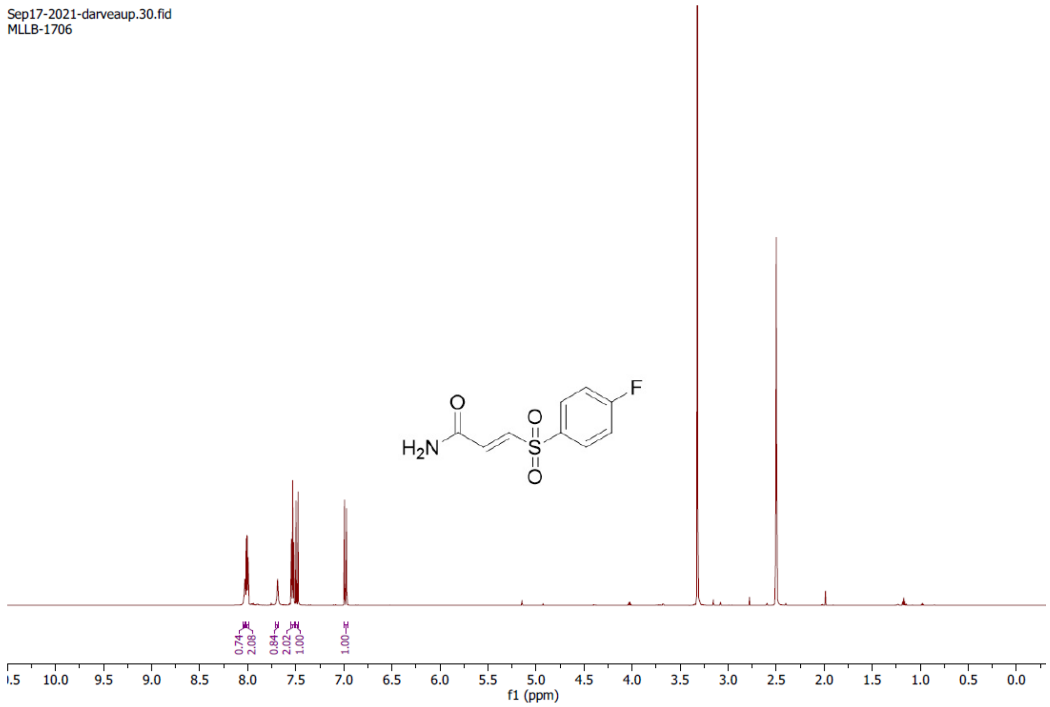


**(*E*)-3-((4-fluorophenyl)sulfonyl)acrylamide (16)**

$^{13}$C NMR (176 MHz, DMSO-$d_6$)

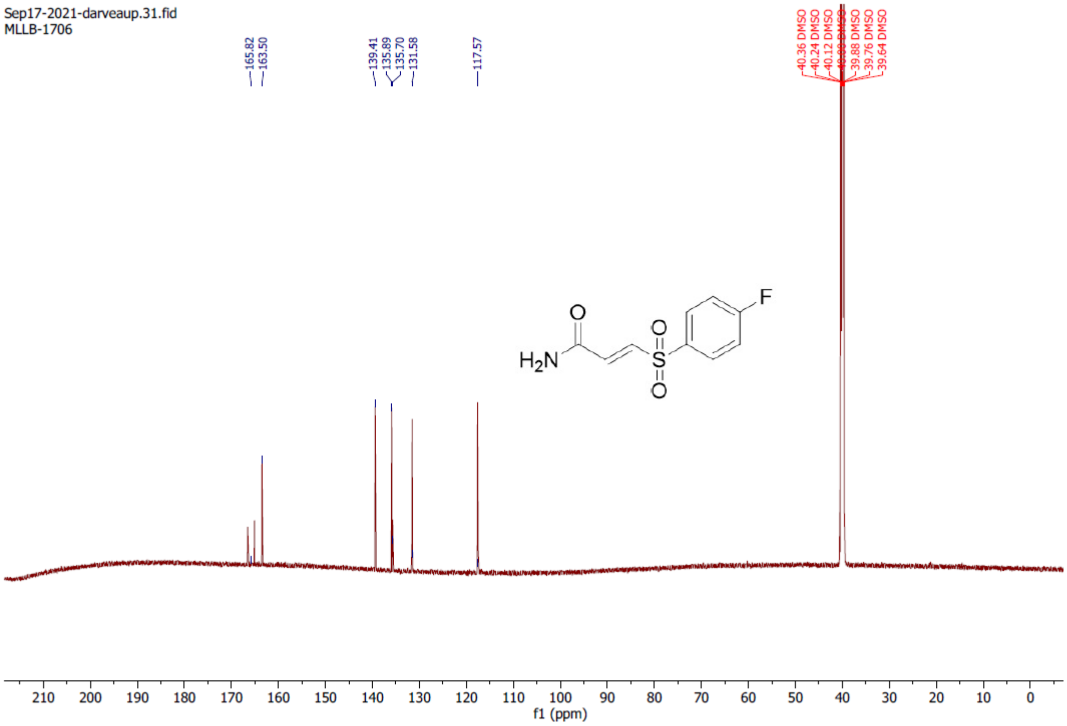

**(*E*)-3-tosylacrylamide (17)**

$^{1}$H NMR (700 MHz, DMSO-$d_6$)

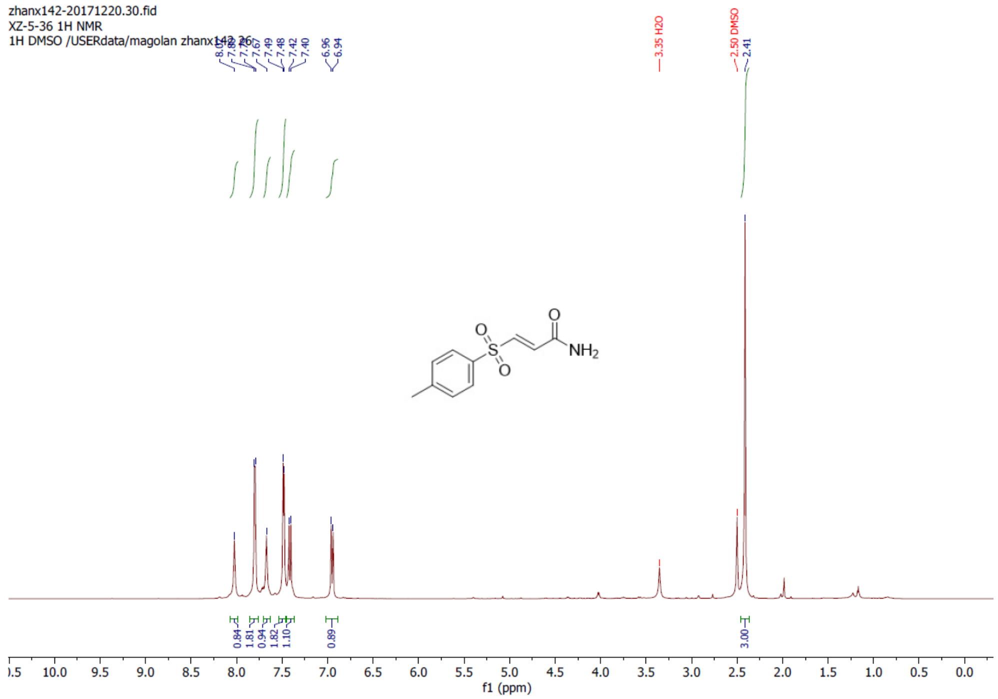


**(*E*)-3-tosylacrylamide (17)**

$^{13}$C NMR (176 MHz, DMSO-$d_6$)

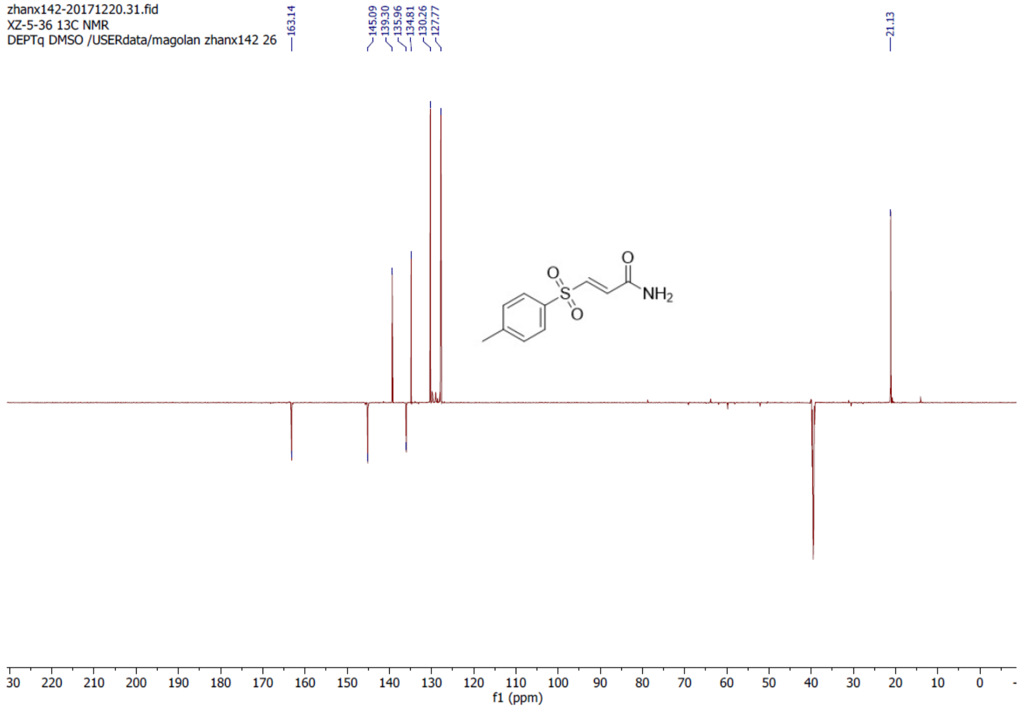

**Methyl (*E*)-3-((4-(trifluoromethyl)phenyl)sulfonyl)acrylate (18)**

$^{1}H$ NMR (700 MHz, DMSO-$d_6$)

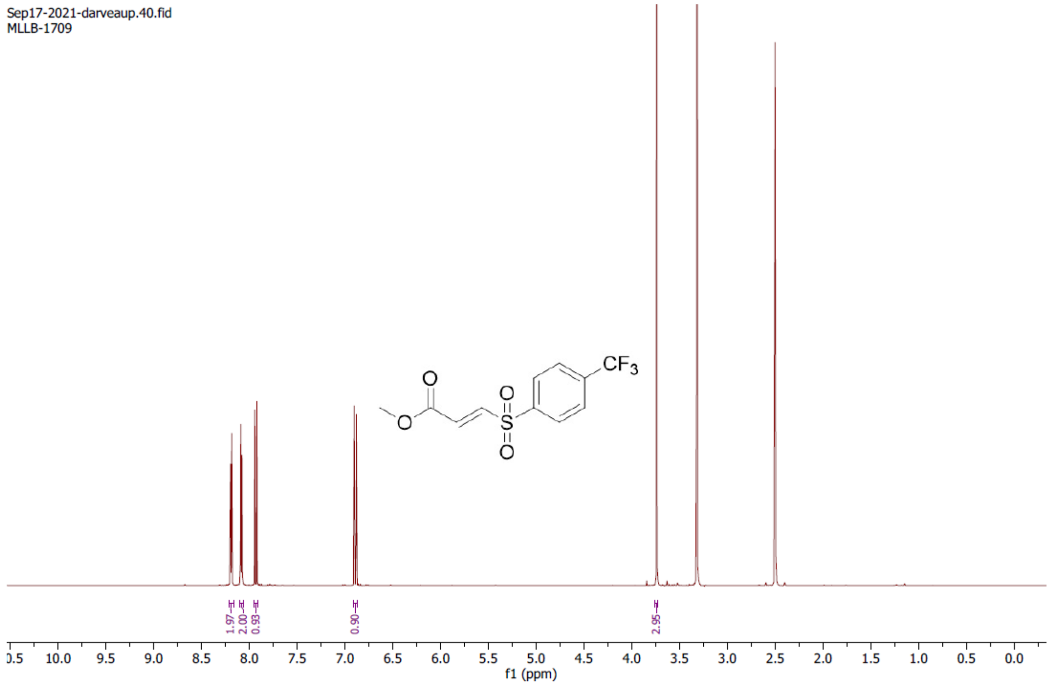


**Methyl (*E*)-3-((4-(trifluoromethyl)phenyl)sulfonyl)acrylate (18)**

$^{13}C$ NMR (176 MHz, DMSO-$d_6$)

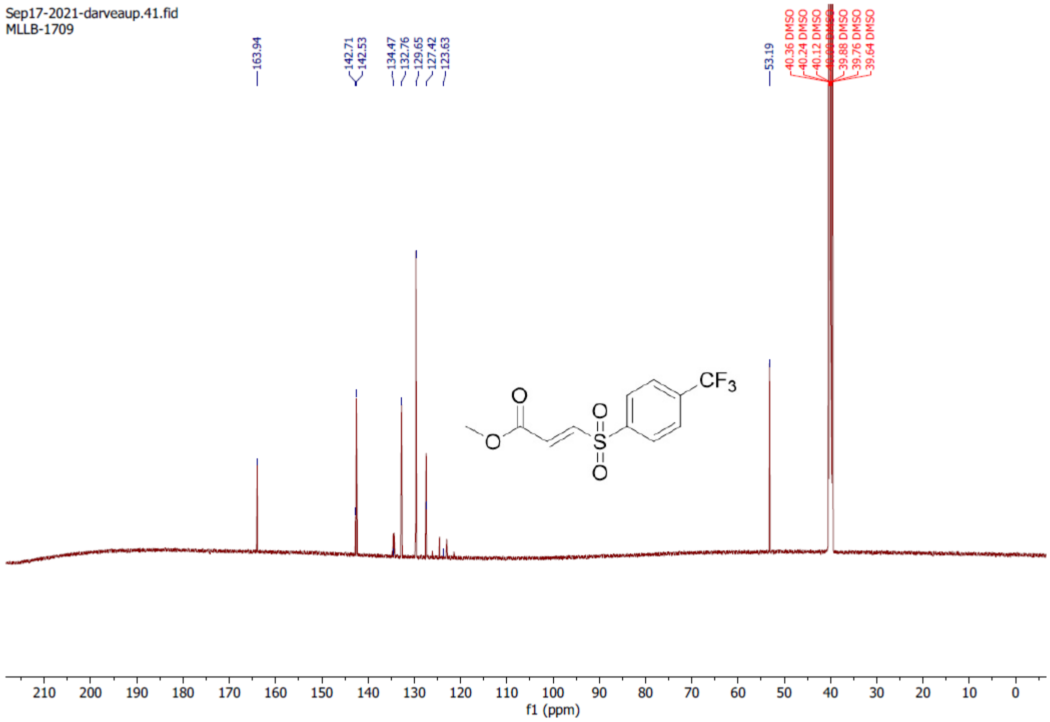

**3-((4-fluorophenyl)sulfonyl)propanenitrile (19)**

$^{1}$H NMR (700 MHz, DMSO-$d_6$)

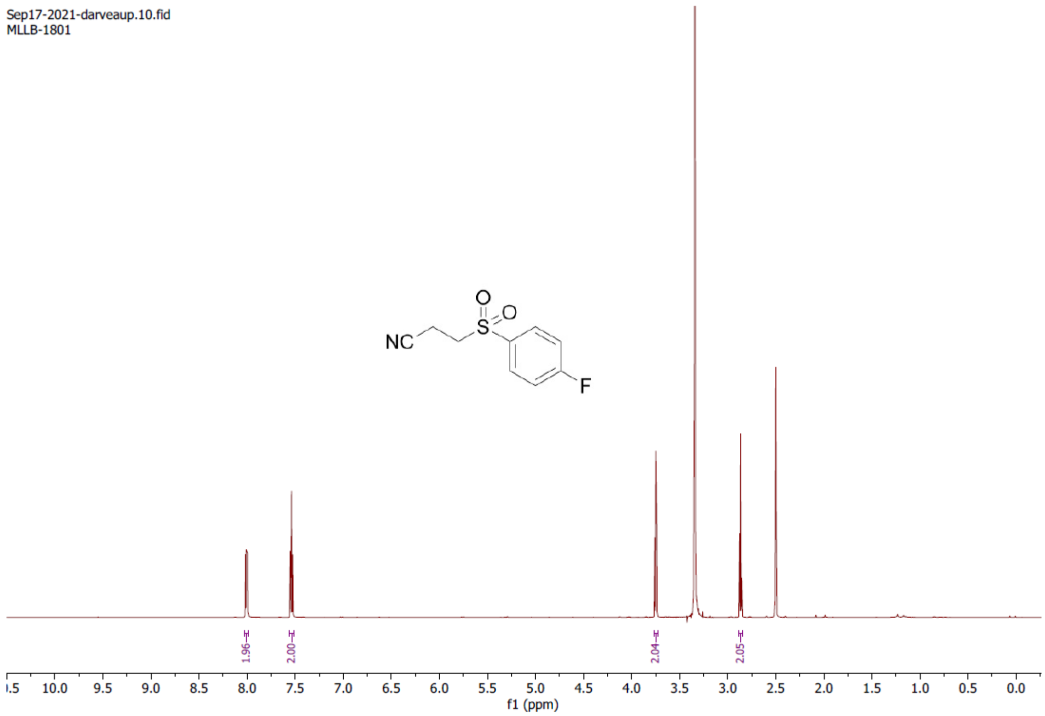


**3-((4-fluorophenyl)sulfonyl)propanenitrile (19)**

$^{13}$C NMR (176 MHz, DMSO-$d_6$)

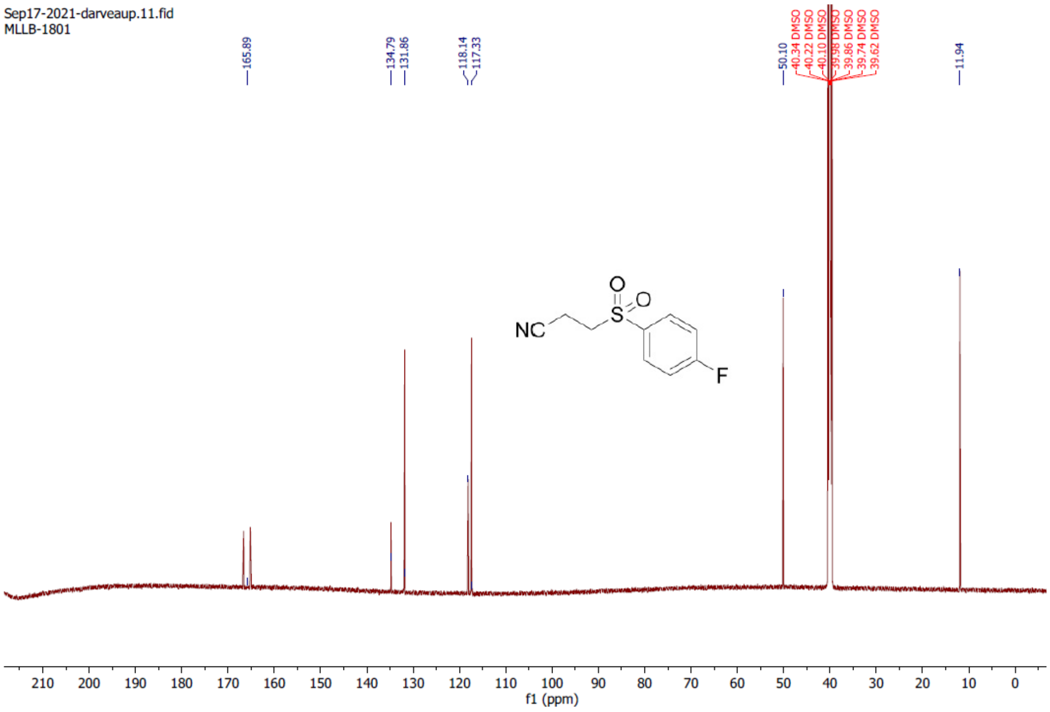

**3-(phenylsulfonyl)pyrazine-2-carbonitrile (20)**

$^{1}$H NMR (700 MHz, Chloroform-*d*)

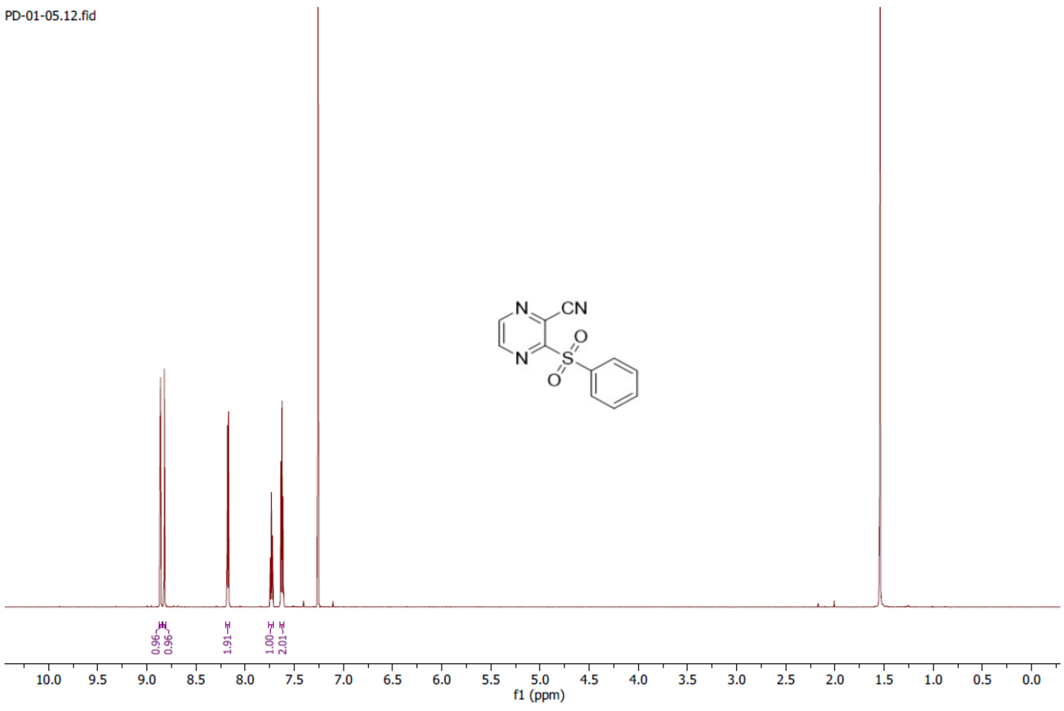


**3-(phenylsulfonyl)pyrazine-2-carbonitrile (20)**

$^{13}$C NMR (176 MHz, Chloroform-*d*)

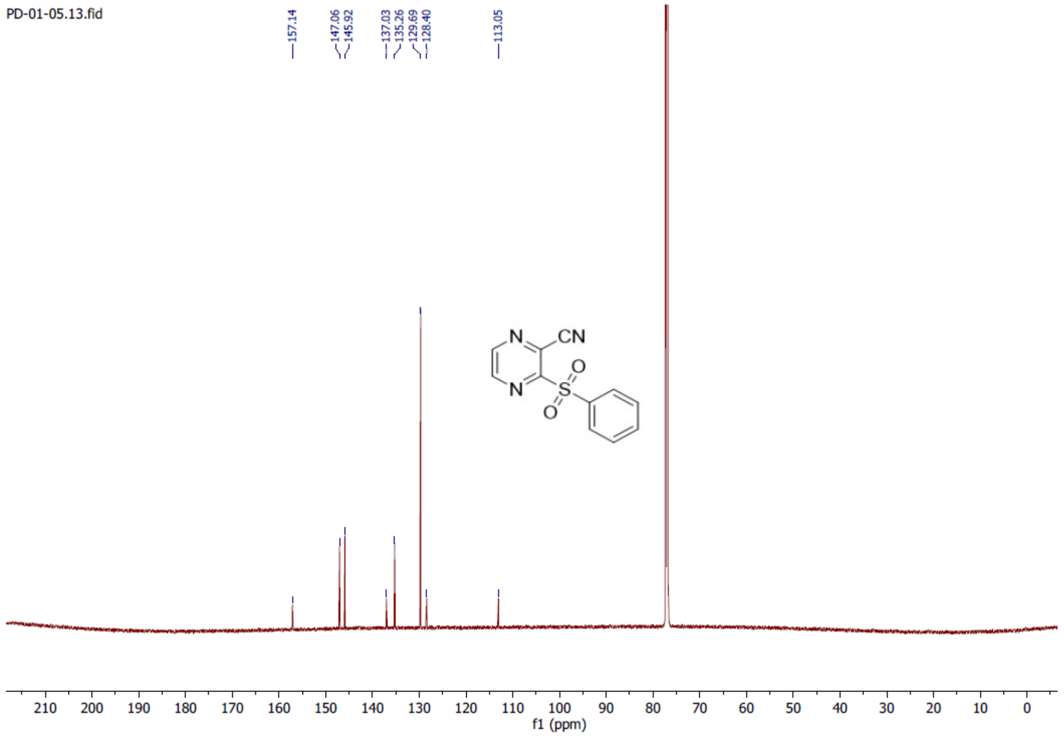

**3-((4-chlorophenyl)sulfonyl)pyrazine-2-carbonitrile (24)**

$^{1}$H NMR (700 MHz, DMSO-$d_6$)

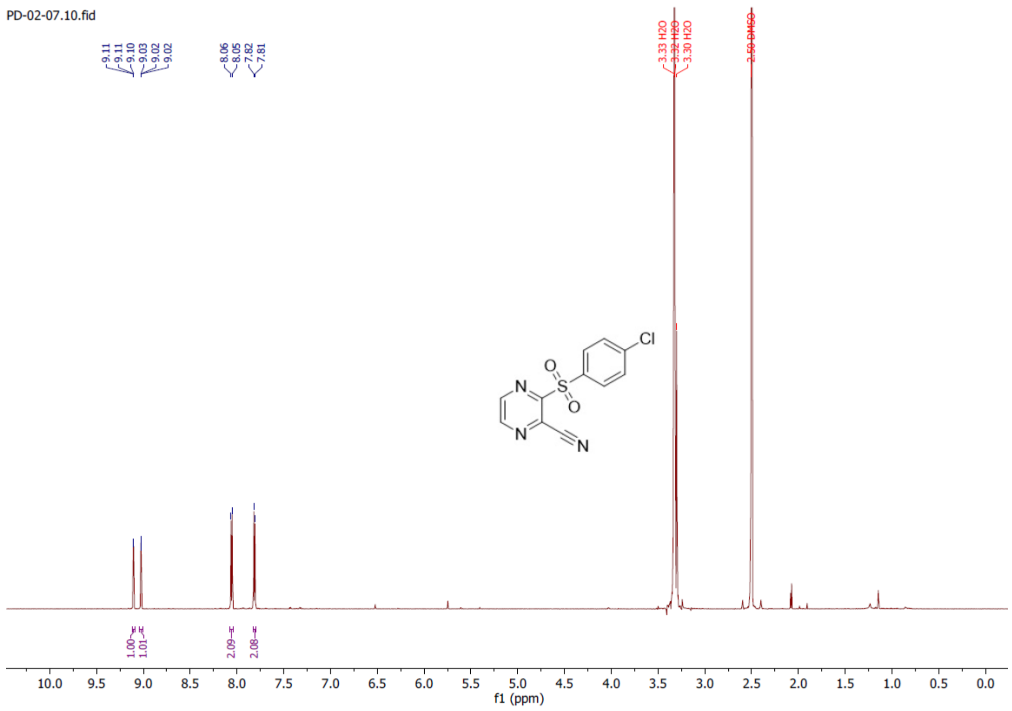


**3-((4-chlorophenyl)sulfonyl)pyrazine-2-carbonitrile (24)**

$^{13}$C NMR (176 MHz, DMSO-$d_6$)

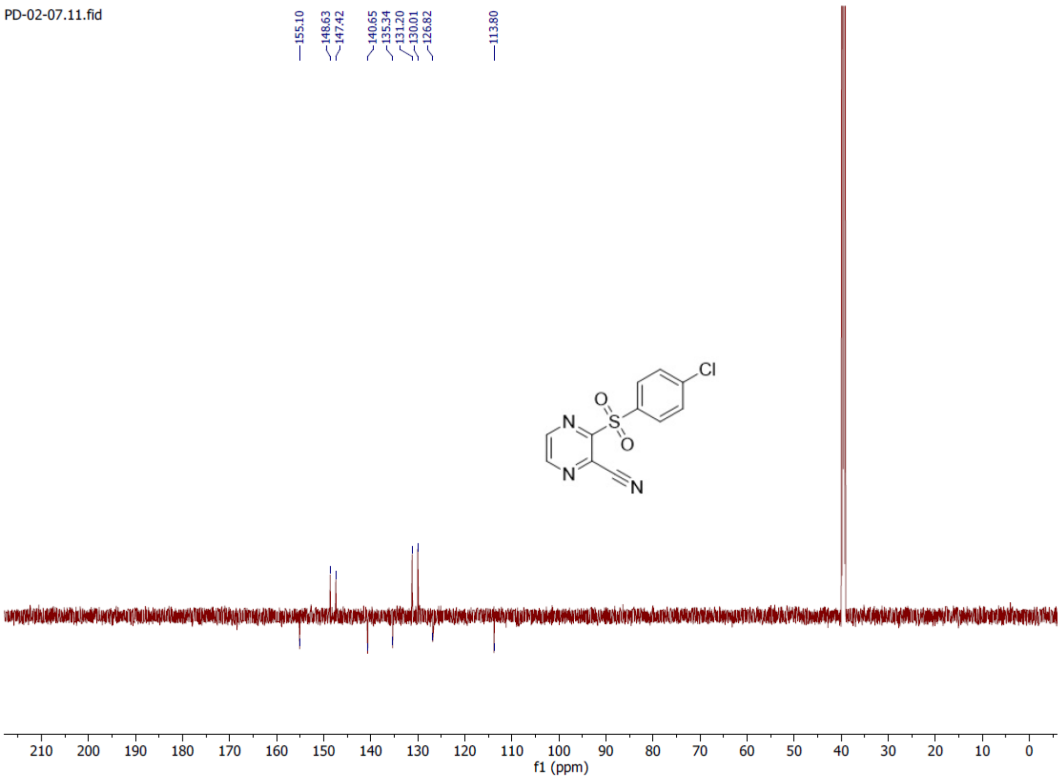

**3-((4-bromophenyl)sulfonyl)pyrazine-2-carbonitrile (25)**

$^{1}H$ NMR (700 MHz, DMSO-$d_6$)

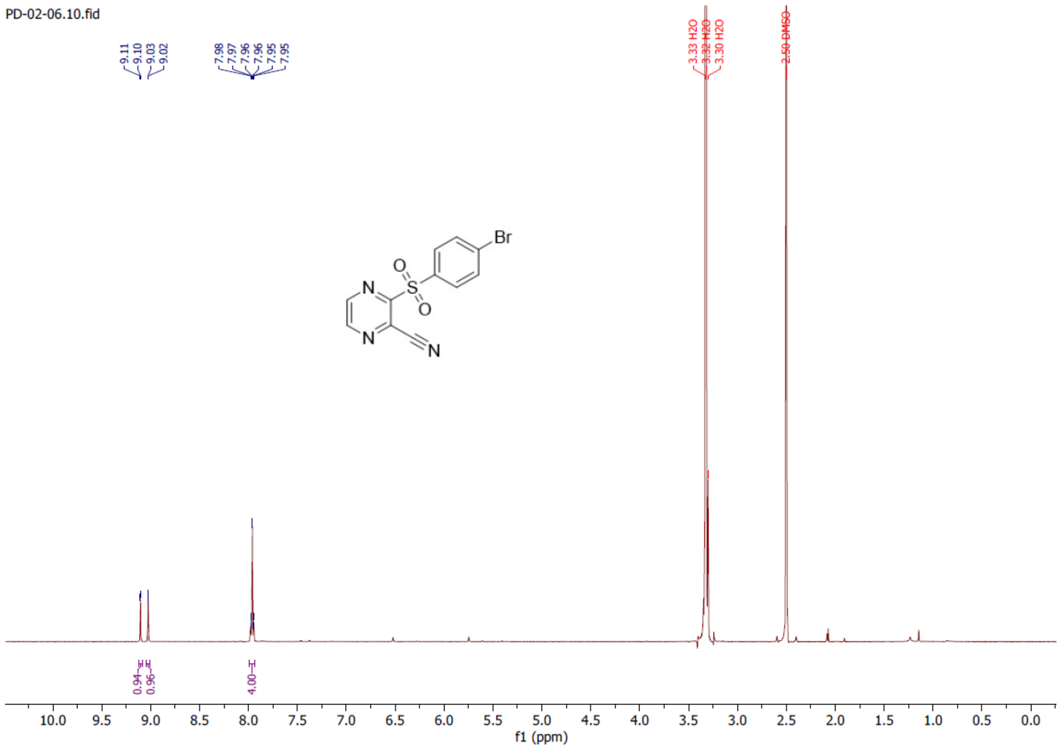


**3-((4-bromophenyl)sulfonyl)pyrazine-2-carbonitrile (25)**

$^{13}C$ NMR (176 MHz, DMSO-$d_6$)

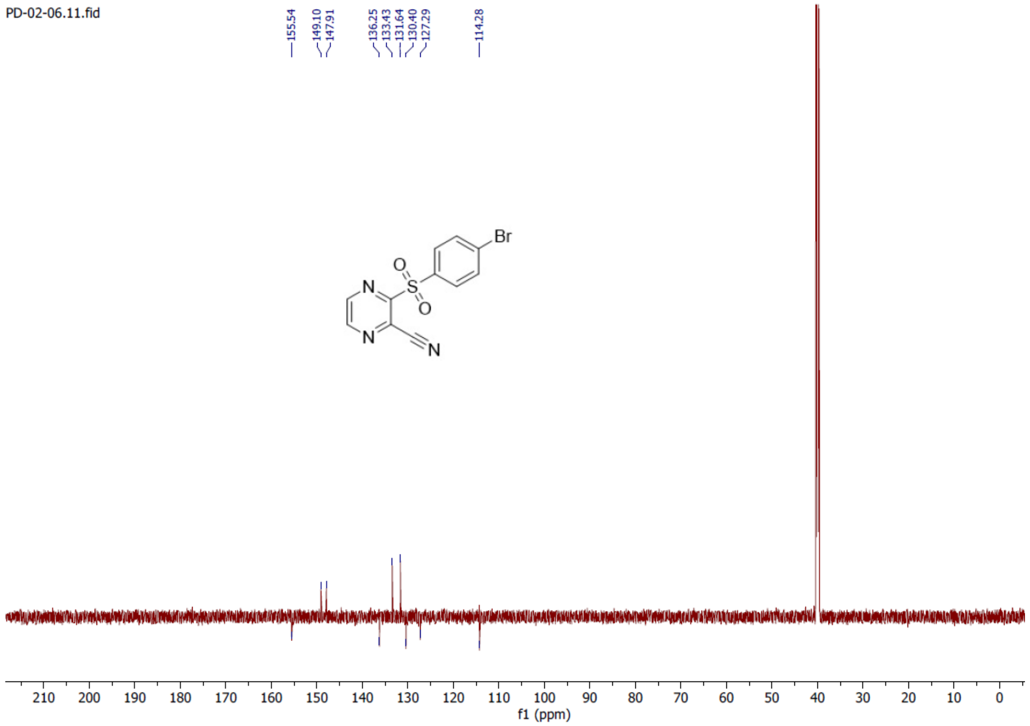

**3-((4-(trifluoromethyl)phenyl)sulfonyl)pyrazine-2-carbonitrile (26)**

$^{1}H$ NMR (700 MHz, DMSO-$d_6$)

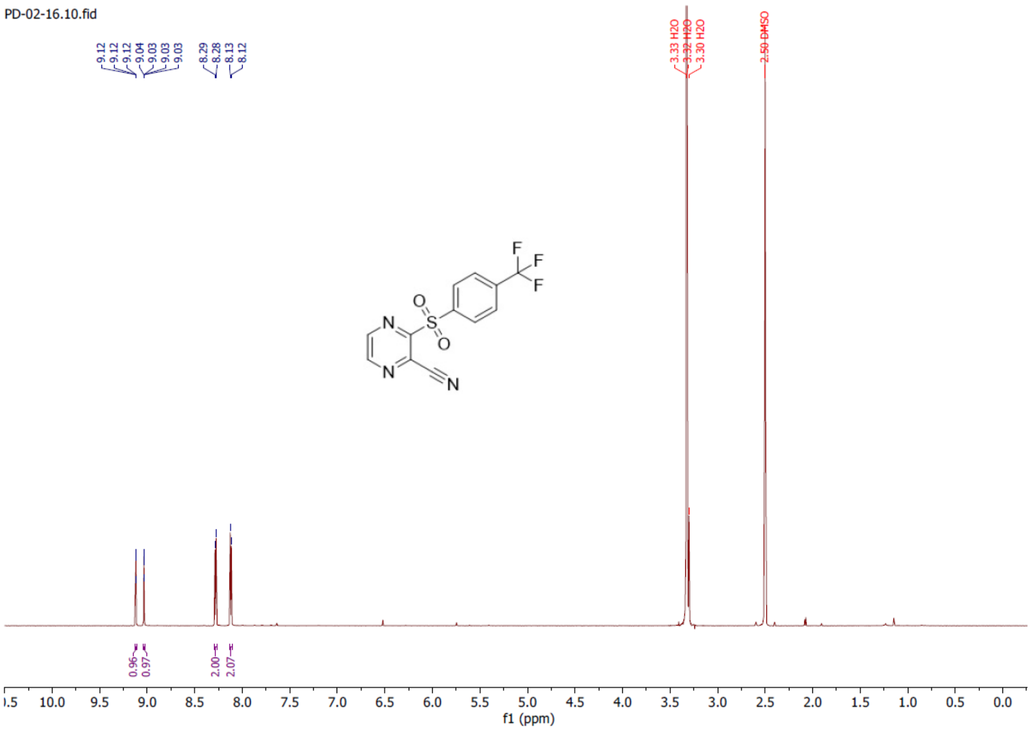


**3-((4-(trifluoromethyl)phenyl)sulfonyl)pyrazine-2-carbonitrile (26)**

$^{13}C$ NMR (176 MHz, DMSO-$d_6$)

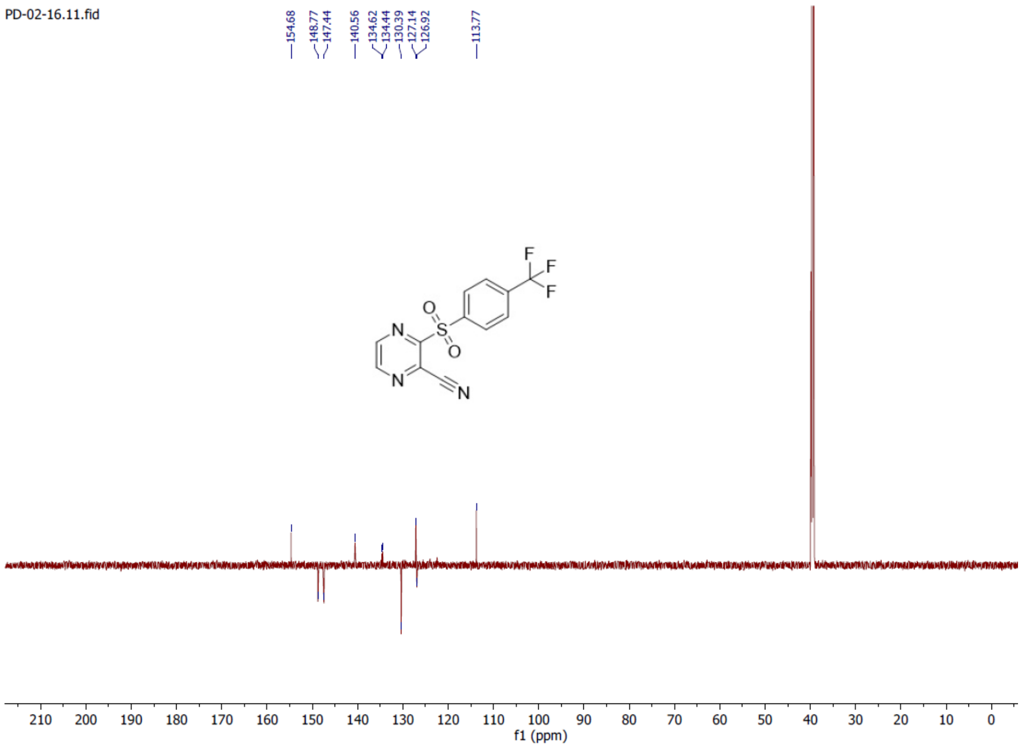

**3-((2,4-difluorophenyl)sulfonyl)pyrazine-2-carbonitrile (27)**

[1]H NMR (700 MHz, DMSO-$d_6$)

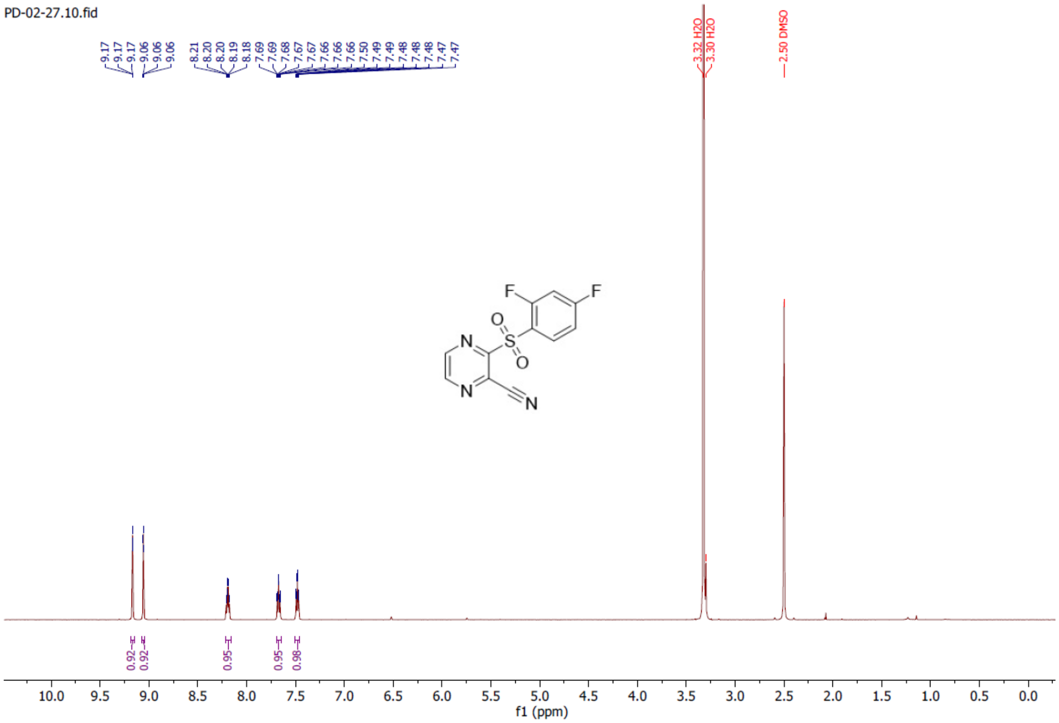


**3-((2,4-difluorophenyl)sulfonyl)pyrazine-2-carbonitrile (27)**

[13]C NMR (176 MHz, DMSO-$d_6$)

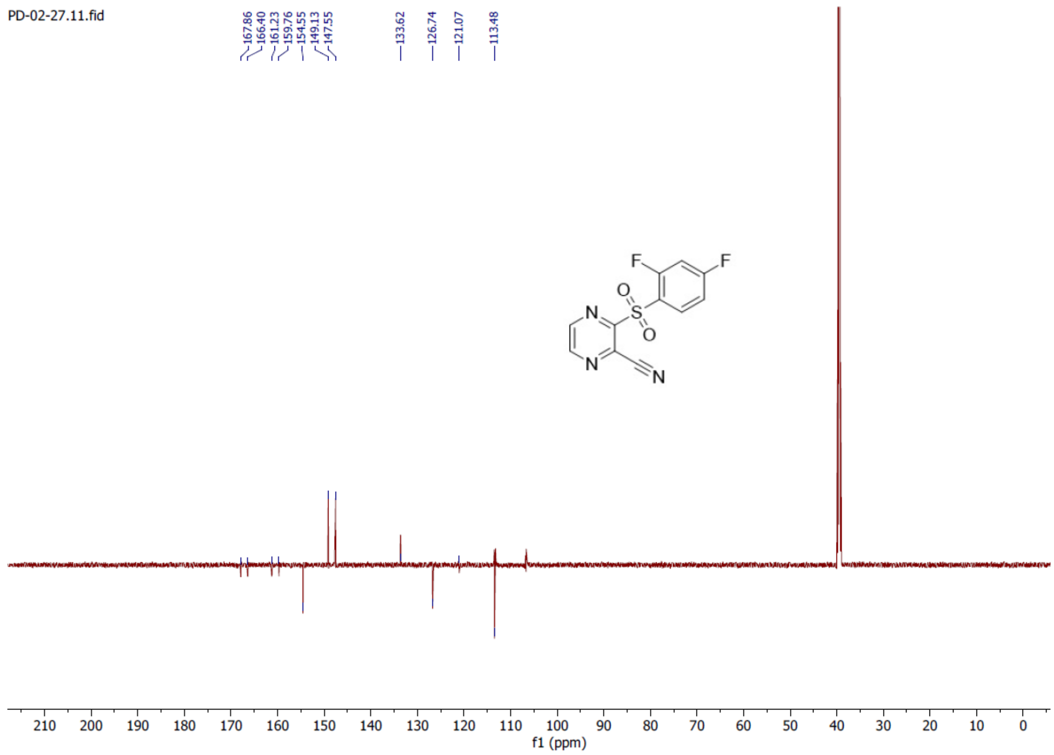

**3-((4-fluorophenyl)sulfonyl)pyrazine-2-carbonitrile (28)**

$^{1}H$ NMR (700 MHz, DMSO-$d_6$)

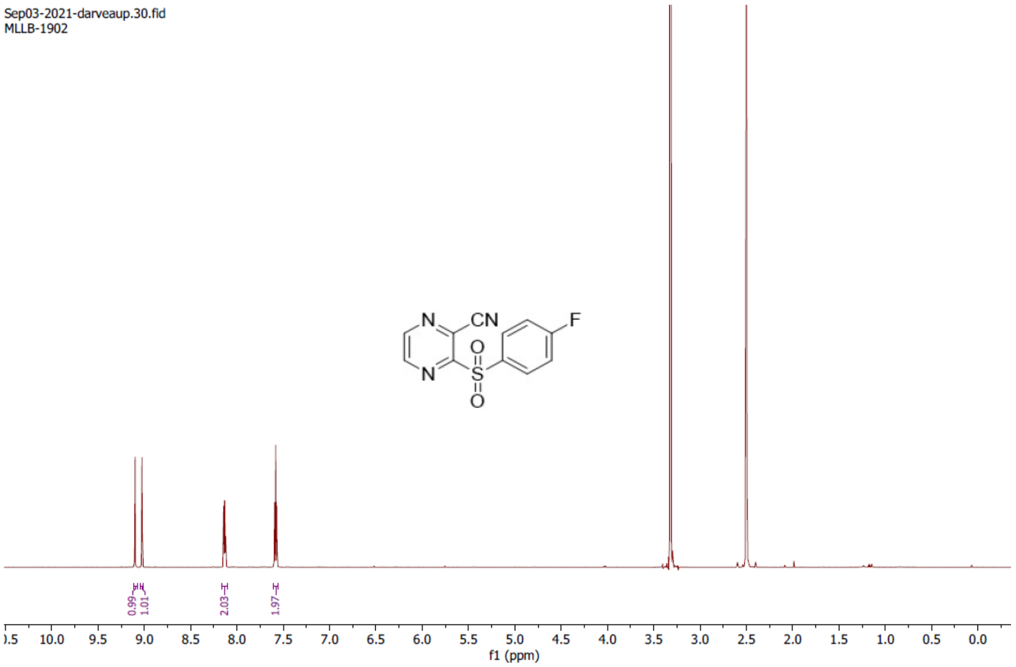


**3-((4-fluorophenyl)sulfonyl)pyrazine-2-carbonitrile (28)**

$^{13}C$ NMR (176 MHz, DMSO-$d_6$)

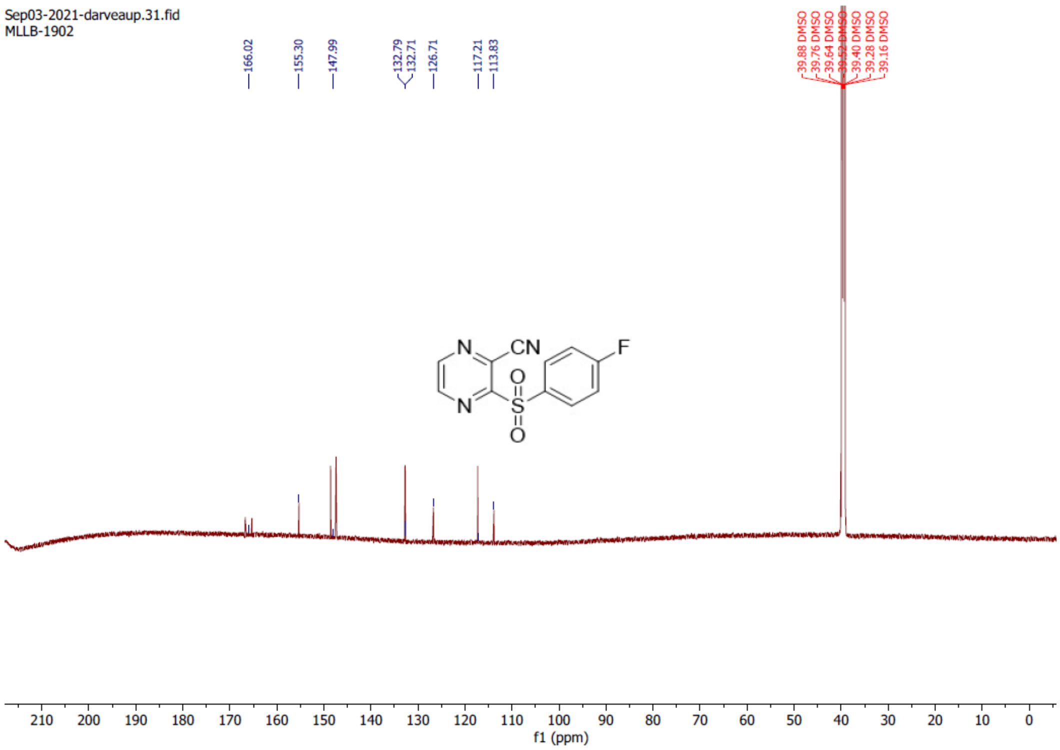

**3-((4-cyanophenyl)sulfonyl)pyrazine-2-carbonitrile (29)**

$^{1}H$ NMR (700 MHz, DMSO-$d_6$)

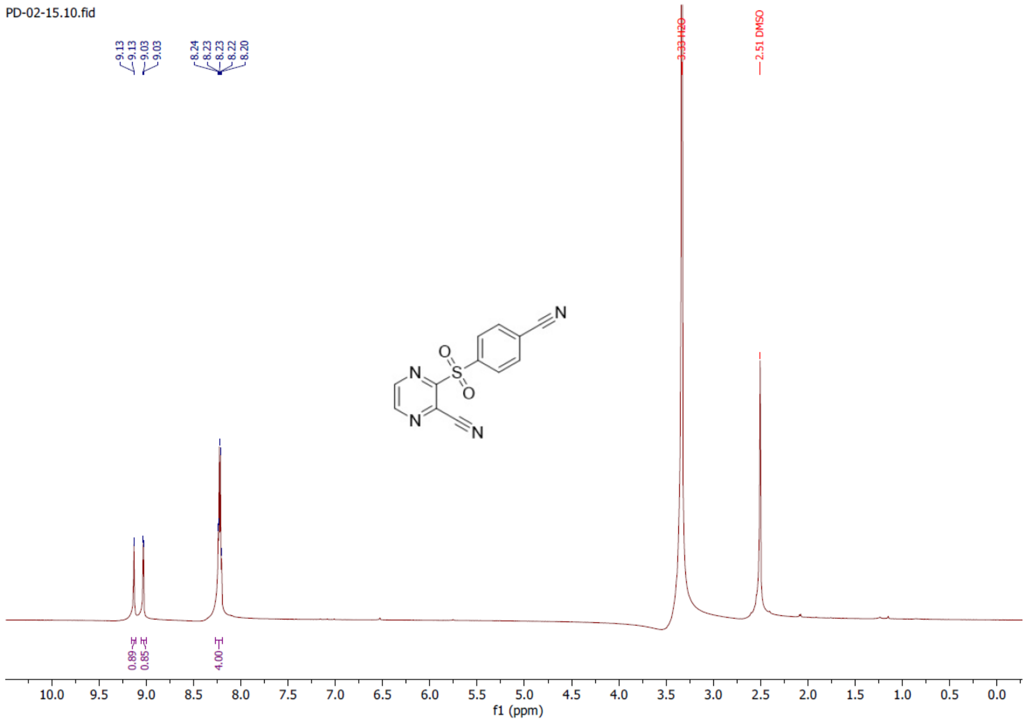


**3-((4-cyanophenyl)sulfonyl)pyrazine-2-carbonitrile (29)**

$^{13}C$ NMR (176 MHz, DMSO-$d_6$)

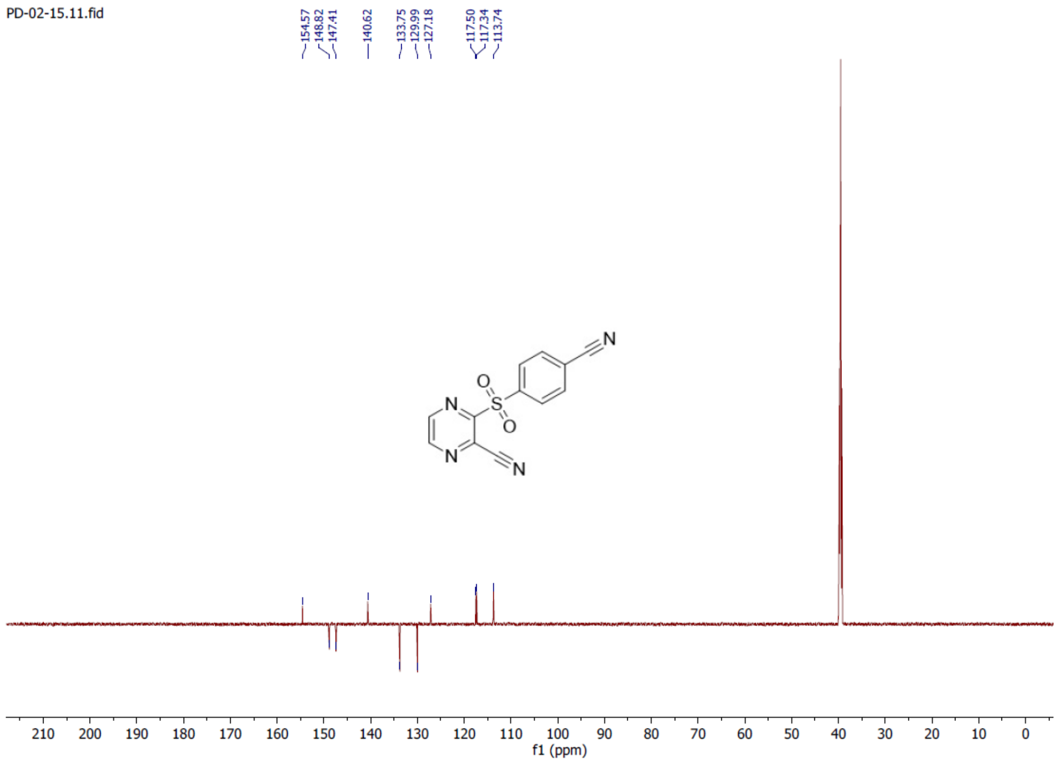

**3-((2-fluorophenyl)sulfonyl)pyrazine-2-carbonitrile (30)**

$^1$H NMR (700 MHz, DMSO-$d_6$)

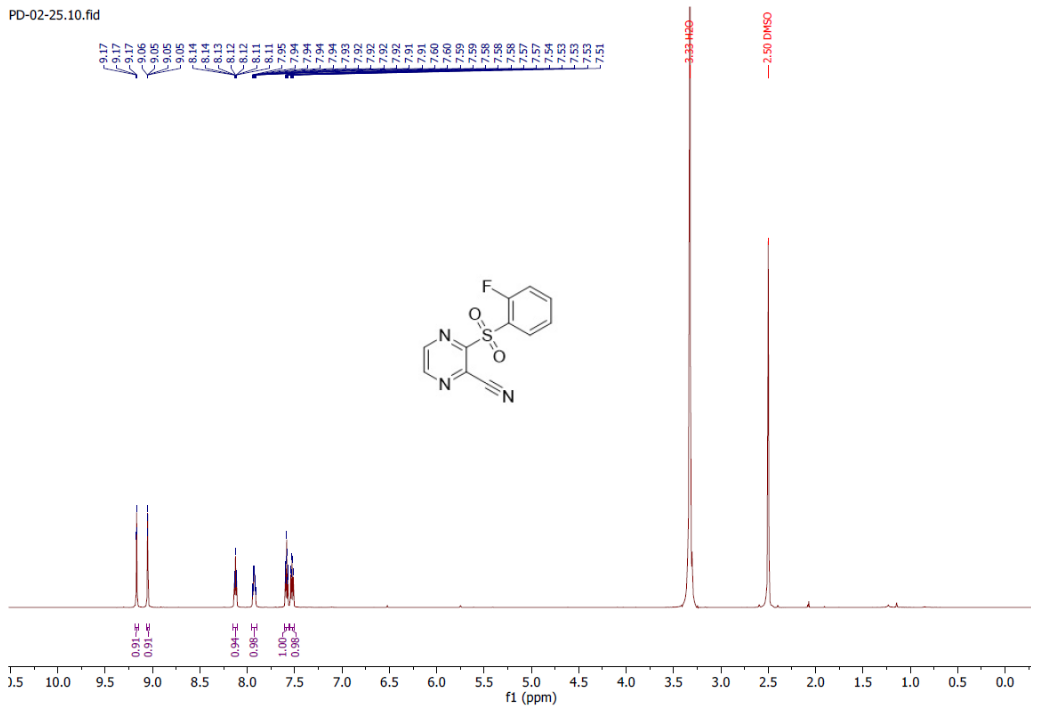


**3-((2-fluorophenyl)sulfonyl)pyrazine-2-carbonitrile (30)**

$^{13}$C NMR (176 MHz, DMSO-$d_6$)

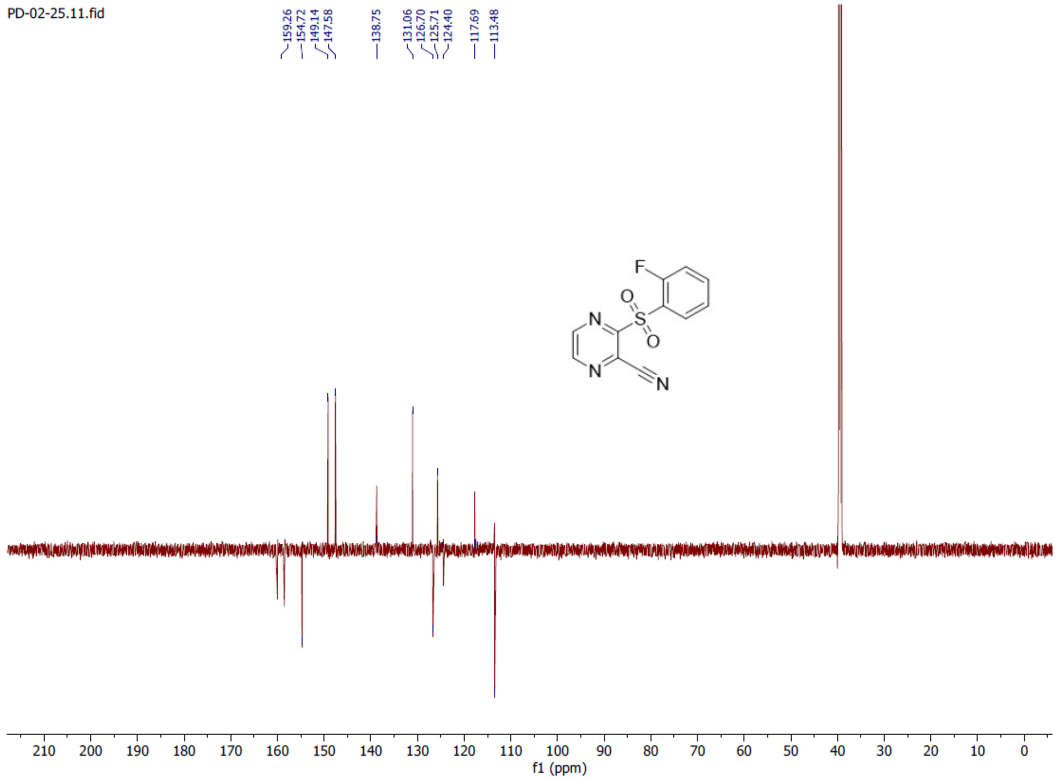

**3-((3,4-difluorophenyl)sulfonyl)pyrazine-2-carbonitrile (31)**

$^{1}H$ NMR (700 MHz, DMSO-$d_6$)

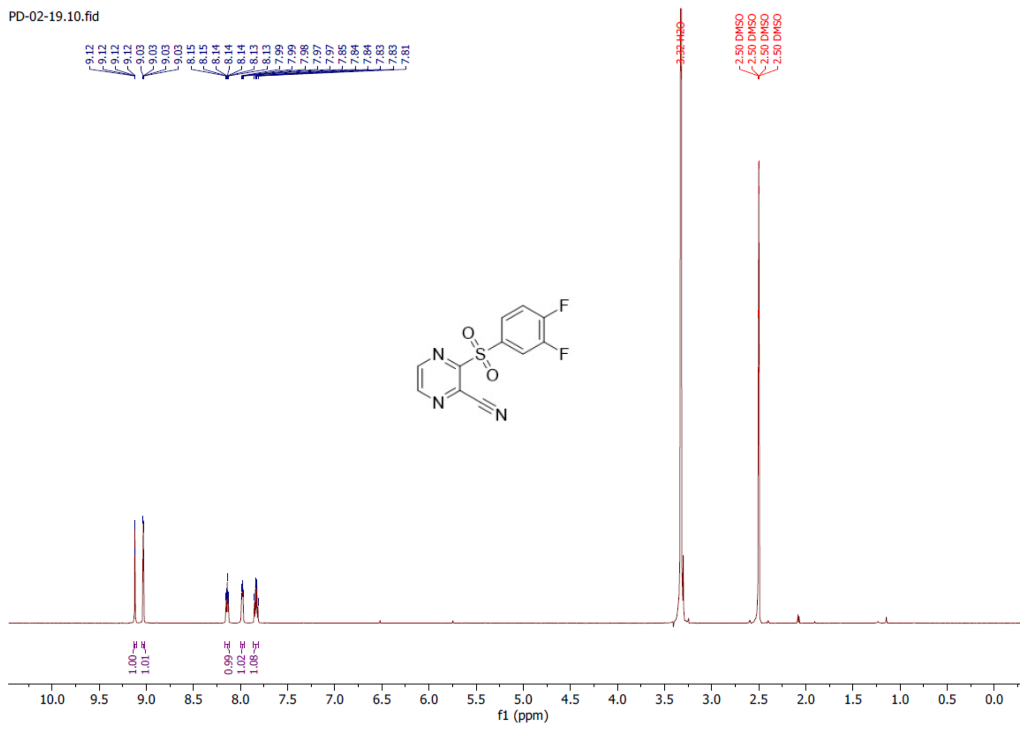


**3-((3,4-difluorophenyl)sulfonyl)pyrazine-2-carbonitrile (31)**

$^{13}C$ NMR (176 MHz, DMSO-$d_6$)

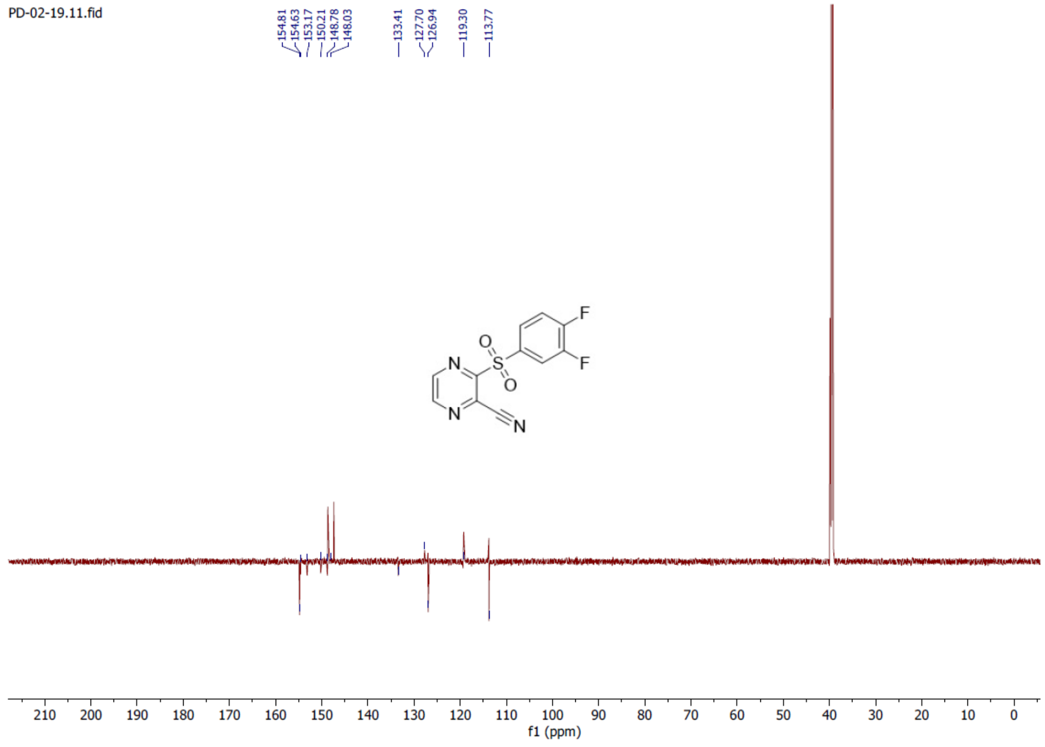

**3-((3,5-difluorophenyl)sulfonyl)pyrazine-2-carbonitrile (32)**

$^{1}$H NMR (700 MHz, DMSO-$d_6$)

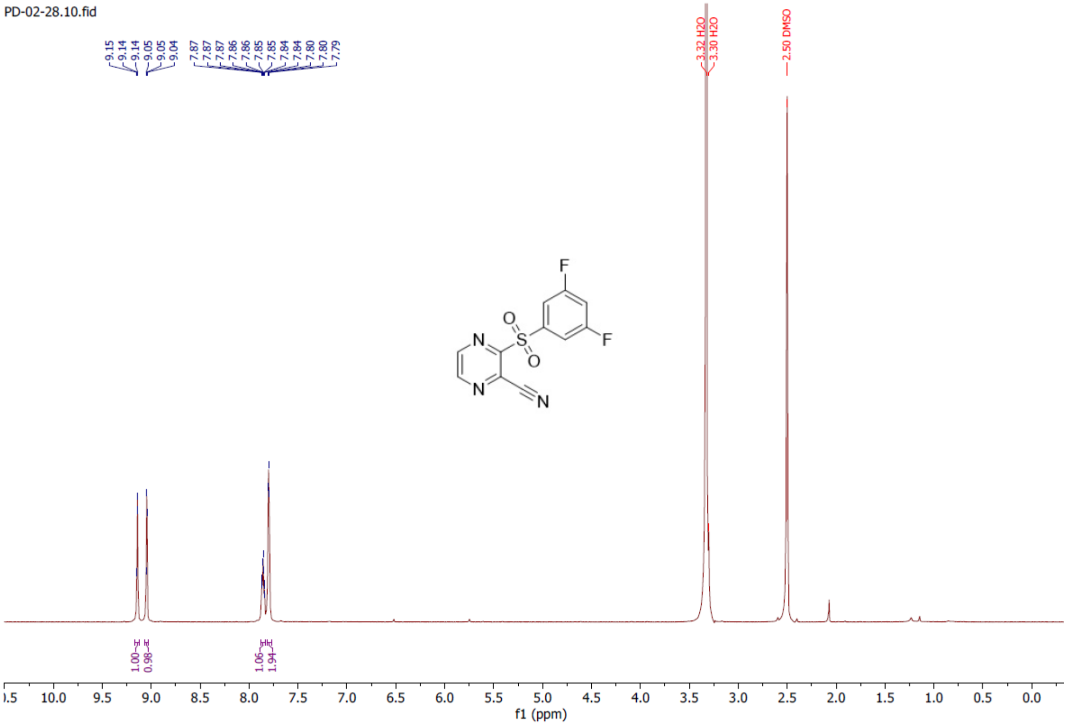


**3-((3,5-difluorophenyl)sulfonyl)pyrazine-2-carbonitrile (32)**

$^{13}$C NMR (176 MHz, DMSO-$d_6$)

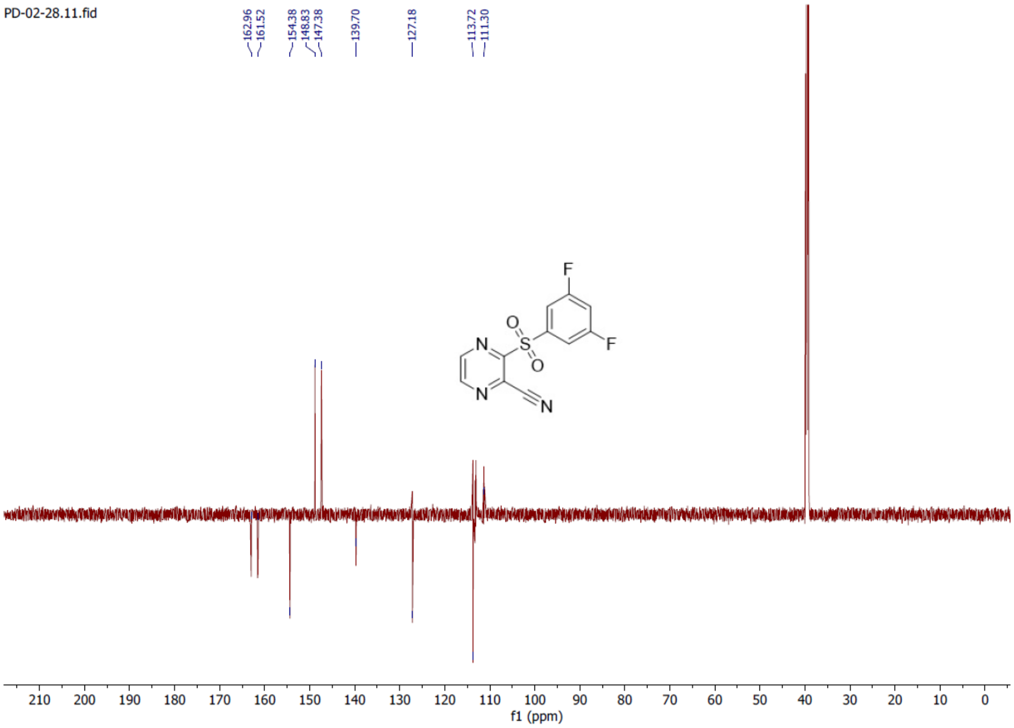

**2-((4-fluorophenyl)sulfonyl)-6-methylnicotinonitrile (33)**

$^{1}H$ NMR (700 MHz, Chloroform-*d*)

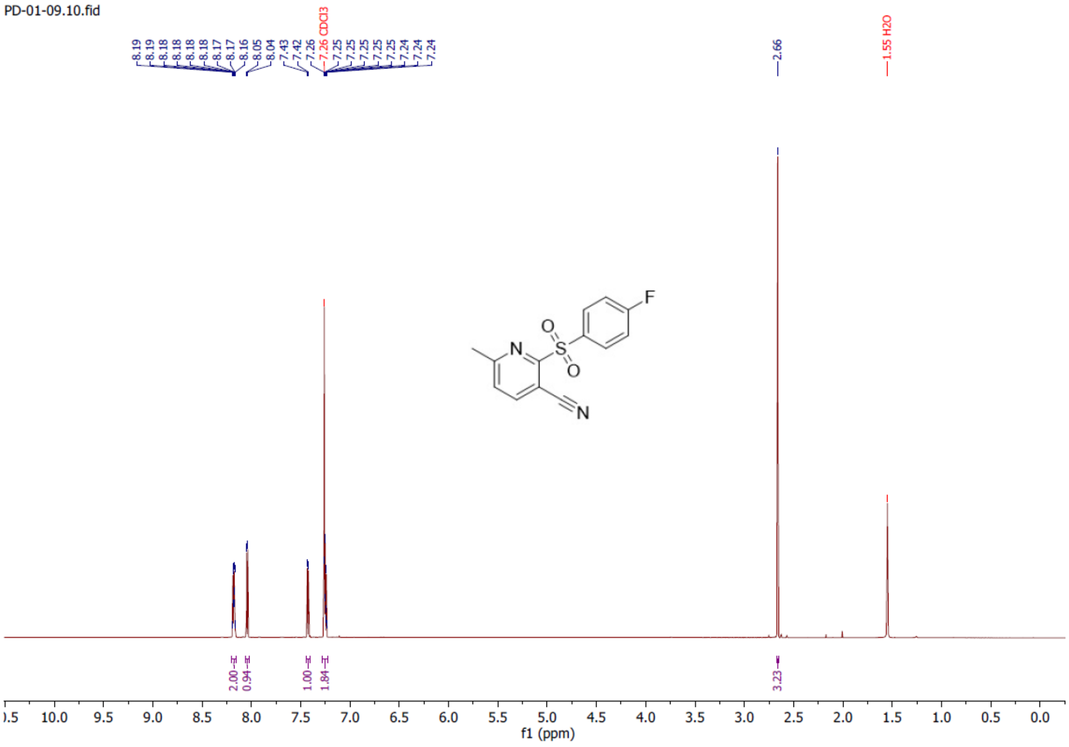


**2-((4-fluorophenyl)sulfonyl)-6-methylnicotinonitrile (33)**

$^{13}C$ NMR (176 MHz, Chloroform-*d*)

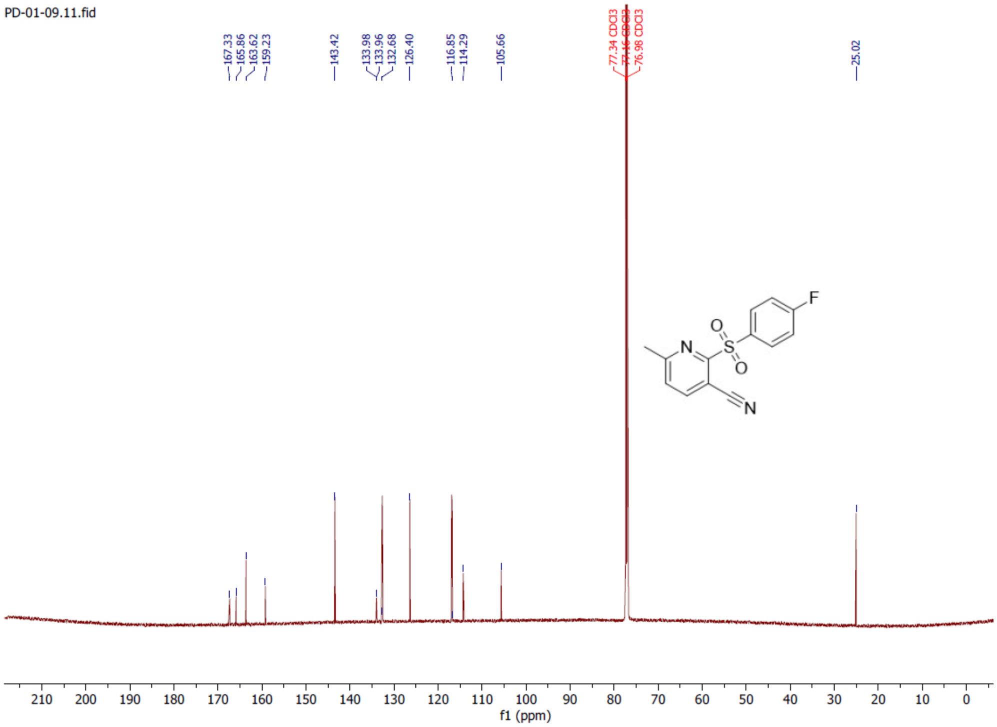

**2-((4-fluorophenyl)sulfonyl)nicotinonitrile (34)**

$^{1}H$ NMR (700 MHz, DMSO-$d_6$)

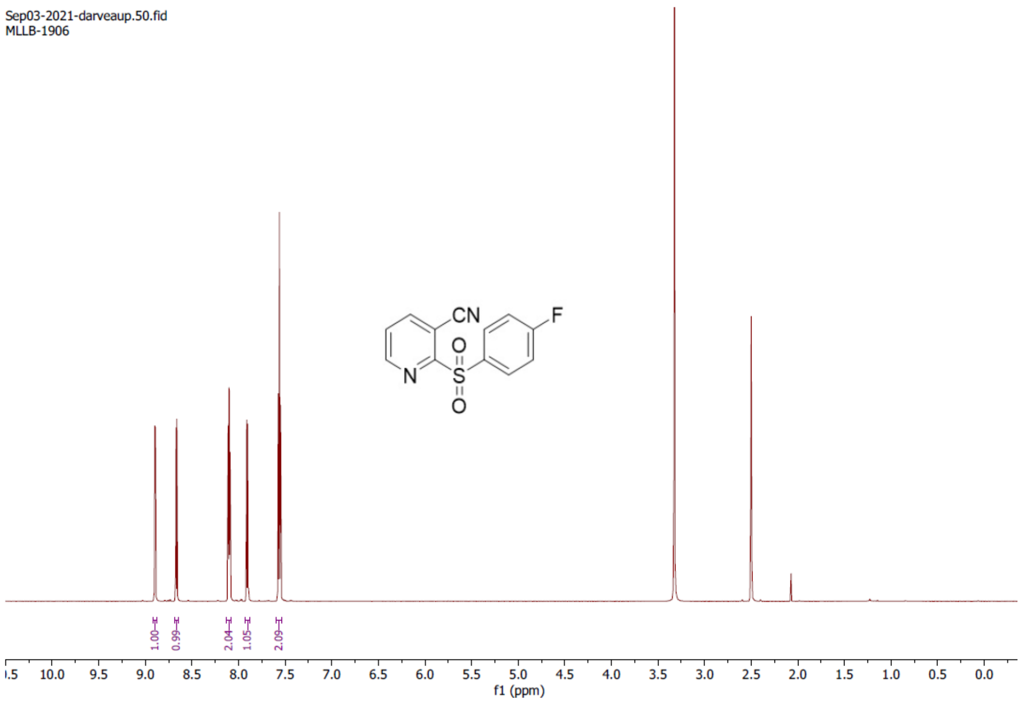


**2-((4-fluorophenyl)sulfonyl)nicotinonitrile (34)**

$^{13}C$ NMR (176 MHz, DMSO-$d_6$)

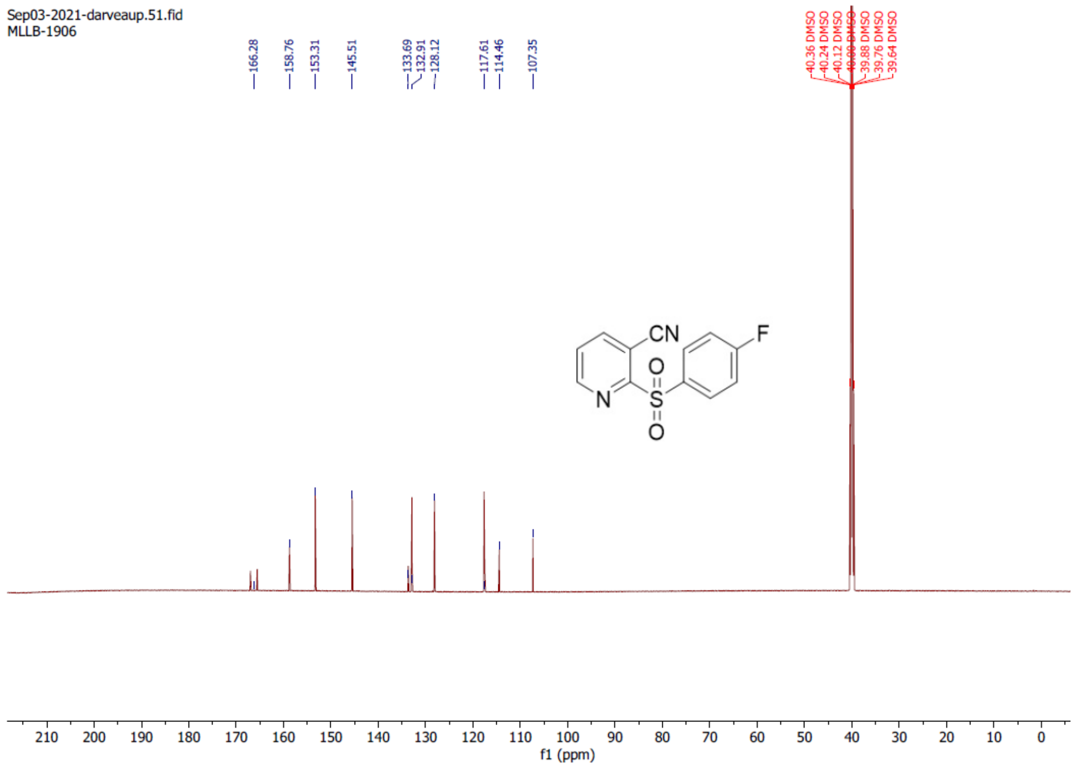
